# Detection of cross-contamination and strong mitonuclear discordance in two species groups of sawfly genus Empria (Hymenoptera, Tenthredinidae)

**DOI:** 10.1101/525626

**Authors:** Marko Prous, Kyung Min Lee, Marko Mutanen

## Abstract

In several sawfly taxa strong mitonuclear discordance has been observed, with nuclear genes supporting species assignments based on morphology, whereas the barcode region of the mitochondrial COI gene suggesting different relationships. As previous studies were based on only few nuclear genes, the causes and the degree of mitonuclear discordance remain ambiguous. Here, we obtain genomic-scale ddRAD data together with Sanger sequencing of mitochondrial COI and two to three nuclear protein coding genes to investigate species limits and mitonuclear discordance in two closely related species groups within the sawfly genus *Empria*. As found previously based on nuclear ITS and mitochondrial COI sequences, species are in most cases supported as monophyletic based on previous and new nuclear data reported here, but not based on mitochondrial COI. This mitonuclear discordance can be explained by occasional mitochondrial introgression with little or no nuclear gene flow, a pattern that might be common in haplodiploid taxa with slowly evolving mitochondrial genomes. Some species in *E. immersa* group are not recovered as monophyletic also based on nuclear data, but this could partly be because of unresolved taxonomy. Preliminary analyses of ddRAD data did not recover monophyly of *E. japonica* within *E. longicornis* group (three Sanger sequenced nuclear genes strongly supported monophyly), but closer examination of the data and additional Sanger sequencing suggested that both specimens were substantially (possibly 10–20% of recovered loci) cross-contaminated. A reason could be due to specimen identification tag jumps during sequencing library preparation of pooled specimens that in previous studies have been shown to affect up to 2.5% of the sequenced reads. We provide an R script to examine patterns of identical loci among the specimens and estimate that cross-contamination rate is not unusually high for our ddRAD dataset as a whole (based on counting identical sequences between *immersa* and *longicornis* groups that are well separated from each other and probably do not hybridise). The high rate of cross-contamination for both *E. japonica* specimens might be explained by small number of recovered loci (~1000) compared to most other specimens (>10 000 in some cases) because of poor sequencing results. We caution drawing unexpected biological conclusions when closely related specimens are pooled before sequencing and tagged only at one end of the molecule or at both ends using unique combination of limited number of tags (less than the number of specimens).

## 1. Introduction

Continuing developments in high-throughput sequencing technologies and falling of prices makes it increasingly easier to collect genome-scale data for many non-model organisms. The large amount of data that could be obtained with high-throughput next generation sequencing methods makes it possible to answer many biological questions simultaneously (in phylogeny, population genetics, evolutionary ecology etc.) and in higher resolution than would be possible with more traditional methods (e.g. Sanger sequencing of one or few markers, genotyping by microsatellites etc.). However, the large amount of data that is generated with next generation sequencing methods introduces its own problems that are hardly relevant when only few markers are analysed. Genome-scale or phylogenomic datasets are plagued mainly by two types or errors: data errors and systematic errors (Philippe et al., 2017). Data errors, such as assembly and alignment artefacts or contaminants for example, are easy to control for in single-gene scale datasets, but prohibitive in genome-scale datasets if done manually. Because automated methods of dataset assembly are not (yet) perfect, some data errors are nearly always introduced in phylogenomic datasets. Even if the dataset is perfectly assembled (all contaminants and non-homologous alignments excluded), one still has to consider systematic errors (e.g. biases in nucleotide or amino acid composition, unequal rates of evolution) which only increase with dataset size and therefore could seriously mislead phylogenomic analyses, although this problem becomes critical only when dealing with ancient divergences (tens and more millions of years ago) (Philippe et al., 2017, 2005; Tarver et al., 2016).

DNA barcoding of single molecular marker for the purpose of species identification can also benefit from high-throughput sequencing, as hundreds or thousands individuals could be sequenced simultaneously (e.g. Cruaud et al., 2017; Hebert et al., 2018; Meier et al., 2016). For animals, a ~650 bp fragment from 5’ end of the mitochondrial cytochrome c oxidase I (COI) has been chosen as the standard barcoding marker (Hebert et al., 2003), which by now has been sequenced from more than five million individuals according to Barcode of Life (BOLD) database (http://www.boldsystems.org). Although this short mitochondrial fragment seems to be suitable in most species rich groups, such as Coleoptera and Lepidoptera (Mutanen et al., 2016; Pentinsaari et al., 2017; Zahiri et al., 2017), rampant mitochondrial introgression is also known in some groups (Sloan et al., 2017). In some cases the usefulness of COI sequences is not clear due to lack of sequencing efforts and / or taxonomic research. For example, while large-scale COI sequencing efforts have been applied also to many hyperdiverse insect groups, congruence with sufficiently informative nuclear genes and / or morpho-taxonomy has not always been evaluated (Alex Smith et al., 2013; Hebert et al., 2016). Nevertheless, some theoretical considerations can give indications in which cases mitonuclear discordances could be expected at increased rate (Ivanov et al., 2018; Sloan et al., 2017). Particularly, Patten et al. (2015) found recently through theoretical modelling that haplodiploid species may be especially prone to biased mitochondrial introgression, which could be amplified by several other adaptive and non-adaptive conditions (reviewed by Sloan et al., 2017). The most species rich group (at least in terms of described species) of haplodiploid animals is Hymenoptera (sawflies, ants, bees, and wasps) and could therefore be a good candidate for investigating mitochondrial introgression and utility of mitochondrial barcodes. Besides haplodiploidy, mutation rate of mitochondrial DNA (mtDNA) could also be a factor affecting rate of mitochondrial introgression. Sloan et al. (2017) suggested that lower mutation rates promote adaptive mitochondrial introgression while higher rates lead more likely to compensatory co-evolution and mitonuclear incompatibilities. As mitochondrial genomes of Apocrita (the bulk of hymenopteran species) evolve faster than those of basal hymenopterans (Kaltenpoth et al., 2012; Ma et al., 2019; Niu et al., 2019; Tang et al., 2019), mitochondrial introgression might be less common in Apocrita compared to sawflies. Within the sawflies, Xyeloidea, Pamphilioidea, and Tenthredinoidea have the slowest evolving mtDNA, while Cephoidea, Orussoidea, Siricoidea, and possibly Anaxyleoidea (which are more closely related to Apocrita), have intermediate or fast evolutionary rate (Ma et al., 2019; Niu et al., 2019; Tang et al., 2019). While we are not aware of cases of large-scale discordance between mitochondrial barcodes and species boundaries in Apocrita, there are several such cases among sawflies, particularly among Tenthredinoidea (Linnen and Farrell, 2007; Schmidt et al., 2017). However, in all those cases discordance were identified based on morphology and COI barcodes or morphology plus few nuclear genes and COI barcodes, and it is likely that in some cases operational factors, such as over-splitting of species, are involved too (cf. Mutanen et al., 2016).

Here we investigate based on genome-scale data the phylogeny and species limits in two closely related species groups (divergence probably not more than few million years) within the sawfly genus *Empria* Lepeletier & Serville, 1828 (Hymenoptera, Tenthredinidae). The genus includes at least 60 species, several of which are still undescribed (Prous, 2012). Most species in the genus are externally rather similar to each other, which makes species identification difficult. However, the differences in the structure of ovipositors and penis valves are often very clear even between closely related species (Prous, 2012).

Taxonomic and limited phylogenetic studies on the genus have revealed two species complexes (*longicornis* and *immersa* groups) where species delimitation has been especially problematic (Prous, 2012; Prous et al., 2014, 2011b). Based on morphology, the main evidence indicating the presence of more than one species in both of the groups, is the structure of the female ovipositor, which often shows clear differences between species and which correlates with host plant use (Prous, 2012; Prous et al., 2011b). Species in *longicornis* group specialise on different herbaceous genera in Rosaceae (specifically in subfamily Rosoideae and genus *Dryas*) and species in *immersa* group on *Betula* or *Salix*. Differences in other morphological characters (including male genitalia) are rather weak between the species, but can nevertheless be helpful in species identification. While sequencing of mitochondrial COI gene did not reveal any correlation with species boundaries defined based on morphological and ecological data, nuclear ITS (internal transcribed spacers 1 and 2) sequence data did (Prous, 2012; Prous et al., 2011b). Discord between mitochondrial and morphological plus nuclear ITS data in these groups is quite remarkable: different species frequently have identical COI barcodes (658 bp) or even complete (1536 bp) COI gene (Prous et al., 2011b), while at the same time different specimens of the same species can diverge by 3.3% in the barcoding region. To better understand this discord and to test species boundaries in *longicornis* and *immersa* groups, we collected genome-wide data using double digest RADseq (Lee et al., 2018; Peterson et al., 2012) and sequenced long fragments of two to three nuclear protein coding genes.

Results showed that in some cases there were substantial amount of cross-contamination in RADseq data, which might have escaped detection without the knowledge of the organisms involved (based on morphological, ecological and single gene data) and initial manual checks. This cross-contamination had significant impact on the phylogenetic tree building and population admixture analyses. We developed a workflow to detect possible cases of cross-contamination that could be excluded from downstream analyses.

## 2. Materials and methods

### 2.1 DNA extraction

For most specimens, DNA had been extracted as described in Prous et al. (2011b). New DNA extractions for this study were obtained with an EZNA Tissue DNA Kit (Omega Bio-tek) according to the manufacturer’s protocol and stored at −20 °C for later use. Typically, the middle right leg was used for DNA extraction, but for males the whole genital capsule was often additionally used to increase DNA yield and to free penis valves from muscles for photography. Specimens that were selected for sequencing are listed in Table 1.

**Table 1.**
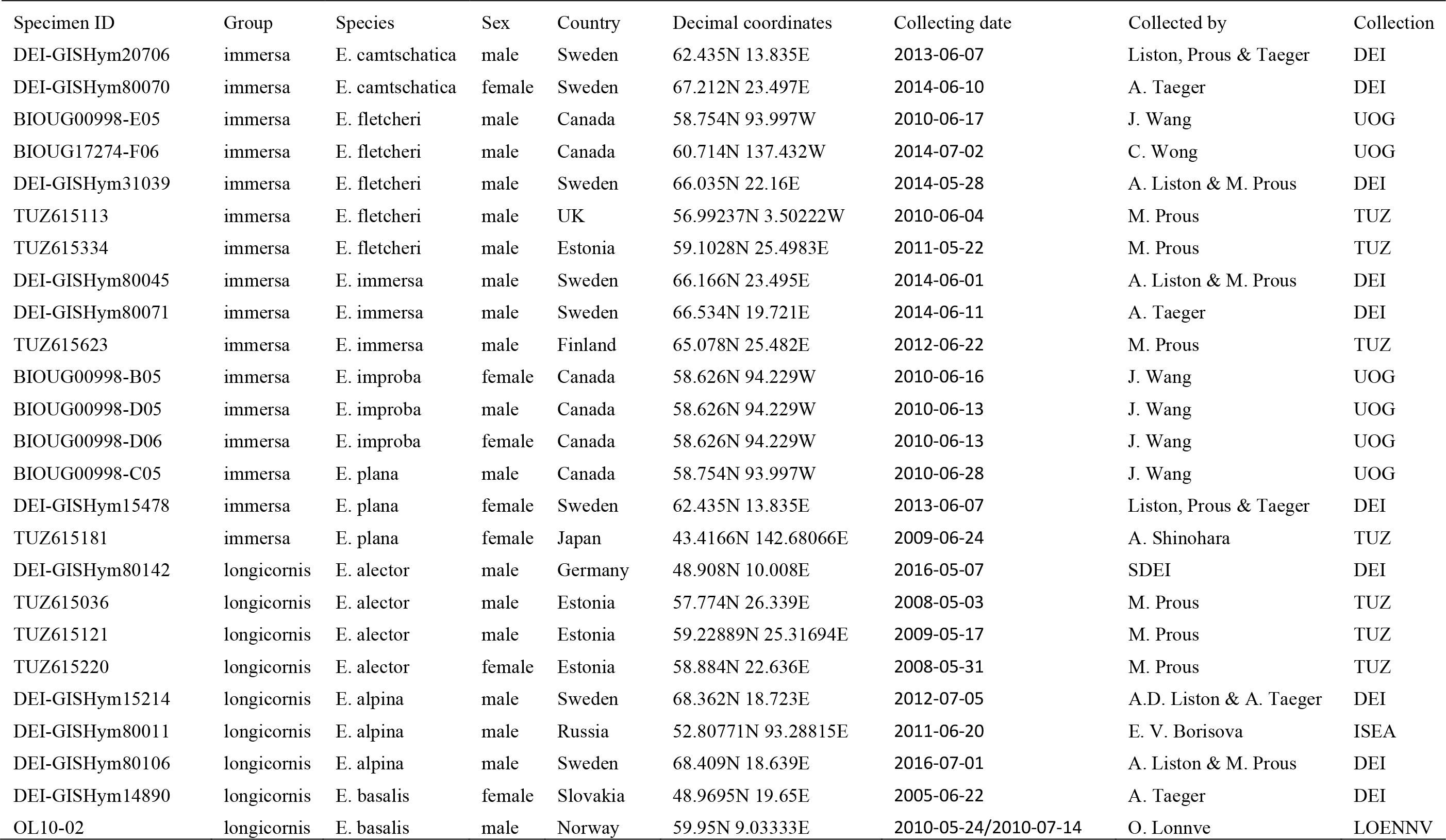
Collecting data of *Empria* specimens selected for sequencing. CMH – Private collection of Mikk Heidemaa (Tartu, Estonia); DEI-Senckenberg Deutsches Entomologisches Institut, Müncheberg, Germany; ISEA - Institute of Systematics and Ecology of Animals, Russian Academy of Sciences, Novosibirsk, Russia; IZBE - Estonian University of Life Sciences, Tartu, Estonia; LOENNV - Private collection Ole Lønnve, Oslo, Norway; TUZ - University of Tartu, Tartu, Estonia; UOG - University of Guelph, Guelph, Canada; USNM - Smithsonian Institution, National Museum of Natural History, Washington DC, USA; ZIN - Zoological Institute, Russian Academy of Sciences, Saint Petersburg, Russia.

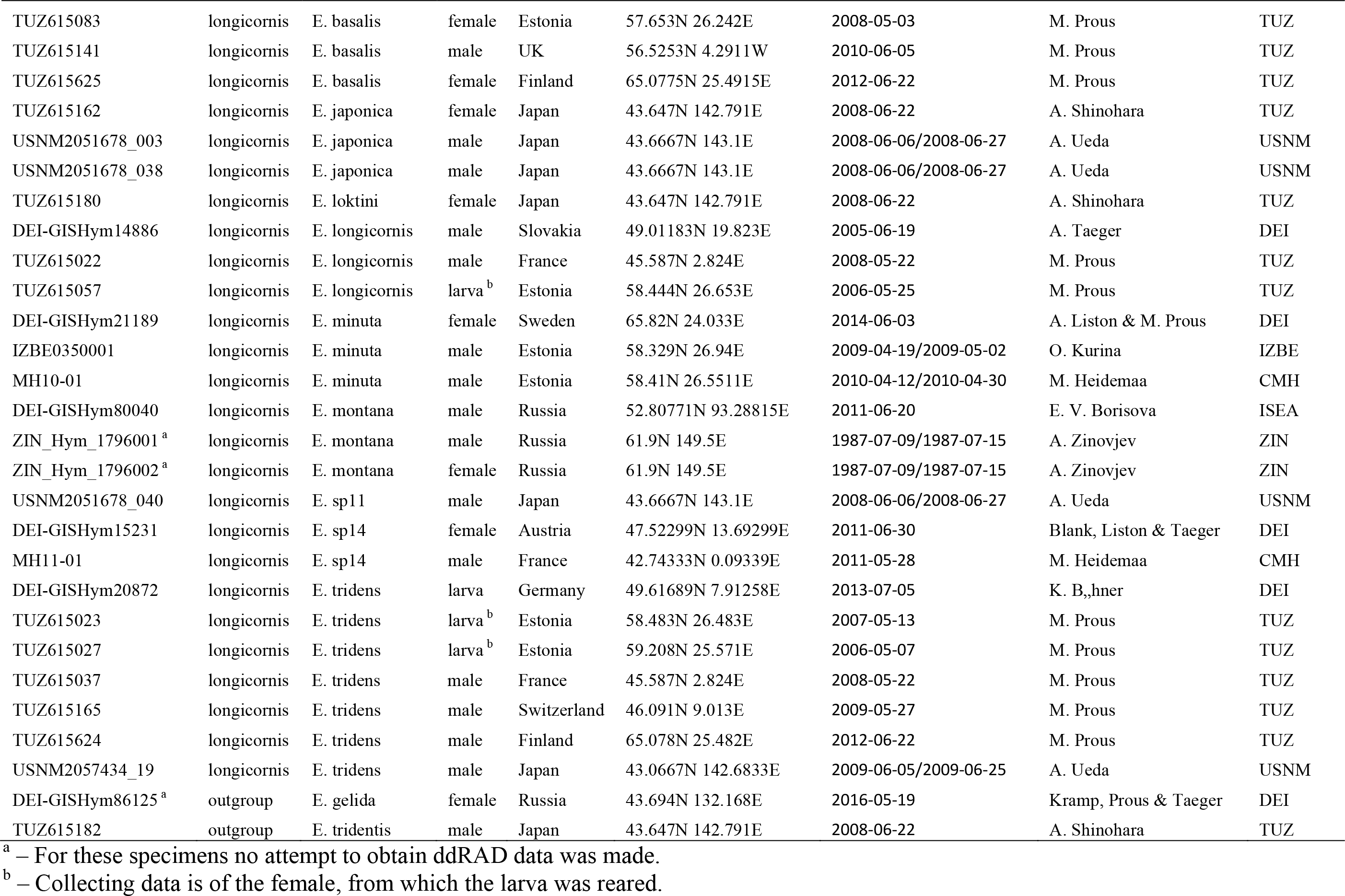

### 2.2 Sanger sequencing

To test congruence between mitochondrial and nuclear gene trees and to compare results based on Sanger sequencing of small number of genes with the genome-scale ddRAD sequencing, we initially amplified fragments of three genes, one mitochondrial and two nuclear. The mitochondrial gene used is complete (amplified and sequenced as described in Prous et al., 2011b) or partial cytochrome oxidase subunit I (COI). For most specimens, the sequenced fragment is at least 1078 bp. One specimen (BIOUG00998-B05, GenBank accession JX830389) had only the 658 bp fragment corresponding to the standard barcode region of the animal kingdom (Hebert et al., 2003). Complete or partial COI barcode sequences of three specimens (BIOUG17274-F06, BIOUG00998-D06, BIOUG00998-B05) were available in BOLD (http://www.boldsystems.org/), two of which were extended to 1078 bp by doing new DNA extractions, amplifications, and sequencing. The two nuclear markers are fragments of sodium/potassium-transporting ATPase subunit alpha (NaK) and DNA dependent RNA polymerase II subunit RPB1 (POL2). The NaK fragment used is a nearly complete sequence of its longest exon, 1654 bp. The POL2 fragment used is composed of two partial exons and one short intron that did not vary in length (87 bp) in the specimens studied here, altogether 2494–2710 bp, depending on the primer set used. After the first analyses of ddRAD data, we suspected cross-contamination in *Empria japonica*. To test this, we selected two variable candidate RAD loci that we suspected to be contaminated in *E. japonica* and designed primers to amplify and re-sequence these regions. One of the selected loci turned out to be a fragment of the zinc finger CCCH domain-containing protein 14 (ZC3H14) for which we designed additional primers to amplify its longest exon (containing also the ddRAD locus), varying between 1582–1639 bp in the studied specimens. For the second candidate locus (anonymous), quite a similar (around 80%) match was found only among the WGS (whole genome shotgun) contigs of *Neodiprion lecontei* (scaffold_346, GenBank accession LGIB01000346). This locus might be non-coding because of apparent frame-shifting indels in some ddRAD sequences, but was of the same length in the PCR amplified specimens, 138 bp. Primers used for amplification and sequencing are listed in Table 2. New POL2 and ZC3H14 primers (Table 2) were designed based on WGS contigs of four sawfly genomes (GenBank accessions AOFN01001568, LGIB01000323, AMWH01001469, AZGP01005167, AOFN02000929, LGIB01000132, AMWH01002139, AZGP02000664), sawfly transcriptomes published by Misof et al. (2014) and Peters et al. (2017), and based on POL2 sequences published by Malm and Nyman (2015). Numbers in the new POL2 and ZC3H14 primer names refer to the binding position of the primer’s 3’ end in the coding region of *Athalia rosae* mRNA (accessions XM_012395805 and XM_012401276). Primers for the anonymous locus were designed based on our ddRAD data.

PCR reactions were carried out in a total volume of 15–30 μl containing 1–2 μl of extracted DNA, 1.0–3.0 μl (5.0–15 pmol) of primers and 7.5–15 μl of 2x Multiplex PCR Plus Master mix (QIAGEN). The PCR protocol consisted of an initial DNA polymerase (HotStar Taq) activation step at 95 °C for 5 min, followed by 38–40 cycles of 30 s at 95 °C, 90 s at 49–59 °C depending on the primer set used, and 60–180 s (depending on the amplicon size) at 72 °C; the last cycle was followed by a final 30 min extension step at 68 °C. 3 μl of PCR product was visualised on a 1.4% agarose gel and then purified with FastAP and Exonuclease I (Thermo Scientific). 1.0–2.0 U of both enzymes were added to 12–27 μl of PCR solution and incubated for 15 min at 37 °C, followed by 15 min at 85 °C. 3–5 μl of purified PCR product per primer in a total volume of 10 μl (5–7 μl of sequencing primer at concentration 5 pmol/μl) were sent to Macrogen (Netherlands) for sequencing. Ambiguous positions (i.e. double peaks in chromatograms) due to heterozygosity or heteroplasmy were coded using IUPAC symbols. Sequences reported here have been deposited in the GenBank (NCBI) database (accession numbers MK299849–MK299982).

**Table 2.**
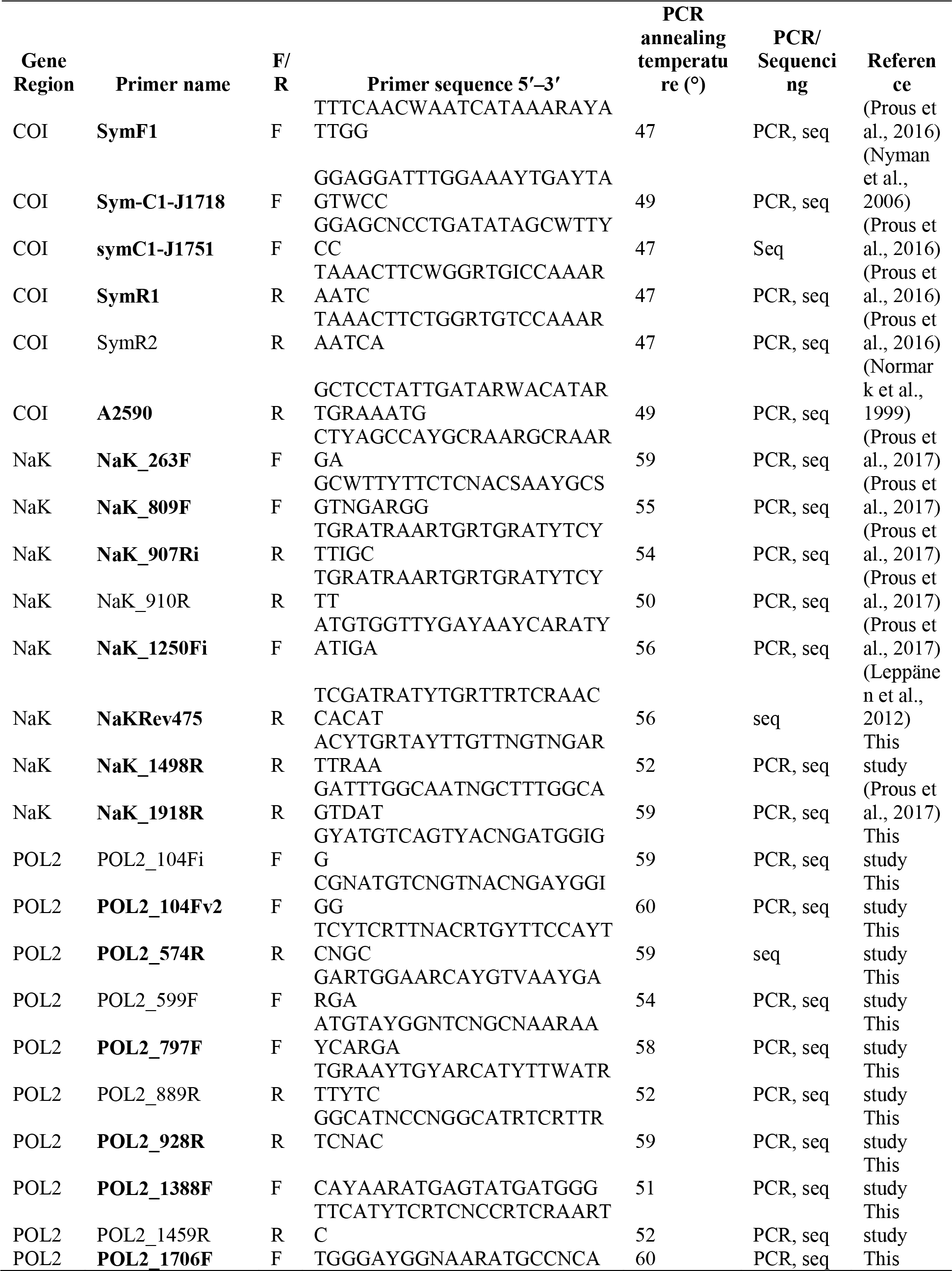
Primers used for PCR and sequencing (preferred primers in bold), with information provided on respective gene fragment, primer name, direction (forward, F or reverse, R), primer sequence, standard PCR annealing temperature, utilization (PCR/ sequencing), and reference. Primer annealing temperatures used for sequencing at Macrogen were 47°C for COI and 50°C for nuclear genes.

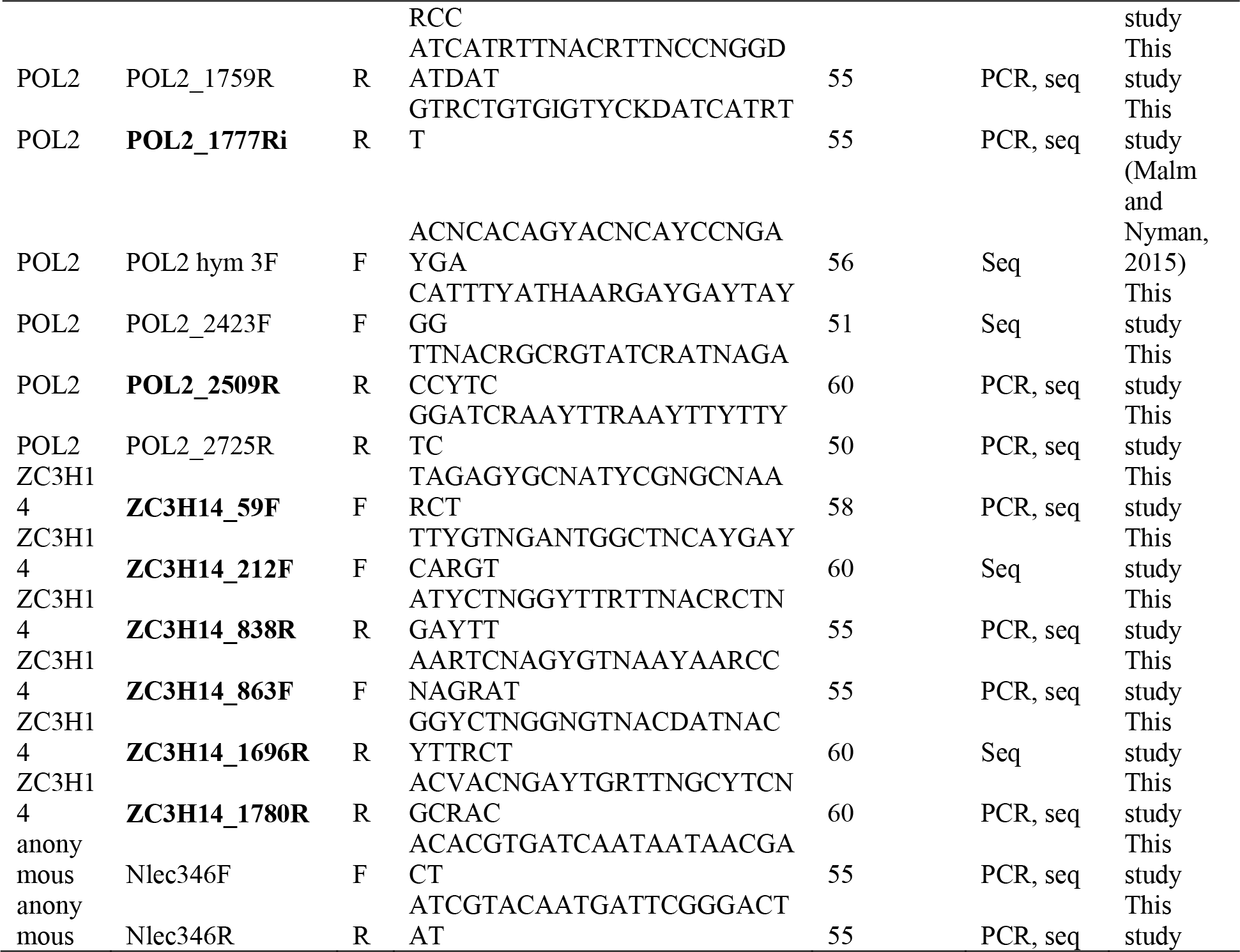

### 2.3 ddRADseq library preparation and bioinformatics

The quantity of genomic DNA (gDNA) was checked using PicoGreen kit (Molecular Probes). To reach sufficient gDNA quality and quantity, whole genome amplification was performed using REPLI-g Mini kit (Qiagen) due to the low concentrations of gDNA. The ddRADseq library was implemented following protocols described in (Lee et al., 2018) with an exception: the size distribution and concentration of the pools were measured with Bioanalyzer (Agilent Technologies). The de-multiplexed *Empria* fastq data are archived in the NCBI SRA: PRJNA505249 (Lee, 2018).

Raw paired-end reads were demultiplexed with no mismatches tolerated using their unique barcode and adapter sequences using *ipyrad* v.0.7.23 (Eaton and Overcast, 2016). All *ipyrad* defaults were used, with the following exceptions: the minimum depth at which majority rule base calls are made was set to 3, the clustering threshold was set to 0.95, the minimum number of samples that must have data at a given locus for it to be retained was set to 4, and the assembly method was set to denovo and reference for independent testing. The reference assembly method is mapped reads to *Athalia rosae* genome sequences (GenBank, GCA_000344095) with BWA using the default bwa-mem setting (Li, 2013) based on 95% of sequence similarity.

### 2.4 Phylogenetic analysis

Maximum likelihood (ML) trees were inferred in RAxML v.8.2.0 or v8.2.10 (Stamatakis, 2014), with bootstrap support estimated by a 1,000 replicates rapid bootstrap analysis from the unpartitioned GTR+GAMMA model. We visualized the resulting phylogeny and assessed bootstrap support using FigTree v.1.4.2 (Rambaut, 2015).

Maximum parsimony (MP) analyses were conducted using PAUP* 4.0b10 program (Swofford, 2003). All heuristic searches were performed with 1,000 replicates, employed the random addition of taxa, retained only the best tree, held 10 trees at each step using tree bisection-reconnection (TBR) branch swapping, collapsed zero-length branches, and using MULTREES. Bootstrap values were calculated using 1,000 replicates with the following options selected: heuristic search, TBR branch swapping, collapse of zero-length branches, and random-sequence-addition with one replicate.

Pairwise sequence divergence based on K2P distances were calculated using MEGA6 (Tamura et al., 2013) and the proportion of missing data was calculated using Mesquite (Maddison and Maddison, 2017). Net synonymous divergence between species was calculated with MEGA7 (Kumar et al., 2016).

### 2.5 Population structure and admixture

An admixture analysis was implemented in STRUCTURE v.2.3.1 (Pritchard et al., 2000) using SNP frequency data to better visualize genomic variation between individuals. Ten replicates were run at each value of *K* between 2 and 5 for *E. immersa* group and *K*=9 for *E. longicornis* group. Each run had a burn-in of 10K generations followed by 20K generations of sampling. We used StrAuto to automate Structure processing of samples (Chhatre and Emerson, 2017). Replicates were permuted in the program CLUMPP (Jakobsson and Rosenberg, 2007) according to the ad hoc Δ*K* statistics (Evanno et al., 2005), which is the second-order rate of change of the likelihood function. Structure results were visualized using the program DISTRUCT (Rosenberg, 2004).

We used four-taxon D-statistics (Durand et al., 2011) for introgression analysis. For the test, 1,000 bootstrap replicates were performed to measure the standard deviation of the D-statistics. Significance was evaluated by converting the Z-score (which represents the number of standard deviations from zero from D-statistics) into two tailed P-values, and using α=0.01 as a conservative cut-off for significance after correcting for multiple comparisons using Holm-Bonferroni correction. All D-statistics were calculated in pyRAD v.3.0.64 (Eaton, 2014). In order to run interactive data analysis, the Python Jupyter notebooks (http://jupyter.org) were used.

### 2.6 Cross-contamination detection

To detect identical loci between specimens or groups of specimens in ddRAD data, an R (R Core Team, 2017) script requiring a package *ape* (Paradis and Schliep, 2018) was written. The script takes as an input a text file containing alignments of ddRAD loci (output from *ipyrad*). A table is produced for every locus (rows) and specimen (columns) where cells contain list of specimens that are identical to a specimen indicated in the column (the cell is empty if there are no identical specimens for a particular specimen and locus). Additional columns are added to get information per locus about identical specimens between two groups, the number of specimens, maximum, median and mean divergence. The two groups examined are *longicornis* (including *E. tridentis*, which taxonomically is not a member of the group, but closely related) and *immersa* groups. For both groups and for every locus, specimens are recorded that are identical to any member in the other group while different from specimens in its own group. The second table produced by the script lists the specimens in the dataset, the number of loci, and the normalised number of loci per specimen. Normalised numbers of loci were calculated as half of the maximum number of loci divided by the number of loci of a particular specimen in the dataset. Then the script proceeds to produce bar plots (output as pdf) for every specimen showing percent of loci and normalised percent of loci that are identical to a particular specimen while different from all others. Two additional bar plots are produced for *longicornis* and *immersa* groups to show percent of loci of a particular specimen that are identical to any specimen in the wrong group while different from specimens in its own group. The script and the dataset with 19 413 loci (Supplementary Data S4, S11) are available on Figshare (http://dx.doi.org/10.6084/m9.figshare.7605404).

## 3. Results

### 3.1. Detection of divergent loci and cross-contamination

In the initial RAxML analysis of the dataset assembled with a clustering threshold of 80% similarity (29 859 loci, 5 945 539 bp; including *E. immersa* group) in the process of exploring RAD data, *E. japonica* did not form a monophyletic group, as the specimen USNM2051678-003 (hereafter as USNM003) was even outside of the *longicornis* group, forming a sister group to *E. tridentis* and the rest of *E. longicornis* group (Supplementary Data S1). Manual examination of the alignment regions present in at least one of the *E. japonica* specimens (about 600 000 bp) revealed that there were about 20 markers that contained regions that were not homologous to other specimens (divergence roughly 5–10 times higher than the average among the other specimens), mostly apparently because of mis-association of paired-end reads (Supplementary Data S2). In one case we also noticed that *E. japonica* specimen USNM003 was identical (when disregarding one indel in the middle of the locus, which can be explained by different lengths of quality trimmed paired-end reads) to *E. immersa* DEI-GISHym80071 while clearly different from all other specimens, indicating possible cross-contamination (p-distance to other *immersa* group specimens 0.6–3.4%, distance to *longicornis* group specimens 8.6–11.2%; Supplementary Data S3). Because most of the loci containing non-homologous regions could be detected with the criterion maximum p-distance more than 0.2, we created a dataset with these loci removed (19 413 loci, 3 517 320 bp). Except for within species relationships, the topology of ML tree based on this smaller dataset was identical to the initial tree (Supplementary Data S4).

Because the phylogenetic position of *E. japonica* specimen USNM003 did not change when the divergent loci were removed, we aimed next to detect possible cross-contamination in the smaller dataset. For this, we identified pairs of specimens that were identical to each other while different from the rest. In addition, to specifically get an idea about the level of cross-contamination between *immersa* and *longicornis* groups, we identified loci in which a particular specimen was identical to any member from the group it does not belong to, e.g. *E. japonica* specimen USNM003 being identical to at least one specimen in *immersa* group while different from the others in *longicornis* group (if present).

Clear outlier with regard to cross-contamination between *immersa* and *longicornis* groups was *E. japonica* USNM003, with 26.7% of its loci (out of 1015) identical to one or more specimens in *immersa* group (Fig. 1a). The cross-contamination in USNM003 seems to be caused by *E. immersa* DEI-GISHym80071 (Fig. 1b) which alone contributes 8.4% of the sequences of USNM003 (there were only two other *immersa* group specimens, both with only one sequence (0.1%) identical to USNM003). The other specimens with possibly significant amount of cross-contamination were *E. tridens* TUZ615037 (5.6% of loci identical to *E. tridentis* TUZ615182, which is not a member of *immersa* or *longicornis* groups), *E. immersa* DEI-GISHym80045 (5.1% identical to any in *longicornis* group), *E. fletcheri* TUZ615334 (4.7% identical to any in *longicornis* group), *E. plana* TUZ615181 (4.4% identical to any in *longicornis* group). For the other specimens these percentages are less than 2.5% and in most cases less than 1% (Supplementary Data S5).

**Fig. 1.**
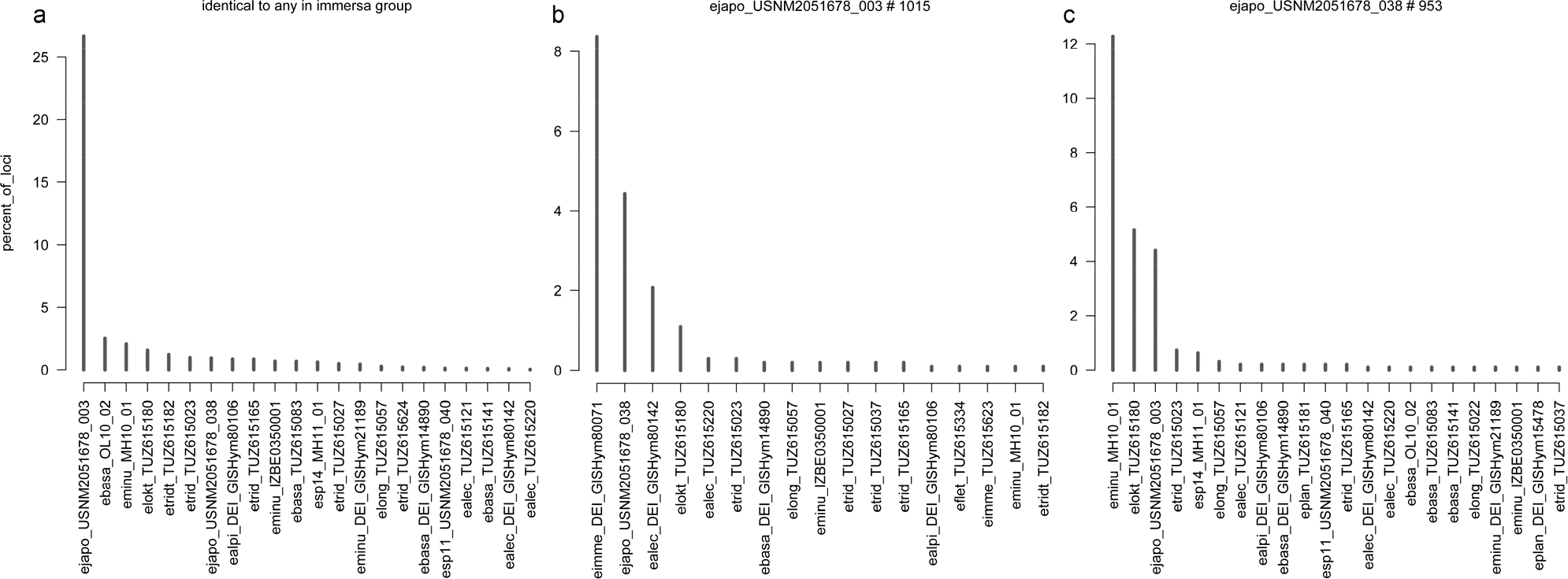
Examples of results of cross-contamination check by counting identical loci among *Empria* specimens (dataset with 19 413 loci, Supplementary Data S4). For results of all samples see Supplementary Data S5. (a) Percent of loci in every *longicornis* group specimen (X-axis) (including *E. tridentis*) that are identical to any specimen in *immersa* group, while different from other specimens in *longicornis* group (if present). (b, c) Percent of loci in *E. japonica* USNM2051678_003 (b) and USNM2051678_038 (c) that are identical to a particular specimen (listed along X-axis) while different from the others in the dataset.

It is more problematic to estimate level of cross-contamination within the *immersa* and *longicornis* groups themselves, because of higher degree of relatedness and at least occasional hybridisations between the species cannot be excluded. Nevertheless, for *longicornis* group it seems that in most cases the specimens are free of cross-contamination. For most specimens, the proportions of loci that were identical to a particular specimen belonging to a different species were less than 2.5% (Supplementary Data S5). Clear exception was *E. japonica* USNM2051678-038 (hereafter as USNM038) with 12.3% of its loci identical to *E. minuta* MH10-01 and 5.1% identical to *E. loktini* (Fig. 1c). The other exception was pair of *E. longicornis* TUZ615057 (4.8% of its loci identical to TUZ615023) and *E. tridens* TUZ615023 (2.9% of its loci identical to TUZ615057), but this might be genuine because these species are closely related and the specimen TUZ615023 seems to be highly heterozygous (0.48% of two-fold degenerate positions; Supplementary Data S6) which increases the chance for sequences to be identified as identical.

For *E. japonica* USNM038, however, the high amount of loci that were identical to *E. loktini* (5.1%) and especially to a particular specimen of *E. minuta* MH10-01 (12.3%) was suspicious. From a morphological perspective, *Empria japonica* is not expected to be specifically related to *E. loktini* or *E. minuta*, but is very similar to *E. tridens* and *E. longicornis*. Indication of cross-contamination is also the fact that practically only one *E. minuta* specimen from Estonia contributed the identical sequences. The other two *E. minuta* specimens from Estonia and Sweden both contributed only one (0.1%) identical locus. To further check the possibility of contamination, we selected two candidate ddRAD loci of USNM038 to see if the results can be replicated using PCR and Sanger sequencing. In the ddRAD alignments, one of the selected sequences of USNM038 was identical to *E. loktini* while clearly different from other species, and the other had seven ambiguous positions (two Ns and five two-fold degenerate positions), which made it identical to 9 (out of 23) other specimens, including *E. loktini* and *E. minuta* MH10-01. In both cases, the Sanger sequencing revealed that *E. japonica* specimens were identical to each other and different from all other specimens, confirming the pattern found in the other Sanger sequenced nuclear markers (Supplementary Data S7). For the other sequenced specimens, the Sanger sequencing results were found to be consistent with the ddRAD data (no substitution differences), but in some cases there were indel differences, which can at least partly be explained by length differences of quality trimmed paired-end reads, because at least some of them were in the middle of the locus and different ddRAD assemblies produced different lengths (Supplementary Data S7).

It is more difficult to recognise possible cross-contamination within *E. immersa* group, because of small number of specimens and unresolved taxonomy. Nevertheless, based on morphology and ecology, *E. fletcheri* can be reliably separated from the others in *immersa* group. At least the two European specimens of *E. fletcheri* (from Estonia and Sweden) do not share significant amount of identical loci with a particular specimen from the other species (for each of these specimens the contribution is less than 2.2%; Supplementary Data S5). However, the *E. fletcheri* specimen from Canada (BIOUG17274-F06), which genetically seems to have little in common with the European counterpart, might be somewhat contaminated because the largest contributor of identical loci and in quite a large amount (3.4%; Supplementary Data S5) is *E. camtschatica* from Sweden (DEI-GISHym80070). For other cases in *immersa* group it is difficult to evaluate if it is a genuine signal or cross-contamination. For example *E. plana* DEI-GISHym15478 has 7.8% of its loci identical to *E. camtschatica* DEI-GISHym80070 (Supplementary Data S5), but both are from Sweden and could be the same species despite of some differences in the saws (ovipositors).

We subsequently analysed *immersa* and *longicornis* groups separately using datasets assembled with clustering threshold of 95% similarity (both, *de novo* and reference assembly), which decreases the chance of introducing highly divergent regions (Tables 3 and 4). Because both *E. japonica* specimens were apparently substantially more cross-contaminated than the other specimens, we decided to exclude them from final analyses, but also examined the effects of including one or both of them in different analyses. We excluded also apparently the third most contaminated specimen *E. tridens* TUZ615037 from the final analyses (5.6% of loci identical to *E. tridentis* TUZ615182, which is not a member of *immersa* or *longicornis* groups), although its inclusion had almost no effect on the results.

**Table 3.**
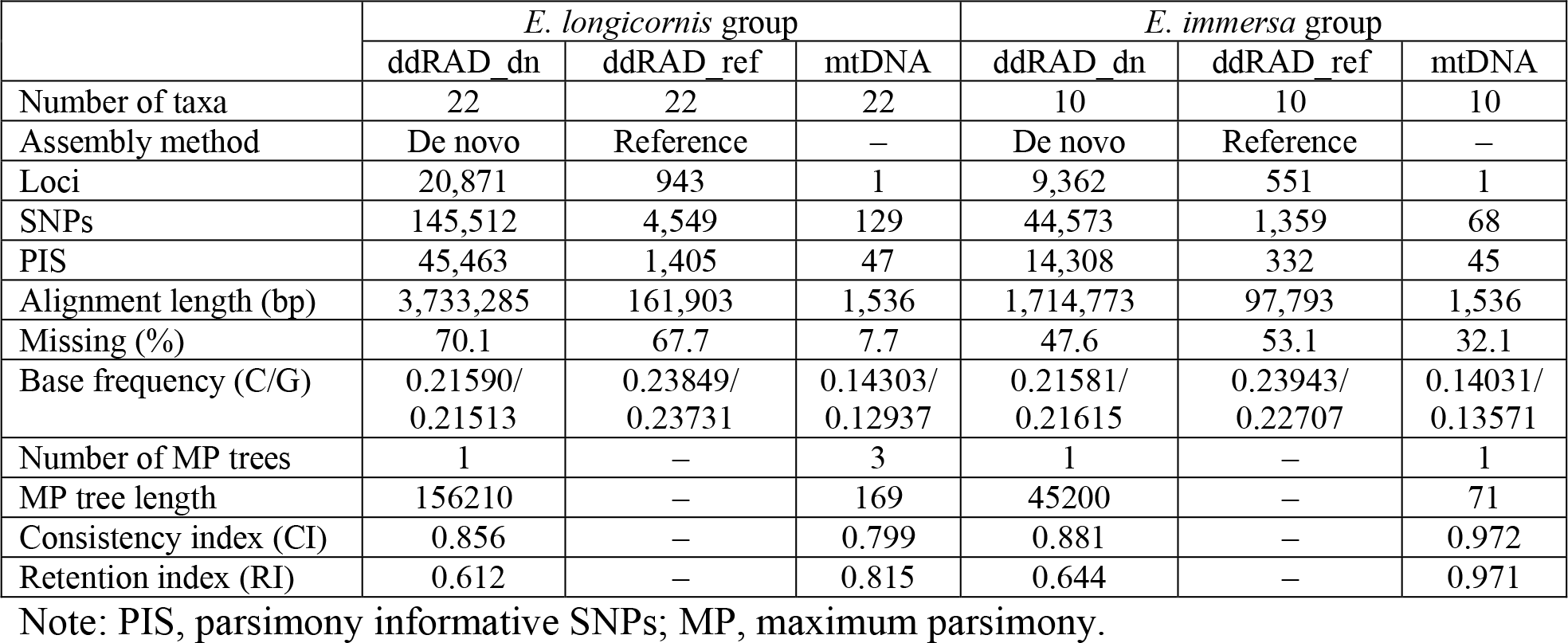
Summary statistics of ddRAD and mitochondrial COI barcode data sets from *E. longicornis* and *E. immersa* group.

**Table 4.** Specimens of *Empria* analysed in this study and a summary of the ddRAD data in *de novo* and reference assembly.

| Species | Sample ID | Total reads<br>(million) | de novo assembly |  |  |  | Reference assembly |  |  |  |  |
| --- | --- | --- | --- | --- | --- | --- | --- | --- | --- | --- | --- |
|  |  |  | Clusters<br>at 95%<sup>a</sup> | Mean<br>depth | Retained<br>loci<sup>b</sup> | Recovered<br>loci | Mapped<br>reads | Clusters<br>total | Clusters<br>depth | Reads<br>consensus | Recovered<br>loci in<br>assembly |
| (a) *Empria longicornis* group |  |  |  |  |  |  |  |  |  |  |  |
| *E. alector* | ealec_DEI_GISHym80142 | 4.78 | 227253 | 20.1 | 59694 | 13725 | 28997 | 8165 | 14.9 | 2699 | 618 |
| *E. alector* | ealec_TUZ615036 | 0.34 | 14426 | 11.3 | 4576 | 2139 | 2163 | 422 | 26.6 | 133 | 70 |
| *E. alector* | ealec_TUZ615121 | 4.81 | 146273 | 31.1 | 42238 | 13311 | 17241 | 5081 | 17.6 | 1782 | 620 |
| *E. alector* | ealec_TUZ615220 | 2.18 | 39319 | 51.6 | 11234 | 4894 | 1 | 1 | 4.0 | 1 | NA |
| *E. alpina* | ealpi_DEI_GISHym80106 | 4.24 | 154599 | 26.2 | 45944 | 6777 | 19941 | 5475 | 17.1 | 2029 | 430 |
| *E. basalis* | ebasa_DEI_GISHym14890 | 0.72 | 41354 | 15.4 | 16358 | 6920 | 2166 | 1245 | 5.8 | 437 | 267 |
| *E. basalis* | ebasa_OL10_02 | 1.01 | 22225 | 34.5 | 4675 | 1628 | 718 | 520 | 2.9 | 124 | 62 |
| *E. basalis* | ebasa_TUZ615083 | 0.09 | 9336 | 8.9 | 2817 | 1354 | 487 | 284 | 2.4 | 72 | 47 |
| *E. basalis* | ebasa_TUZ615141 | 0.86 | 50972 | 15.4 | 18278 | 8815 | 3397 | 1805 | 5.9 | 630 | 362 |
| *E. loktini* | elokt_TUZ615180 | 3.73 | 87639 | 38.4 | 29215 | 3204 | 9162 | 2681 | 13.7 | 1049 | 230 |
| *E. longicornis* | elong_TUZ615022 | 2.35 | 26206 | 74.4 | 5274 | 1888 | 2710 | 362 | 6.7 | 158 | 77 |
| *E. longicornis* | elong_TUZ615057 | 7.94 | 169150 | 43.4 | 60811 | 12411 | 27292 | 5494 | 34.5 | 2648 | 624 |
| *E. minuta* | eminu_DEI_GISHym21189 | 0.47 | 15693 | 23.2 | 8430 | 1701 | 2245 | 615 | 13.9 | 321 | 110 |
| *E. minuta* | eminu_IZBE0350001 | 0.99 | 17678 | 50.7 | 7576 | 1712 | 678 | 129 | 37.7 | 95 | 63 |
| *E. minuta* | eminu_MH10_01 | 2.47 | 67598 | 35.0 | 30890 | 5477 | 8133 | 2266 | 16.3 | 1280 | 366 |
| *E. sp. 11* | esp11_USNM2051678_040 | 0.34 | 10506 | 23.1 | 4687 | 904 | 542 | 337 | 3.1 | 137 | 39 |
| *E. sp. 14* | esp14_MH11_01 | 1.34 | 47172 | 26.7 | 23470 | 6544 | 3217 | 857 | 31.0 | 495 | 287 |
| *E. tridens* | etrid_TUZ615023 | 2.46 | 88462 | 27.0 | 42207 | 14400 | 12408 | 4209 | 14.0 | 2184 | 732 |
| *E. tridens* | etrid_TUZ615027 | 2.23 | 81560 | 26.5 | 36613 | 12705 | 13373 | 3713 | 19.5 | 1919 | 645 |
| *E. tridens* | etrid_TUZ615165 | 7.32 | 87708 | 65.9 | 29019 | 10128 | 7056 | 2643 | 10.4 | 1257 | 507 |
| *E. tridens* | etrid_TUZ615624 | 1.18 | 25886 | 42.8 | 9701 | 3738 | 1815 | 896 | 8.8 | 407 | 169 |
| *E. tridentis* | etridt_TUZ615182 | 2.93 | 71026 | 34.6 | 28846 | 2866 | 5943 | 2607 | 8.9 | 1308 | 270 |
|  | AVERAGE | 2.49 | 68275 | 33.0 | 23752 | 6238 | 7713 | 2264 | 14.3 | 962 | 314 |
| (b) *Empria immersa* group |  |  |  |  |  |  |  |  |  |  |  |
| *E. camtschatica* | ecamt_DEI_GISHym80070 | 1.22 | 48116 | 24.2 | 17103 | 6835 | 3061 | 1591 | 4.9 | 562 | 311 |
| *E. fletcheri* | eflet_BIOUG17274_F06 | 11.77 | 162626 | 68.7 | 26945 | 3803 | 3008 | 1474 | 9.1 | 415 | 227 |
| *E. fletcheri* | eflet_DEI_GISHym31039 | 0.51 | 30181 | 14.6 | 10985 | 4306 | 2333 | 1131 | 8.0 | 398 | 215 |
| *E. fletcheri* | eflet_TUZ615334 | 0.27 | 28908 | 9.2 | 9900 | 3805 | 1735 | 1083 | 4.2 | 306 | 187 |
| *E. immersa* | eimme_DEI_GISHym80045 | 0.98 | 53371 | 17.1 | 20648 | 7866 | 4804 | 2265 | 14.3 | 823 | 429 |
| *E. immersa* | eimme_DEI_GISHym80071 | 4.09 | 124234 | 30.7 | 41156 | 8318 | 13554 | 4017 | 21.5 | 1697 | 470 |
| *E. immersa* | eimme_TUZ615623 | 2.96 | 79575 | 34.0 | 33109 | 8058 | 8117 | 2695 | 22.8 | 1231 | 453 |
| *E. improba* | eimpr_BIOUG00998_D06 | 0.39 | 12918 | 27.2 | 4762 | 1414 | 668 | 379 | 3.2 | 146 | 69 |
| *E. plana* | eplan_DEI_GISHym15478 | 1.84 | 24812 | 29.4 | 9043 | 2594 | 1374 | 488 | 8.6 | 268 | 110 |
| *E. plana* | eplan_TUZ615181 | 0.58 | 20249 | 26.3 | 8695 | 2109 | 2226 | 599 | 10.1 | 275 | 93 |
|  | **AVERAGE** | **2.46** | **58499** | **28.1** | **18235** | **4911** | **4088** | **1572** | **10.7** | **612** | **256** |

<sup>a</sup>Clusters that passed filtering for 6x minimum coverage.
<sup>b</sup>Loci retained after passing coverage and paralog filters.
NA, not applicable.

### 3.2. Empria longicornis group

All species with more than one individual sampled are found to be monophyletic and in most cases strongly supported in maximum likelihood trees reconstructed from ddRAD datasets based on *de novo* and reference assemblies (Figs 2a and 3a). Monophyly of only *E. tridens* is moderately supported based on reference assembly (Fig. 3a). There are some differences in tree topology above the species level based on reference and *de novo* assembly (phylogenetic positions of *E. sp11* and *E. loktini*), but these differences are poorly supported, particularly in the smaller dataset based on the reference assembly (Figs 2a and 3a).

**Fig. 2.**
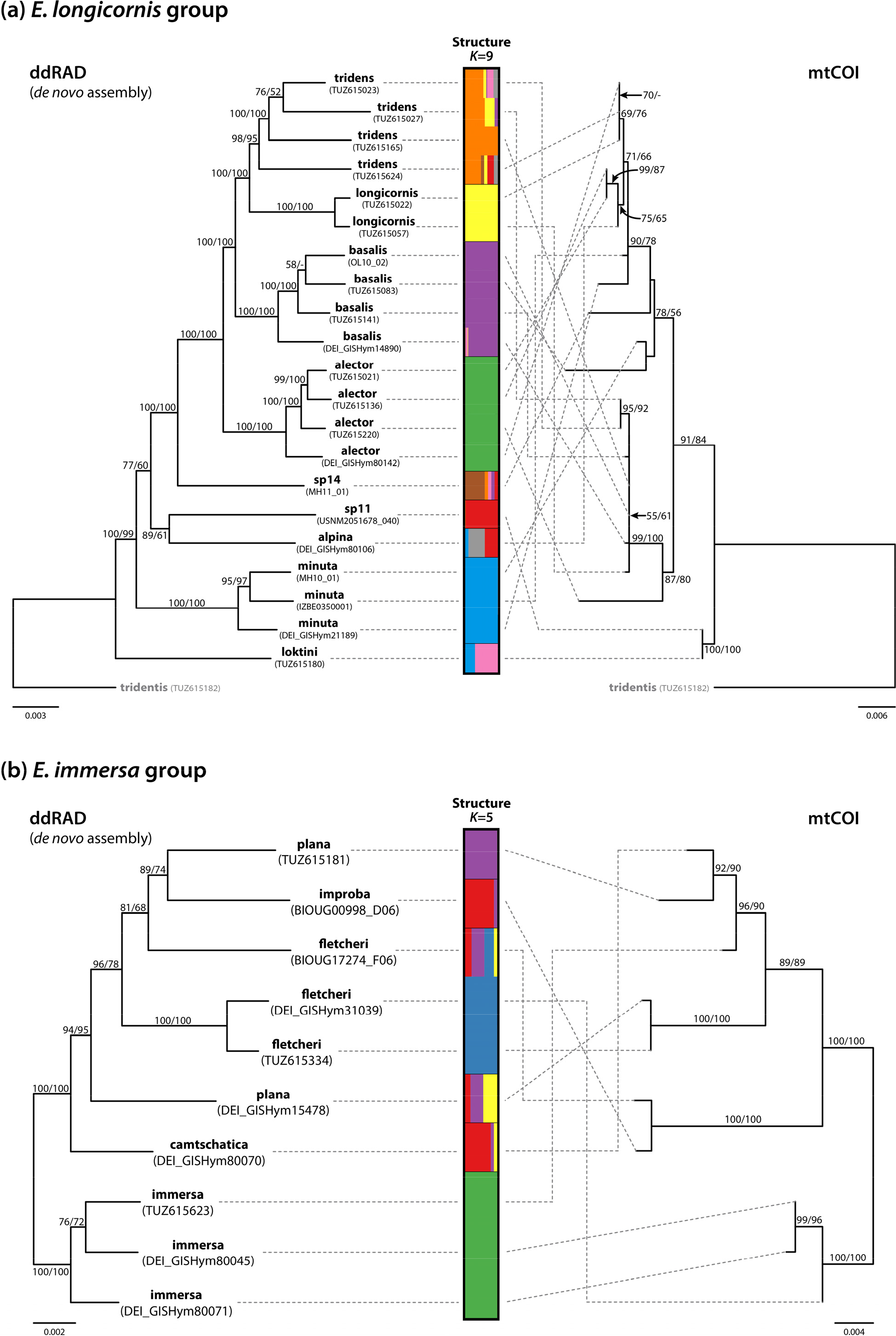
Maximum likelihood trees of two *Empria* species groups and population admixture analyses based on ddRAD *de novo* assemblies in comparison with mitochondrial COI maximum likelihood tree (1536 bp). (a) *E. longicornis* group (ddRAD dataset with 20 871 loci). (b) *E. immersa* group (ddRAD dataset with 9 362 loci). Bootstrap support values (%) below or above branches resulting from maximum likelihood (ML) and maximum parsimony (MP) analyses are shown as ML/MP.

**Fig. 3.**
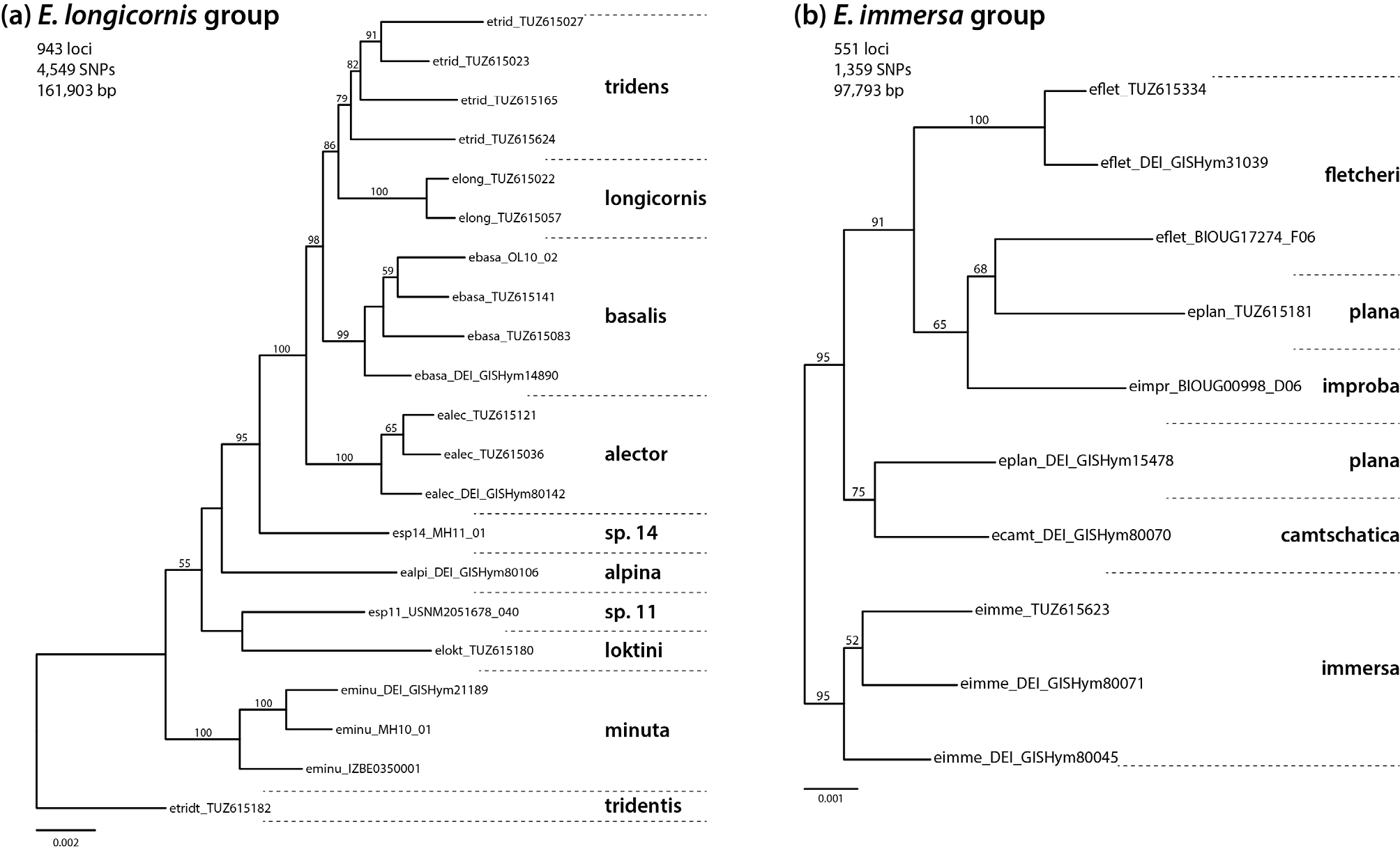
Maximum likelihood trees of two *Empria* species groups based on ddRAD reference assembly. (a) *E. longicornis* group. (b) *E. immersa* group.

Monophyly of most species defined based on morphology is also well supported by concatenated analysis of three nuclear protein coding genes obtained by Sanger sequencing (Fig. 4d). Of the species for which more than one individual was sampled, only *E. tridens* is not monophyletic according to three-gene tree. The species for which only one individual was sampled (*E. alpina*, *E. loktini*, sp11) are well separated from the other species as well as from each other based both on ddRAD and Sanger data, supporting their species status (Figs 2a, 3a, 4d). For *E. montana* Koch, 1984 we were not able to obtain enough ddRAD data to be included in the analyses (Supplementary Data S8), but based on Sanger sequencing of three specimens, we can confirm that it belongs to *longicornis* group (Supplementary Data S9 and S10). *Empria montana* was not recognised by Prous et al. (2011b) as a member of *longicornis* group because of divergent penis valve (only holotype male was known at the time). Two of the studied specimens of *E. montana* (http://dx.doi.org/10.6084/m9.figshare.7447847; http://dx.doi.org/10.6084/m9.figshare.7447874) were reared from *Dasiphora fruticosa*, which was previously unknown. Based on current genetic sampling of three *E. montana* specimens (two from Magadan oblast, one from Krasnoyarsk Krai, Russia), this species is monophyletic according to mitochondrial COI (Supplementary Data S10). Partial fragments of three nuclear genes used here are available for only one *E. montana* specimen, based on which this species groups together with *E. alpina*, *E. minuta*, and sp11, but in unresolved position (Supplementary Data S9).

**Fig. 4.**
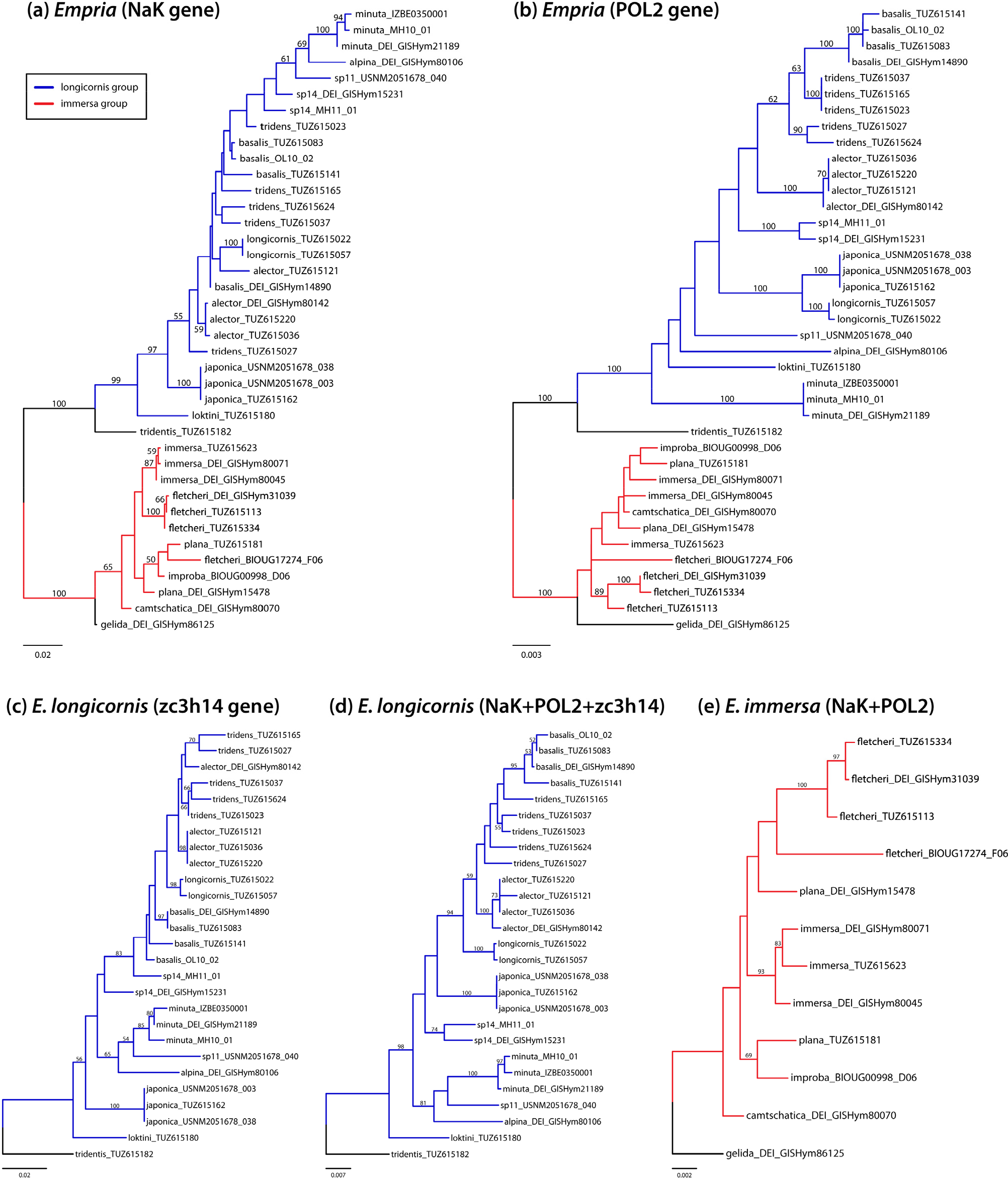
Maximum likelihood trees based on three nuclear protein coding genes obtained by Sanger sequencing. (a) NaK (1654 bp). (b) POL2 (2494 bp). (c) ZC3H14 (alignment 1654 bp). (d) Concatenated NaK, POL2, and ZC3H14 (alignment 5802 bp) for *longicornis* group. (e) Concatenated NaK and POL2 (4148 bp) for *immersa* group.

Distance calculations of ddRAD data (*de novo* assembly) are also consistent with species limits defined based on morphology (Table 5). Mean within species divergence varies between 0.18– 0.92%, while distances among species are about twice as high, 1.05–1.87%.

**Table 5.** Mean pairwise distances of ddRAD (below diagonal) and mtDNA data (above diagonal) within (grey shaded cells; ddRAD de novo assembly data/mtDNA) and between the species.

(a) *E. longicornis* group and *E. tridentis*
|  | alec | alpi | basa | lokt | long | minu | sp11 | sp14 | trid | tridt |
| --- | --- | --- | --- | --- | --- | --- | --- | --- | --- | --- |
| alecor | 0.35/1.20 | 0.74 | 1.54 | 1.82 | 1.00 | 1.07 | 1.82 | 1.80 | 1.36 | 5.07 |
| alpina | 1.78 | NA | 1.58 | 1.76 | 0.79 | 0.59 | 1.76 | 1.67 | 1.21 | 5.01 |
| basalis | 1.08 | 1.70 | 0.50/1.01 | 1.58 | 0.93 | 1.51 | 1.90 | 1.90 | 0.94 | 5.26 |
| loktini | 1.87 | 1.83 | 1.77 | NA | 1.53 | 1.50 | 0 | 1.95 | 1.66 | 4.56 |
| longicornis | 1.24 | 1.77 | 1.13 | 1.80 | 0.18/1.24 | 0.99 | 1.53 | 1.63 | 0.74 | 5.05 |
| minuta | 1.73 | 1.70 | 1.61 | 1.85 | 1.69 | 0.52/0.79 | 1.50 | 1.49 | 1.33 | 4.87 |
| sp.11 | 1.82 | 1.60 | 1.72 | 1.73 | 1.87 | 1.69 | NA | 1.95 | 1.66 | 4.56 |
| sp.14 | 1.55 | 1.60 | 1.53 | 1.84 | 1.52 | 1.75 | 1.79 | NA | 1.86 | 5.34 |
| tridens | 1.17 | 1.73 | 1.05 | 1.84 | 1.09 | 1.74 | 1.84 | 1.52 | 0.92/0.97 | 5.23 |
| tridentis | 2.08 | 2.07 | 1.99 | 1.99 | 2.10 | 2.10 | 2.02 | 2.08 | 2.01 | NA |

(b) *E. immersa* group
|  | camt | flet | imme | impr | plan |
| --- | --- | --- | --- | --- | --- |
| camtschatica | NA | 2.30 | 1.90 | 2.74 | 1.42 |
| fletcheri | 1.14 | 0.94/2.88 | 1.99 | 2.24 | 2.39 |
| immersa | 0.88 | 1.34 | 0.49/1.21 | 2.68 | 2.47 |
| improba | 1.18 | 1.31 | 1.42 | NA | 3.72 |
| plana | 1.02 | 1.31 | 1.22 | 1.17 | 1.15/2.02 |

NA, not applicable because of single specimen.

Based on nuclear NaK, POL2, and ZC3H14 (altogether 5646 bp), net synonymous divergences among most of the species (7 species for which more than one individual was sampled) were between 2.3–9.4%, suggesting that these species are well separated (Roux et al., 2016). Divergence between only *E. basalis* and *E. tridens* (1.5%) falls within the grey zone of speciation according to Roux et al. (2016).

Phylogenies based on ddRAD and combined data of three nuclear protein coding genes (NaK, POL2, and ZC3H14) were largely congruent when considering well supported relationships (bootstrap support more than 70%). Moderately supported differences involved relationships among *E. minuta*, *E. alpina*, and sp11, which formed a clade based on the three-gene dataset (Fig. 4d), but not based on ddRAD data (Figs 2a and 3a). There are some strongly or moderately supported phylogenetic differences among the three protein coding genes. According to the gene tree of POL2 (Fig. 4b), *E. longicornis* and *E. japonica* form strongly supported clade, which is absent in the other two gene trees (NaK and ZC3H14) (Figs 4a and 4c). The second, moderately supported difference is non-monophyly of sp14 according to ZC3H14 (Fig. 4c), contrary to POL2 (Fig. 4b) (according to NaK it is unresolved). Another, moderately supported difference is between NaK and ZC3H14 on one hand and POL2 on the other. According to NaK and ZC3H14 (Figs 4a and 4c), *E. alpina*, *E. minuta*, and sp11 form a clade (although relationships among these three species differ, but without strong support), but *E. minuta* is weakly supported as basal to all other *longicornis* group species in POL2 tree (Fig. 4b). These differences between single genes can be expected in closely related species complexes because of incomplete lineage sorting, which can cause incompatibilities between gene and species trees even without hybridisations.

Admixture analysis with STRUCTURE at *K*=9 (the number of species based on morphology, excluding *E. japonica*) supports most species as largely separate populations from each other (Fig. 2a). Best supported are *E. alector*, *E. basalis*, *E. longicornis*, *E. minuta* and sp11, which appear to have very little or no contribution from other species (Fig. 2a). Reasonably well supported are *E. loktini* and sp14, while *E. tridens* and *E. alpina* apparently have significant contributions from some other species (Fig. 2a). However, there are inconsistencies among different STRUCTURE analyses. When both *E. japonica* specimens are included (at *K*=10), *E. alpina* is better supported, while *E. longicornis* receives less support, as it seems to have a large contribution from *E. basalis* (Supplementary Data S11).

The results of four-taxon D-statistic tests suggest numerous cases of introgressions between different species. The most strongly supported case involves *E. minuta* and sp14 (Table 6), but curiously this receives (almost) no support from STRUCTURE analyses (Fig. 2a, Supplementary Data S11). In contrast to STRUCTURE (Fig. 2a) and D-statistic tests (Table 6), the counting of the number of identical loci between the specimens suggests the largest contributor as *E. alpina* (1.5%) in case of sp14 (*E. minuta* contributes 0.5%, Supplementary Data S5), which finds some support in the STRUCTURE analysis when both *E. japonica* specimens are included (Supplementary Data S11). There are numerous other inconsistencies among the four-taxon D-statistic tests, different STRUCTURE analyses and the counting of identical loci.

**Table 6.** Four-taxon D-statistic tests results showing significant replicates for introgression in *Empria*.

| Test | P1<sup>1</sup> | P2 | P3 | O | Range Z<sup>2</sup> | nSig/n<sup>3</sup> | nSig/n (%)<sup>4</sup> |
| --- | --- | --- | --- | --- | --- | --- | --- |
| **(a) *E. longicornis* group (28 cases out of 44)** |  |  |  |  |  |  |  |
| a1 | T | T | L | Tt | 0.1 – **15.9** | 5/11 | 45.5 |
| a2 | T | T | B | Tt | 0.3 – **15.4** | 7/23 | 30.4 |
| a3 | T | T | Ac | Tt | 0.0 – **10.2** | 4/17 | 23.5 |
| a4 | T | T | Sp14 | Tt | 0.1 – **17.5** | 2/5 | 40.0 |
| a5 | T | T | Sp11 | Tt | 0.3 – **9.4** | 2/5 | 40.0 |
| a6 | T | T | M | Tt | 0.0 – **20.8** | 6/17 | 35.3 |
| a7 | T | T | Ap | Tt | 0.0 – **6.9** | 1/5 | 20.0 |
| a8 | T | T | Lt | Tt | 0.1 – **5.6** | 2/5 | 40.0 |
| a9 | L | L | B | Tt | 1.5 – **3.3** | 1/3 | 33.3 |
| a10 | L | L | M | Tt | 0.5 – **18.8** | 1/3 | 33.3 |
| a11 | B | B | T | Tt | 0.0 – **13.6** | 2/23 | 8.7 |
| a12 | B | B | L | Tt | 0.0 – **3.5** | 1/11 | 9.1 |
| a13 | B | B | Ac | Tt | 0.0 – **8.2** | 3/17 | 17.6 |
| a14 | B | B | sp14 | Tt | 0.2 – **13.3** | 3/5 | 60.0 |
| a15 | B | B | sp11 | Tt | 0.0 – **3.5** | 1/5 | 20.0 |
| a16 | B | B | M | Tt | 0.5 – **6.4** | 1/5 | 20.0 |
| a17 | B | B | Lt | Tt | 0.0 – **10.2** | 4/17 | 23.5 |
| a18 | Ac | Ac | B | Tt | 0.0 – **4.6** | 1/11 | 9.1 |
| a19 | M | M | T | Tt | 0.0 – **17.2** | 2/11 | 18.2 |
| a20 | M | M | L | Tt | 0.2 – **3.8** | 1/5 | 20.0 |
| a21 | M | M | B | Tt | 0.1 – **12.3** | 3/11 | 27.3 |
| a22 | M | M | Ac | Tt | 0.0 – **3.9** | 2/8 | 25.0 |
| a23 | M | M | Sp14 | Tt | 17.8 – **38.5** | 3/3 | 100.0 |
| a24 | M | M | Sp11 | Tt | 0.0 – **10.0** | 1/3 | 33.3 |
| a25 | (T+L) | (T+L) | B | Tt | 0.0 – **15.3** | 26/59 | 44.1 |
| a26 | B | B | (T+L) | Tt | 0.0 – **13.5** | 2/35 | 5.7 |
| a27 | (T+L+B) | (T+L+B) | Ac | Tt | 0.0 – **13.7** | 23/134 | 17.2 |
| a28 | Ac | Ac | (T+L+B) | Tt | 0.0 – **4.7** | 1/29 | 3.4 |
| **(b) *E. immersa* group (7 cases out of 26)** |  |  |  |  |  |  |  |
| b1 | F | F | C | Im | 0.0 – **3.5** | 1/3 | 33.3 |
| b2 | Im | Im | F | Ip | 0.4 – **8.8** | 1/8 | 12.5 |
| b3 | Im | Im | Ip | F | 0.0 – **3.5** | 1/5 | 20.0 |
| b4 | (F+Ip) | (F+Ip) | (C+P) | Im | 0.0 – **3.8** | 1/19 | 5.3 |
| b5 | (C+P) | (C+P) | (F+Ip) | Im | 2.3 – **5.4** | 3/4 | 75.0 |
| b6 | (F+Ip) | (F+Ip) | Im | P | 0.0 – **6.0** | 6/29 | 20.7 |
| b7 | (F+Ip) | (F+Ip) | Im | C | 0.0 – **6.7** | 3/29 | 10.3 |

<sup>1</sup>Taxon names are abbreviated: In *E. longicornis* group, Ac: *E. alector*, Ap: *E. alpina*, B: *E. basalis*, L: *E. longicornis*, Lt: *E. loktini*, M: *E. minuta*, Sp11: *Empria* sp.11, Sp14: *Empria* sp.14, T: *E. triden*, Tt: *E. tridentis*. In *E. immersa* group, C: *E. camtschatica*, F: *E. fletcheri*, Im: *E. immersa*, Ip: *E. improba*, P: *E. plana*. Tests are referred to by number in the text. Insignificant cases were not shown.
<sup>2</sup>Bold indicates significance at $\alpha=0.01$ .
<sup>3</sup>Significant tests over possible sampled individuals.
<sup>4</sup>The percentage was calculated for the number of significant replicates shown (nSig) out of all possible four-sample replicates (n) in each test.

In contrast to nuclear data, mitochondrial COI does not support monophyly of any species (except possibly *E. montana* and sp14: Supplementary Data S10) and in some cases the non-monophyly is strongly supported (Fig. 2a). Besides the non-monophyly of species, the general topology of COI tree is very different from nuclear ML tree (Fig. 2a).

### 3.3. Empria immersa group

Morphologically defined species boundaries in *immersa* group (Prous et al., 2014) are not as well supported as in *longicornis* group. Only *E. immersa* based on ddRAD data and combined analyses of NaK and POL2 genes (but not in analysis based only on POL2), and *E. improba* based on limited amount of ddRAD data are found to be monophyletic (Figs 2b, 3b, Supplementary Data S12). Although two or three (depending on the dataset) specimens of European *E. fletcheri* (from Scotland, Sweden, and Estonia) unambiguously group together in all analyses (Figs 2b, 3b, Supplementary Data S12), they do not appear to be closely related to the single analysed North-American counterpart (Figs 2b, 3b). Within and between species divergences based on ddRAD data (*de novo* assembly) in *immersa* group partly overlap. Within species divergences (0.49–1.15%) are somewhat larger compared to *E. longicornis* group, while between species divergences are somewhat smaller (0.88–1.42%) (Table 5).

Similarly to *longicornis* group, there are some differences between NaK and POL2 phylogenies, one of which is rather well supported. *Empria immersa* is moderately supported as monophyletic according to NaK (Fig. 4a), but not according to POL2 (Fig. 4b), in which case two specimens of *E. immersa* are not separated from *E. camtschatica*, *E. improba*, and *E. plana*. Based on these two genes (altogether 4061 bp), net synonymous divergences between three species for which more than one individual was sampled were between 1.0–2.0%, therefore falling within the grey zone of speciation according to Roux et al. (2016). When only *E. immersa* and European *E. fletcheri* are considered (because North-American *E. fletcheri* did not group with European ones and *E. plana* was not monophyletic), net synonymous divergences between these species are 2.9%, falling outside the grey zone (Roux et al., 2016).

Admixture analysis with STRUCTURE at *K*=5 (the number of species based on morphology) does not support current taxonomy very well either, as only *E. immersa* and the European specimens of *E. fletcheri* are consistently supported as distinct populations from the others (Fig. 2b, Supplementary Data S11). Curiously, *E. camtschatica* and *E. improba* are supported as part of almost the same population, even though they are far apart in the ddRAD tree (Fig. 2a). Morphologically these two species could be the same, but STRUCTURE analyses with a different taxon sampling suggests that *E. camtschatica* and *E. improba* are largely separate populations (Supplementary Data S11). Interestingly, at *K*=3, STRUCTURE suggests that there are three clearly separated populations: European *E. fletcheri*, *E. immersa*, and the other species together (Fig. 5, Supplementary Data S11). North-American *E. fletcheri* is a mixture of *E. camtschatica*, *E. improba, E. plana,* and European *E. fletcheri* according to STRUCTURE analyses at *K*=2 to *K*=5 (Fig. 5).

**Fig. 5.**
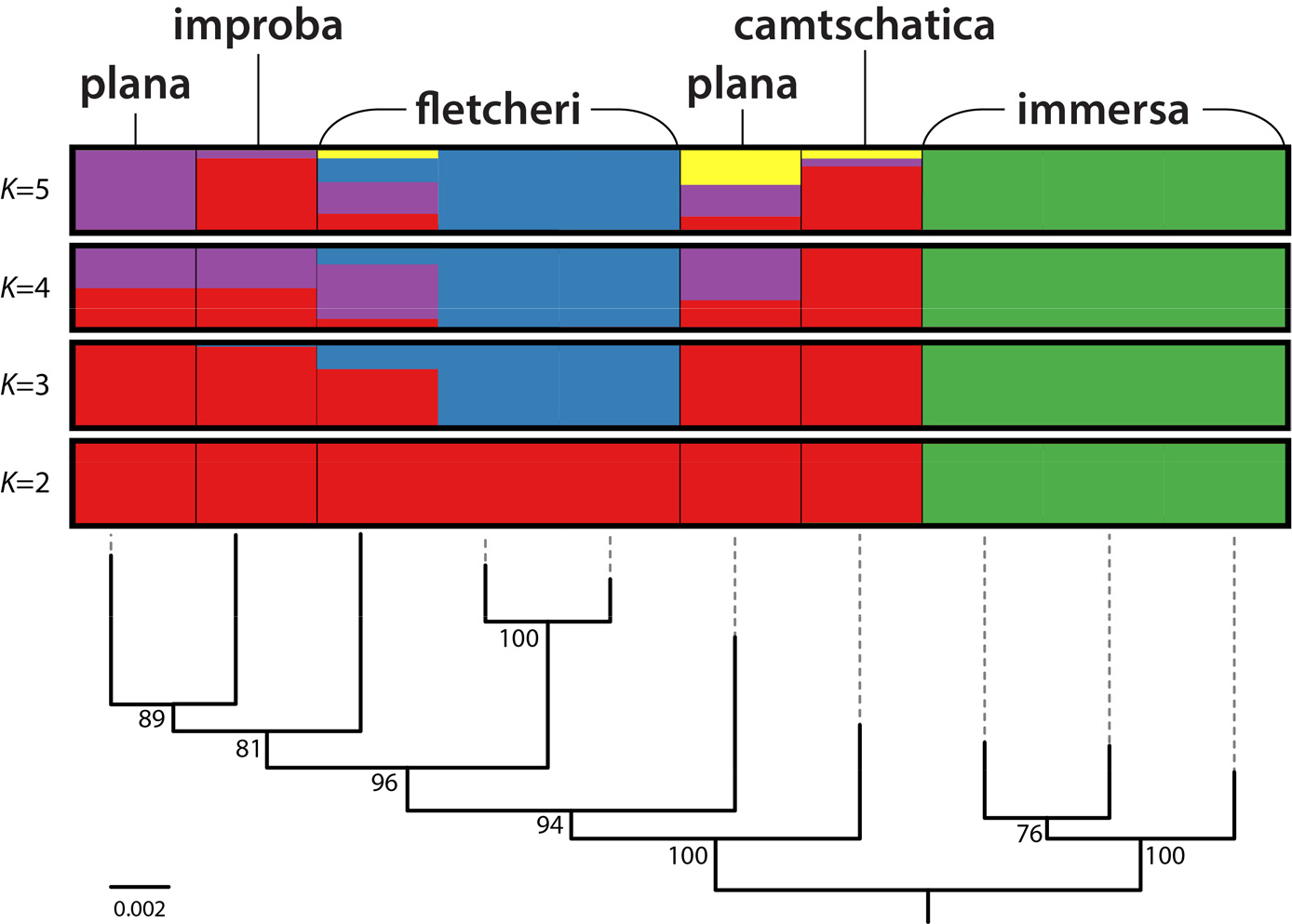
Population admixture analysis of *Empria immersa* group with Structure at *K*=3 to *K*=5 (*de novo* assembly). Maximum likelihood tree from Fig. 2 is shown below the results of Structure analyses.

The results of four-taxon D-statistic tests suggest some cases of possible introgression between different species (Table 6).

As in case of *longicornis* group, nuclear and mitochondrial trees of *immersa* group are very different from each other and monophyly of species is not supported (Fig. 2b).

## 4. Discussion

### 4.1. Cross-contamination or hybridisation?

Large amount of data generated with high-throughput sequencing methods precludes manual checking of every alignment. The main problems for these datasets are introduction of non-homologous alignment regions and contaminations, which would be easy to notice when dealing only with few genes by checking every alignment and gene tree. Because the species in our dataset are all closely related, the exclusion of non-homologous alignments is relatively easy by increasing clustering similarity threshold (up to 90% or 95% in our case) together with an addition of stringent filtering steps, which excludes all or most problematic alignments. However, in studies that pool specimens for sequencing and use single barcode or barcode combinations (but fewer unique barcodes than specimens) to link the reads to specimens after sequencing, it becomes especially difficult to detect cross-contamination when study organisms are closely related, because it might not be obvious if identical loci between specimens are due to biological or technical reasons. Nevertheless, the two species-groups of *Empria* studied here are morphologically and genetically (based on both mitochondrial and nuclear DNA) well separated (Prous, 2012) and hybridisations between them are unlikely. Therefore, by examining patterns of identical loci between the groups, it is possible to get an estimation of the level of cross-contamination in our dataset. In the dataset excluding divergent loci (19 413 loci, Supplementary Data S11), about 6% of the sequences involve identical pairs between *immersa* and *longicornis* groups (in order not to exclude any specimens, *E. tridentis* was treated as a member of *longicornis* group in these pairwise comparisons). This percentage is certainly over-estimation regarding cross-contamination as many of the loci (which are short, about 180 bp) might be too conserved to reveal differences between the groups. Better indication would be examination of the cases where one specimen is identical to any other in the wrong group while different from specimens in its own group (if present). In this case 1% of all sequences are identical to specimen(s) in the wrong group. Assuming the same cross-contamination rate within *immersa* and *longicornis* groups themselves suggests 2.4% of the sequences of the whole dataset to be affected (58% of pairwise comparisons are within *immersa* and *longicornis* groups). It is difficult to say if this is still over-estimation or instead under-estimation. Based on a more stringent criterion, e.g. considering loci present for four or more specimens from both groups (i.e. at least 8 specimens) gives cross-contamination rate of 0.6% for the whole dataset. Studies that specifically examined cross-contamination have found that barcode jumps can cause 0.3–2.5% of the sequence reads to be assigned to a wrong individual (Kircher et al., 2012; Schnell et al., 2015). Cross-contamination level of 1–2% could be considered acceptable if it affected every specimen in equal proportions, but might still lead to questionable conclusions if some specimens were affected much more than others (Valk et al., 2018). In our dataset, *E. japonica* USNM003 was a clear outlier in regard to possible cross-contamination between *immersa* and *longicornis* groups (Fig. 1a). Remarkably, when examining specific individuals that exclusively contribute identical loci to *E. japonica* USNM003, *E. immersa* DEI-GISHym80071 is the most common (85 loci out of 1015, 8.4%), while other specimens of *E. immersa* group contribute practically nothing (two other specimens contributed exclusively both only one locus) (Fig. 1b). It is highly unlikely that one specimen of *E. immersa* from Sweden and a one specimen of *E. japonica* from Hokkaido (Japan) share genes because of introgression, while the same is not seen between other specimens of the same species. If introgression was likely between *immersa* and *longicornis* groups, one would expect to see comparable levels of gene sharing in other specimens and preferably between specimens in geographic proximity (most studied specimens are from Europe). Similarly for the other *E. japonica* specimen (USNM038) from Hokkaido that we had to exclude because of possible cross-contamination (although involving specimens from the same species group), the largest contributor of identical sequences (12.3%) was specific *E. minuta* specimen (MH10-01) from Estonia, while the other *E. minuta* specimens contributed practically nothing (Fig. 1c). The second largest contributor of identical sequences (5.1%) to USNM038 was *E. loktini* (Fig. 1c), which does live in sympatry with *E. japonica*. Although species might hybridise in *E. longicornis* group (at least in the past as suggested by mitochondrial phylogeny), this high proportion of identical sequences in USNM038 specifically from *E. minuta* and *E. loktini* seems unlikely from morphological perspective (*E. japonica* can unambiguously be distinguished from those two species, but not so well from *E. longicornis* and *E. tridens*) and generally suspicious compared to most other specimens (in most cases each heterospecific individual contributed less than 2.5% of identical sequences). Additionally, PCR and Sanger sequencing of two loci that were suspected to be cross-contamination in USNM038 contradicted the ddRAD data: sequences of two or three *E. japonica* specimens were identical to each other while different from all the other sequenced specimens (Supplementary Data S7). Although our dataset overall might not be affected by higher level of cross-contamination than usual (e.g. Schnell et al., 2015), the relatively high level of unequal recovery of loci (difference more than 10 times) among the specimens might explain why some of them were proportionally more affected than others (Valk et al., 2018). Among the specimens retained for the analyses, both *E. japonica* specimens were at the lower end of the number of loci recovered. If more loci had been recovered from both *E. japonica* specimens, these might have diluted the cross-contamination and we might not have realised that possibility. It is likely that some (perhaps even most) other specimens in our dataset have also been affected by cross-contamination (including among conspecific individuals, which we could not detect) to some degree, which weakens our biological conclusions for the studied species groups. Nevertheless, except in case of *E. japonica*, the results based ddRAD data seem plausible in light of morphological studies (Prous, 2012; Prous et al., 2014, 2011b) and Sanger sequencing (Fig. 4), but should be re-examined in future studies that better control for cross-contamination. Although more expensive, we suggest that whenever possible, additional replications should be done with different combinations of pooled specimens, pooling only distantly related species, sequencing every specimen separately, or adding specimen specific barcodes to both ends of the DNA fragments (not just different combinations of limited number of barcodes).

Even if 1–2% is considered an acceptable level of erroneous sequences (because of non-homology and/or contaminations) in phylogenomic datasets, this can still significantly affect downstream analyses. It is likely that even smaller proportion of errors could be detrimental to reconstructing rapid speciation events if the small amount signal is swamped be larger amount of error. On the other hand, resolving rapid speciation events would be among the questions to which large datasets could give the largest contribution (uncontroversial clades can be reliably reconstructed already based on small number of genes) and therefore data quality is essential before deciding among alternative phylogenetic hypotheses. Unfortunately, the bioinformatic tools are not yet reliable enough to completely remove manual interventions in phylogenomic datasets (Philippe et al., 2017; Simion et al., 2017).

### 4.2. Causes of mitonuclear discordance

In both studied species groups, mitochondrial phylogeny is very different from nuclear phylogeny (Fig. 2). Particularly striking is the non-monophyly of all or most species according to mitochondrial DNA, while there is little incongruence between nuclear and morphological evidence (Figs 2–4; Prous, 2012; Prous et al., 2011b). Strong mitonuclear discordance has been observed in other animal groups and usually interpreted as evidence of mitochondrial introgression (e.g. Bronstein et al., 2016; Papakostas et al., 2016; Tang et al., 2012), which does seem to be the most likely explanation (Bonnet et al., 2017; Sloan et al., 2017), although it may also result from incomplete lineage sorting (Funk and Omland, 2003). Recently, Patten et al. (2015) found based on theoretical modelling that haplodiploid species may be especially prone to biased mitochondrial introgression, which could explain widespread mitonuclear discordance in several other species rich sawfly groups besides *Empria*, like *Neodiprion* (Linnen and Farrell, 2008, 2007) and *Pristiphora* (Prous et al., 2017). Nevertheless, haplodiploidy is probably not the only reason to promote widespread mitonuclear discordance, because in many other (most?) Hymenoptera, mitochondrial barcoding seems to work relatively well for species identification (Derocles et al., 2012; Klopfstein, 2014; Schmidt et al., 2015). One other factor that might influence rate of mitochondrial introgression is its mutation rate, lower rates making introgression more probable than higher rates, latter of which should more likely lead to compensatory co-evolution and mitonuclear incompatibilities (see Table 3 in Sloan et al., 2017). As mitochondrial genomes of basal hymenopterans do evolve significantly slower compared to Apocrita (particularly Xyeloidea, Pamphilioidea, and Tenthredionoidea; Niu et al., 2019; Tang et al., 2019), the combination of haplodiploidy and slow rate of mitochondrial evolution might better explain widespread mitonuclear discordance in some (many?) species rich groups of sawflies rather than just haplodiploidy. There is evidence that in parasitic lineages mitochondrial evolution tends to be much faster than non-parasitic lineages (Pentinsaari et al., 2016), explaining perhaps faster evolution of mtDNA in Apocrita which are ancestrally parasitic (and most species still are). It could be then that in most Hymenoptera (Apocrita), COI barcoding might be reliable for species identification despite of haplodiploidy.

### 4.3. Taxonomy of E. longicornis group

Since the revision of the species group (Prous et al., 2011b), two additional putative species have been found, and a third already described species (*E. montana*) is here for the first time recognised as member of *longicornis* group (Supplementary Data S9 and S10), bringing the total number of species to 12. The male specimen USNM2051678-040 from Hokkaido is so far the only known representative of a putative species “sp11” (sp.1 in Prous et al., 2011a). Because there has been very little sampling of sawflies in arctic habitats above treeline in Japan (and more generally outside Europe), the sp11 might normally be restricted to arctic habitats like *E. alpina* (sister species of sp11 according to some ddRAD trees: Fig. 2), although the single known specimen was collected in a Malaise trap well below treeline (at 1000 m, about 15 km East of Mount Asahi, the highest point on Hokkaido). We have studied several specimens of the second putative species, sp14 (male MH11-01 and female DEI-GISHym15231 reported here) collected in the Pyrenees and the Alps, in most cases below treeline, but at higher altitudes than 1500 m. Morphologically, the only rather clear indication that sp14 might be a different species from *E. alpina*, is the different structure of female ovipositor. Male penis valves do not seem to be different in *E. alpina* and sp14, but there is variation in the length of antennae. Confusingly, though, *E. alpina* in the Alps have distinctly longer antenna (both male and female) from the specimens in the northern Fennoscandia, the latter of which have antenna more similar to sp14 in the Alps. Although several additional female and possibly male specimens of sp14 are available, we refrain from describing new species, because more studies are required to more reliably resolve taxonomy of *E. alpina* and sp14, and to associate males and females. Prous et al. (2011b) noted that there might be an additional species amongst *E. tridens*, based on differences in larval colour pattern and diverging ITS sequences, but the data was too limited to decide this (no differences in adult morphology were detected). One of the specimens analysed here is the larva with diverging ITS sequence and a different colour pattern (TUZ615027, 06-05a in Prous et al., 2011b), but our ddRAD data and Sanger sequenced genes (Figs 2–4) do not clearly indicate that it should be treated as a different species. When excluding *E. alpina*, for which we lack sufficient amount of fresh material, *E. tridens* is known to be geographically the most widely distributed (from Europe to Hokkaido) among the remaining species (Prous et al., 2011a), which might explain the higher genetic diversity of this taxon, rather than indicating presence of an additional species. For other species with more than one individual sampled (*E. alector, E. basalis, E. longicornis, E. minuta*) our results (Figs 2–4) agree perfectly with current taxonomy (Prous et al., 2011b) and do not suggest the presences of additional species. In case of *E. japonica*, Sanger data unambiguously supports validity of this species (Fig. 4) and it is found to be monophyletic also based on ddRAD data when *immersa* group (which might be source of cross-contamination in one *E. japonica* specimen) is excluded (Supplementary Data S12). Validity of the species is also supported by net synonymous divergences among species based on three protein coding genes (NaK, POL2, and ZC3H14), which in most cases (2.3–9.4%) fall outside the grey zone of speciation according to Roux et al. (2016). Only divergence (1.5%) falling within the grey zone is between *E. basalis* and *E. tridens* (according to Roux et al., 2016 the grey zone of speciation falls within 0.5%– 2.0% of net synonymous divergence between species). Limitation of our dataset regarding species delimitation could be lack of sampling of specimens of same species from wider area than Europe, although at least some of them are known from West or Central Europe to East Asia (Prous et al., 2011b; Taeger et al., 2018). *Empria alector, E. basalis, E. longicornis,* and *E. minuta* were well supported as monophyletic (Figs 2–4), but this needs to be tested by sampling additional specimens from West and East Siberia.

### 4.4. Taxonomy of E. immersa group

Based on adult morphology there should be at least two species within *E. immersa* group, *E. fletcheri* (feeding on *Betula*) and the others (feeding on *Salix*). The saw (ovipositor) of *E. fletcheri* is clearly different from *Salix* feeding species. There is quite clear difference also in the structure of tarsal claws, *E. fletcheri* has small subapical tooth, while in the others it is distinctly longer (except in one possibly additional species not sampled here, *E. asiatica* that has a saw indistinguishable from *E. camtschatica* and *E. improba*). All of the *Salix* feeding species are very similar in adult morphology and might as well belong to same species. However, among the *Salix* feeders, genetic data suggests separation of *E. immersa* from the others (Figs 2–5), which might be supported by differences in colouration of larvae (based on unpublished *ex-ovo* rearings of *E. immersa* and *E. camtschatica*) and some morphological differences in the adults (Prous et al., 2014). Admixture analysis with Structure at *K*=3 (Fig. 5) suggest that *Empria camtschatica*, *E. plana*, and *E. improba* might belong together, which does not seem unlikely based on morphology (in this case the species name to be applied would be *E. improba* (Cresson, 1880), as the oldest), although more sampling throughout Asia and North-America for genetic studies would be preferably to decide among competing scenarios. Another issue requiring more attention is the apparently clear genetic separation of European and North-American *E. fletcheri*, which from a morphological perspective clearly belong together. Considering that habitats of *E. fletcheri* (bogs in boreal forests and tundra) in Eurasia and North-America were at least partly connected about 10 000 years ago, this species might well be Holarctic in distribution (in Eurasia the eastern-most specimens confirmed so far are from Irkutsk region). Unfortunately, our data does not currently allow disentangling effects of wide geographic separation and biological barriers to gene-flow in *immersa* and *longicornis* groups. In case of *longicornis* group, where taxonomy of most species is better resolved than in *immersa* group, none of the species analysed here include individuals collected within an area larger than Europe, although at least some of the species reach East Asia (Prous et al., 2011b; Taeger et al., 2018). Because species within *longicornis* and *immersa* groups are closely related (within and between-species genetic distances are quite similar, Table 5) conspecific samples analysed from much larger area than Europe might significantly complicate species delimitation based on genetic data. To test this, additional sampling from Central and Eastern parts of Asia should be analysed. In case of *E. fletcheri*, detection of two distinct genetic lineages living in sympatry in East-Asia or North-America would be a strong indication for additional species, but our data is currently insufficient do decide this (for example *E. plana* from Sweden and Hokkaido are also genetically far apart: Figs 2–4).

## Acknowledgements

We thank Laura Törmälä for her efficient work in the biology laboratory at University of Oulu. We also wish to acknowledge CSC – IT Centre for Science, Finland for computational resources.

Matthias Hoffman (ZALF, Müncheberg) and Riina Klais (Tartu) helped with an R script. Funding: This work was supported by Finnish Academy grant #277984 to MM.

## Appendix A. Supplementary material

Supplementary data to this article can be found online at https://doi.org/10.1016/…

## Supplementary Data S1–S12

**Supplementary Data S1.**
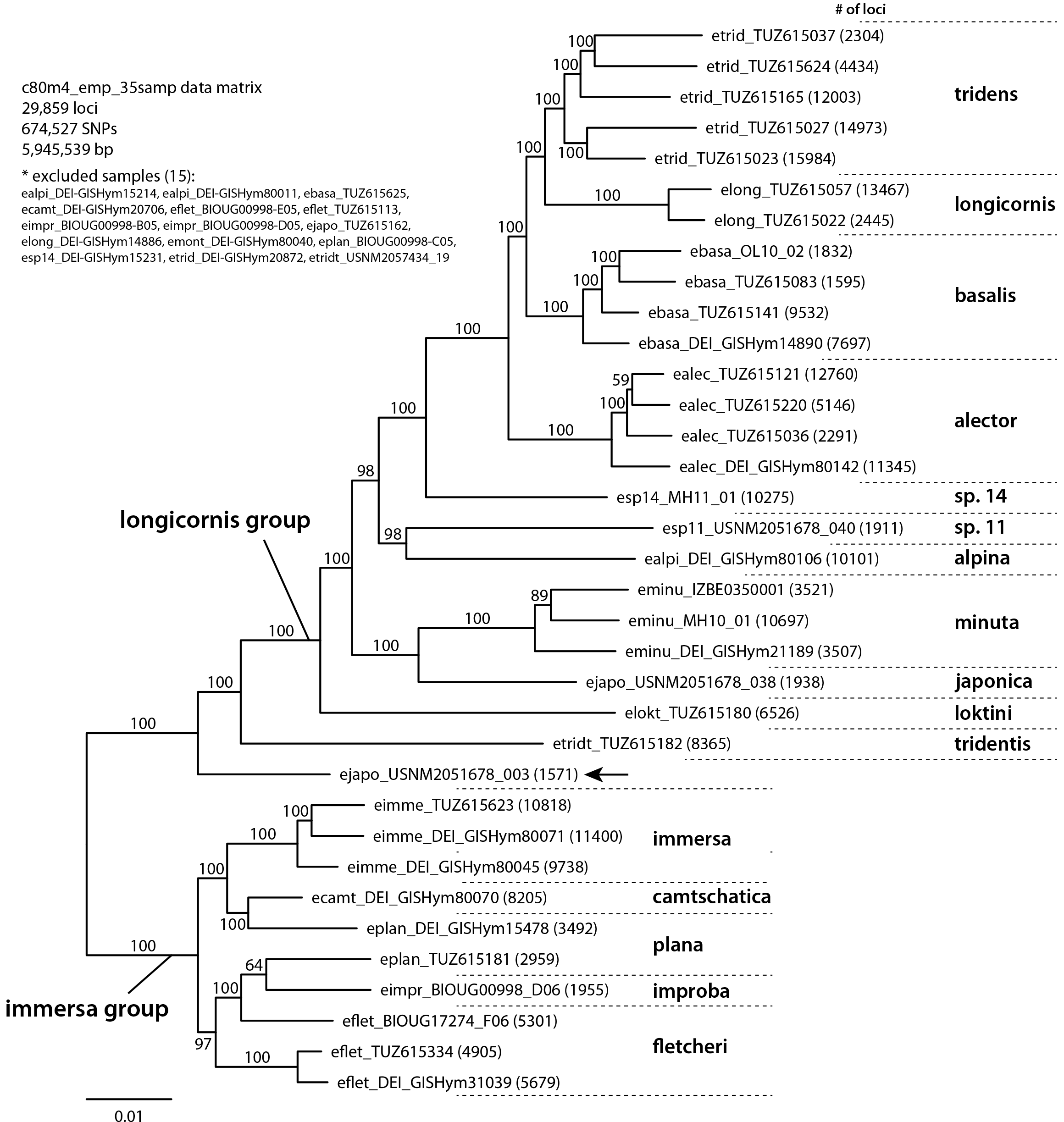
ddRAD ML tree_Empria

**Supplementary Data S2.**
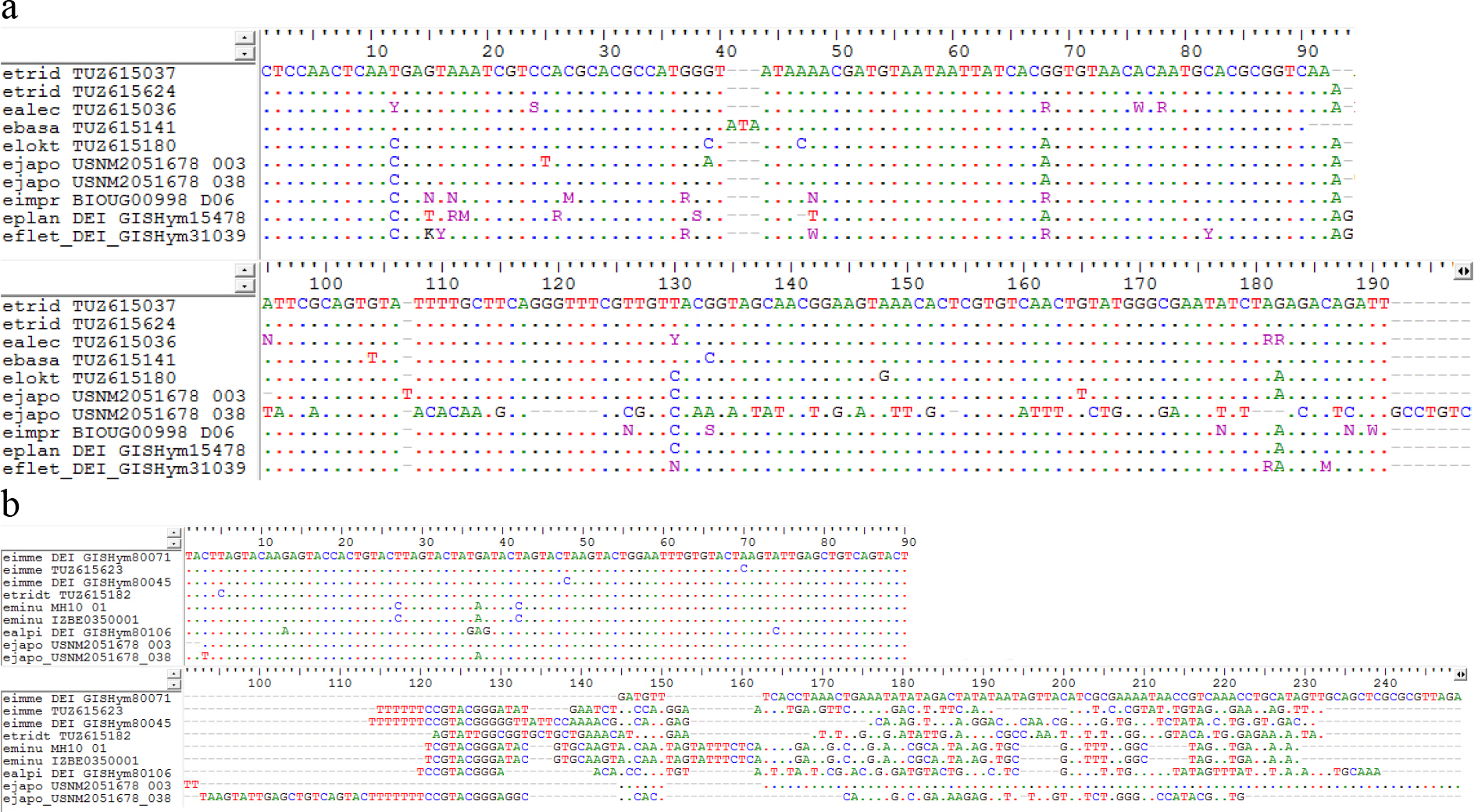

Two examples of problematic loci (possibly due to mis-association of paired-end reads) from the initial dataset with 29 859 loci (*de novo* assembly at clustering threshold of 80% similarity; Supplementary Data S1). In the first example (a) *Empria japonica* USNM2051678_038 is unalignable compared to others in the second half of the locus. In the second example (b) most specimens are very different from each other in the second half of the locus.

**Supplementary Data S3.**
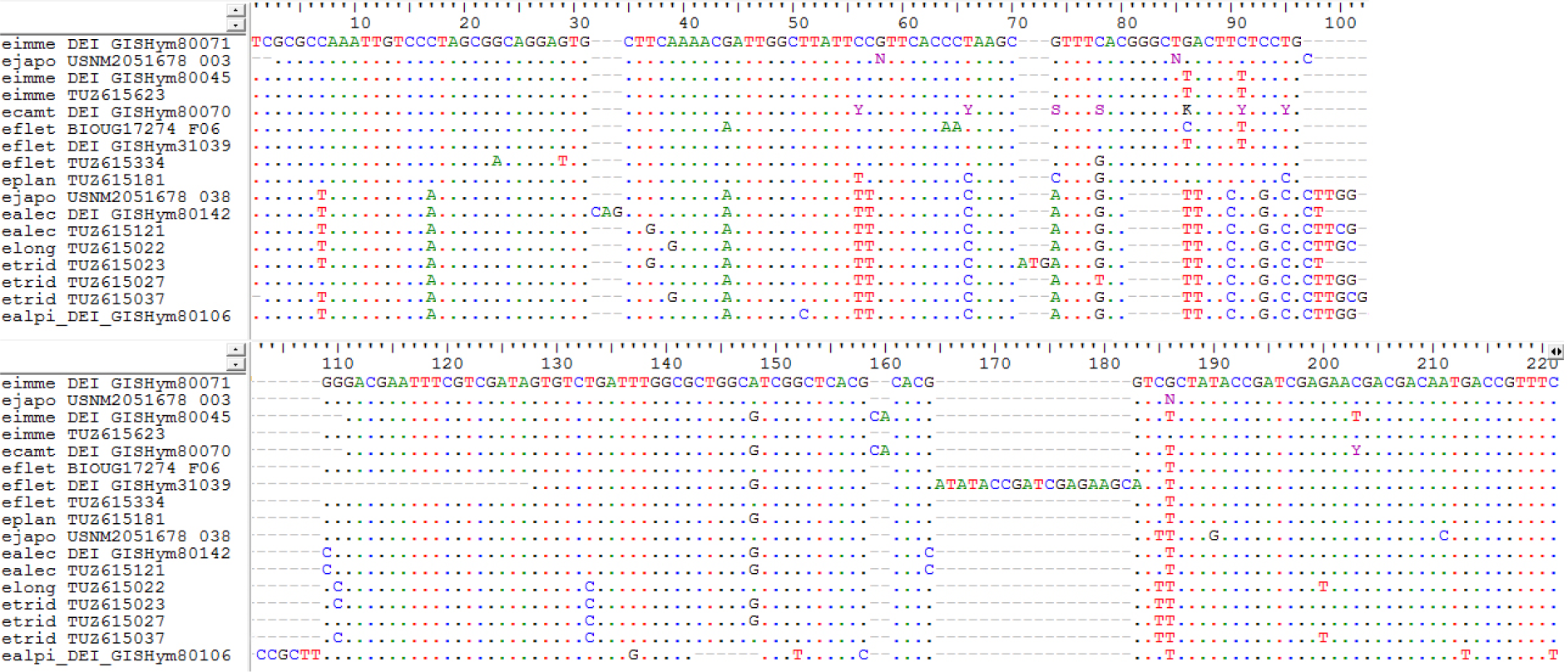

An example of locus from the initial dataset with 29 859 loci (*de novo* assembly at clustering threshold of 80% similarity; Supplementary Data S1) where *Empria japonica* USNM2051678_003 is identical (except for one indel) to *E. immersa* DEI-GISHym80071 while clearly different from other specimens, especially from specimens of *longicornis* group. The one additional “C” in the middle of USNM2051678_003 compared to DEI-GISHym80071 could be due to differences in lengths of quality trimmed paired-end reads.

**Supplementary Data S4.**
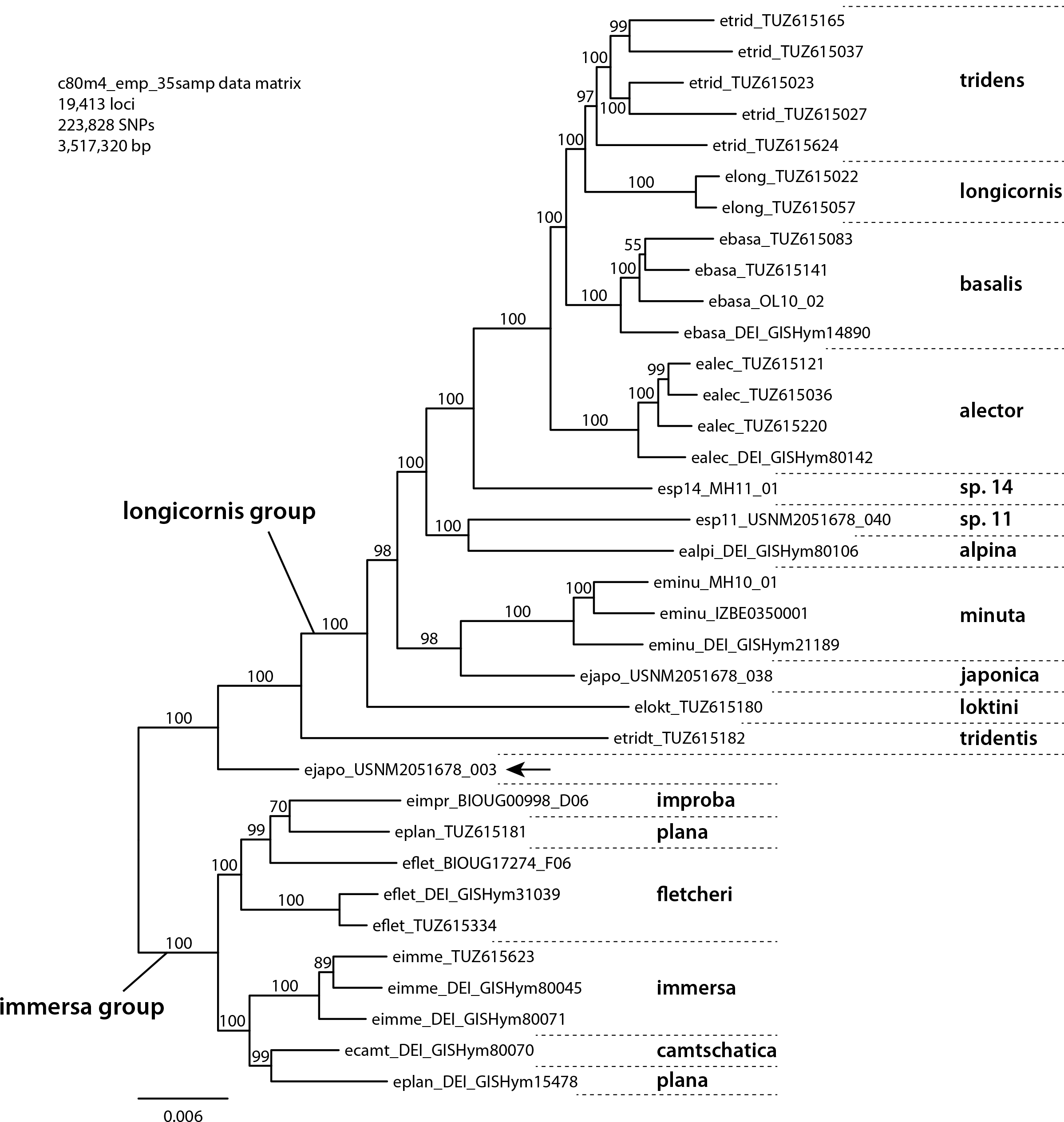

Next 20 pages show results of loci counts that are identical between *Empria longicornis* and *E. immersa* groups and among all specimens (*de novo* assembly, 19 413 loci; Supplementary Data S4). First two graphs show percent of loci in every *longicornis* (including *E. tridentis*) and *immersa* group specimen (X-axis) that are identical to any specimen in *immersa* or *longicorns* group, respectively, while different from other specimens in its own group (if present). The following graphs on 18 pages show for every specimen the percent of loci and normalised percent of loci that are identical to a particular specimen (X-axis) while different from all others in the dataset. Normalised numbers of loci for specimens were calculated as half of the maximum number of loci divided by the number of loci of that specimen. # -number of loci present for that specimen.

**Supplementary Data S5.**
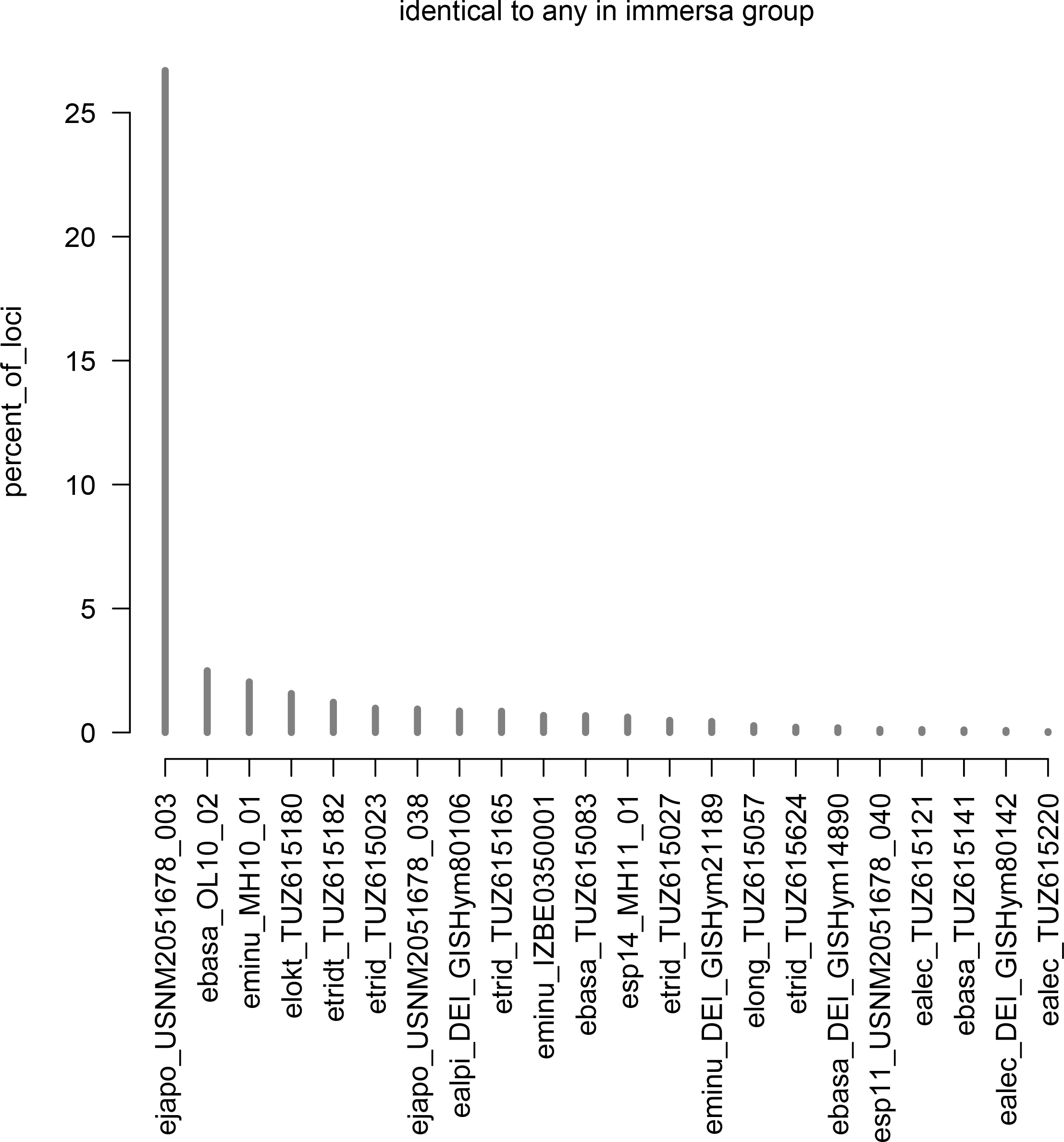

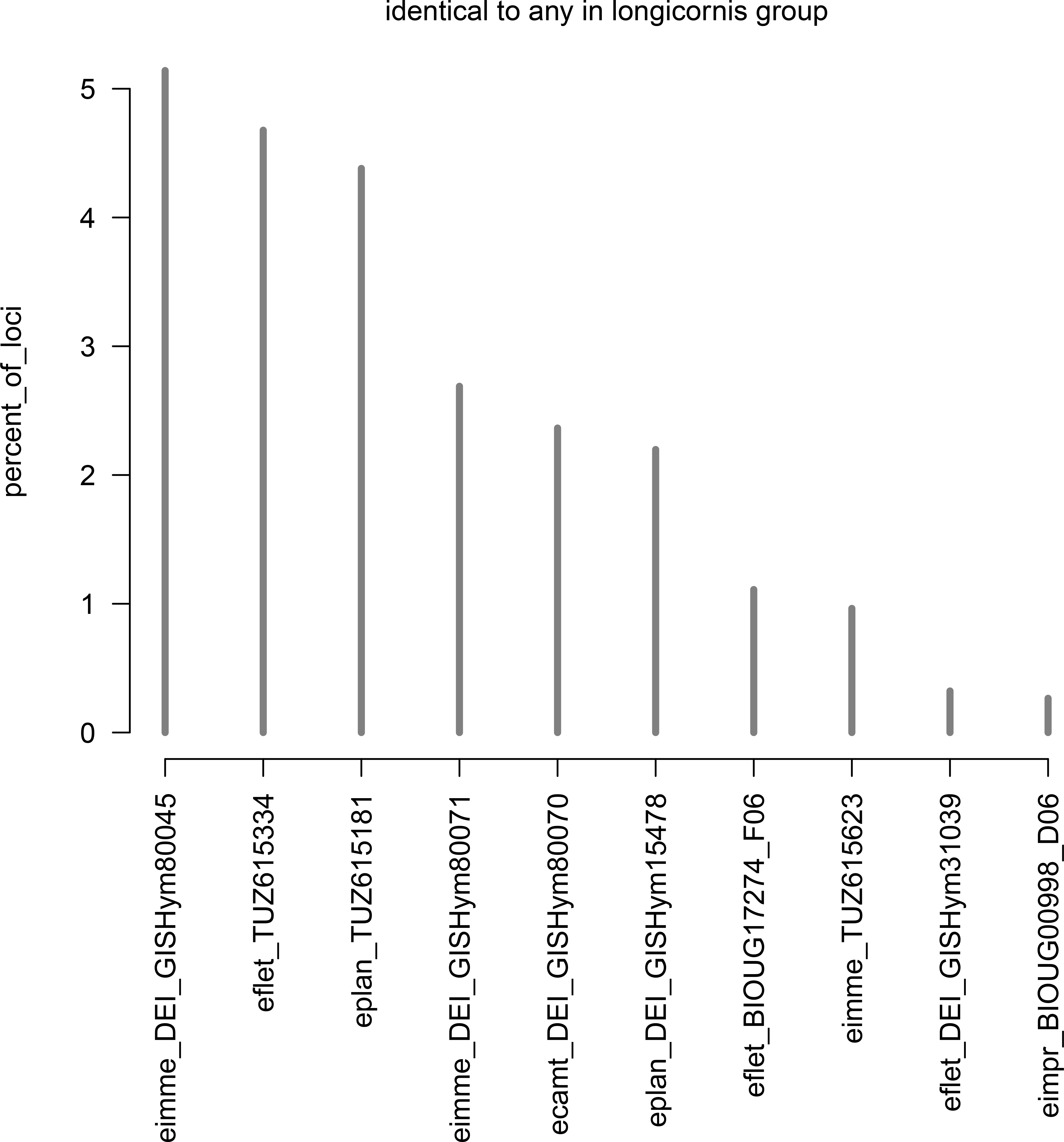

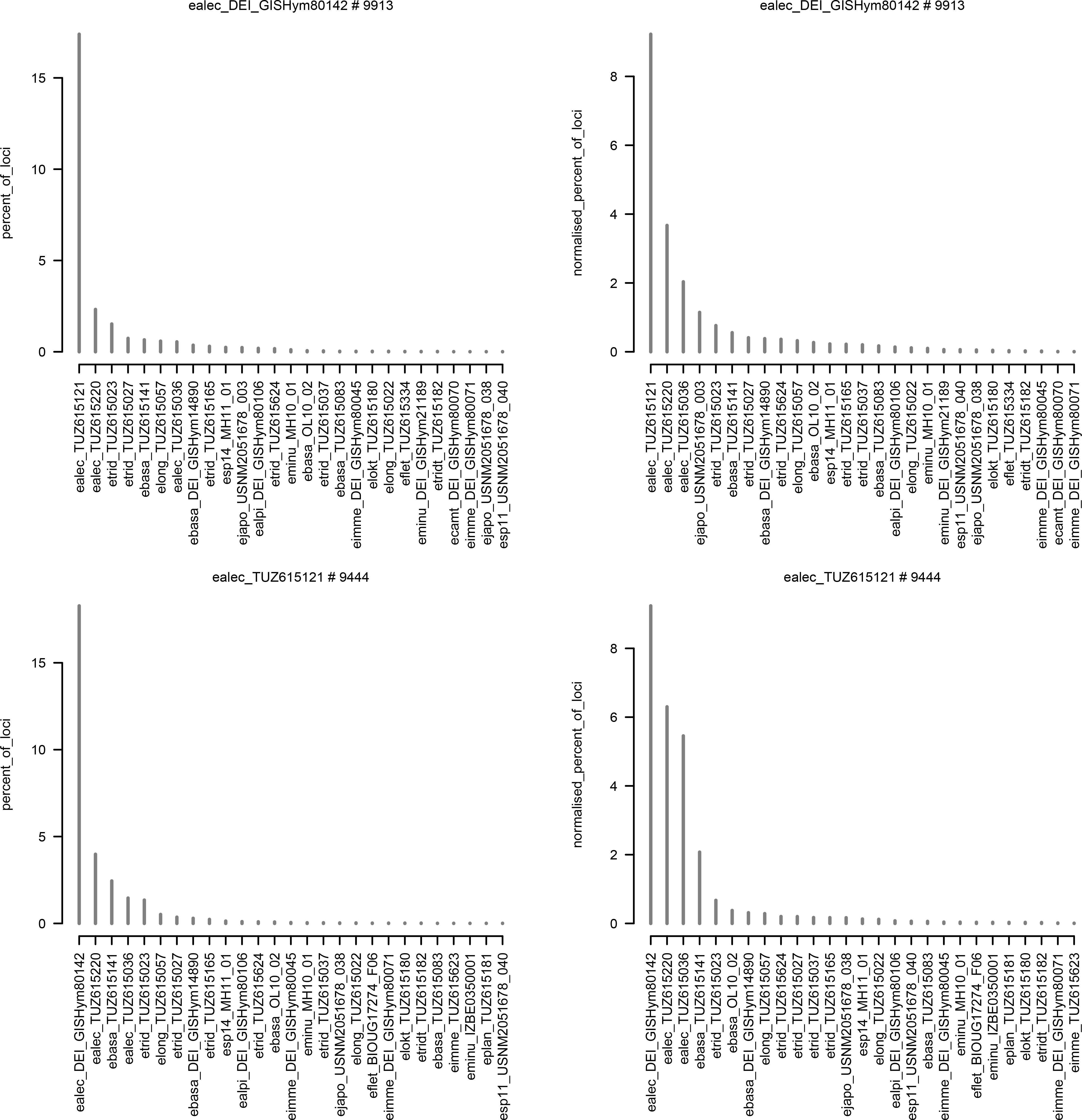

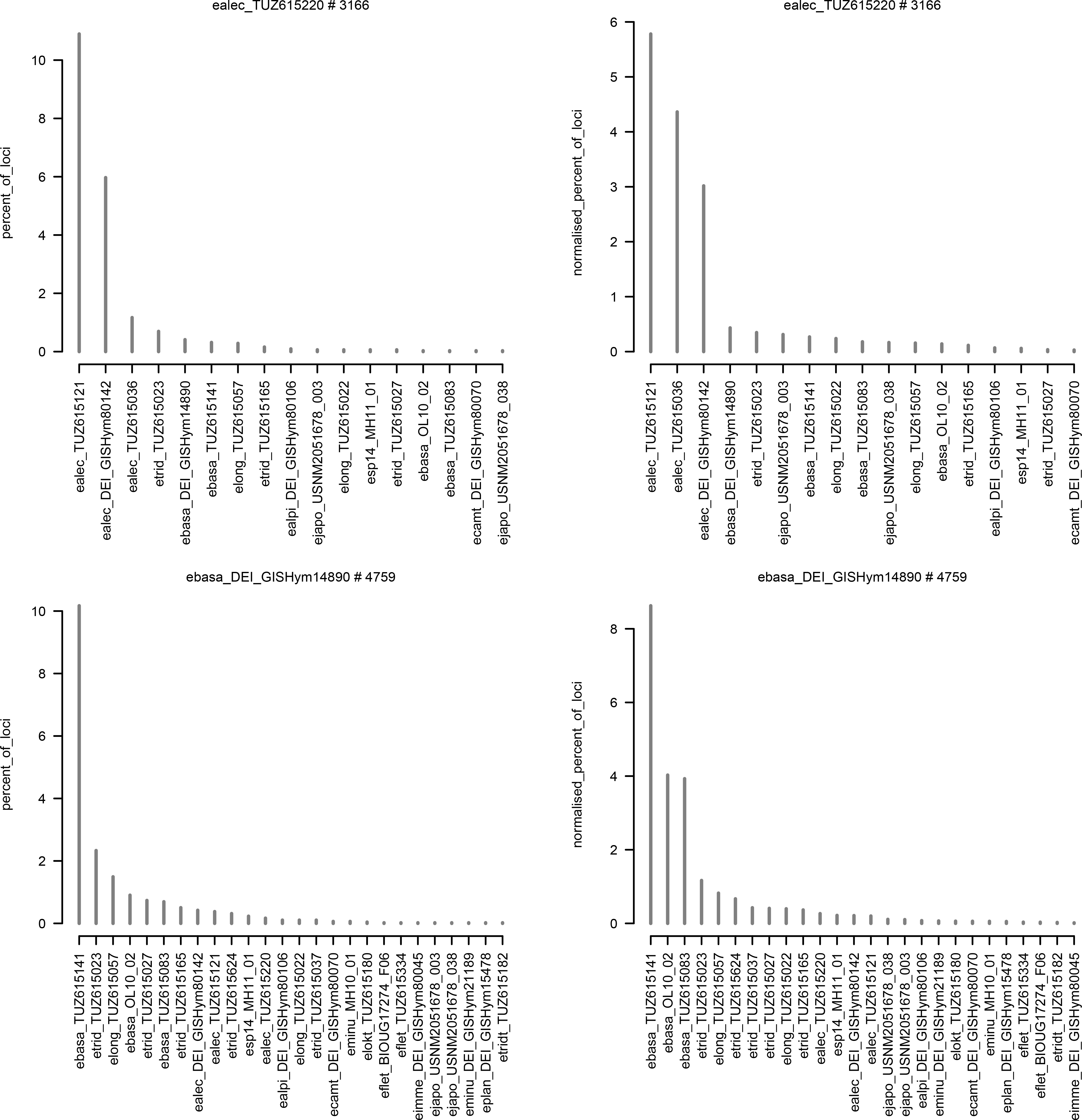

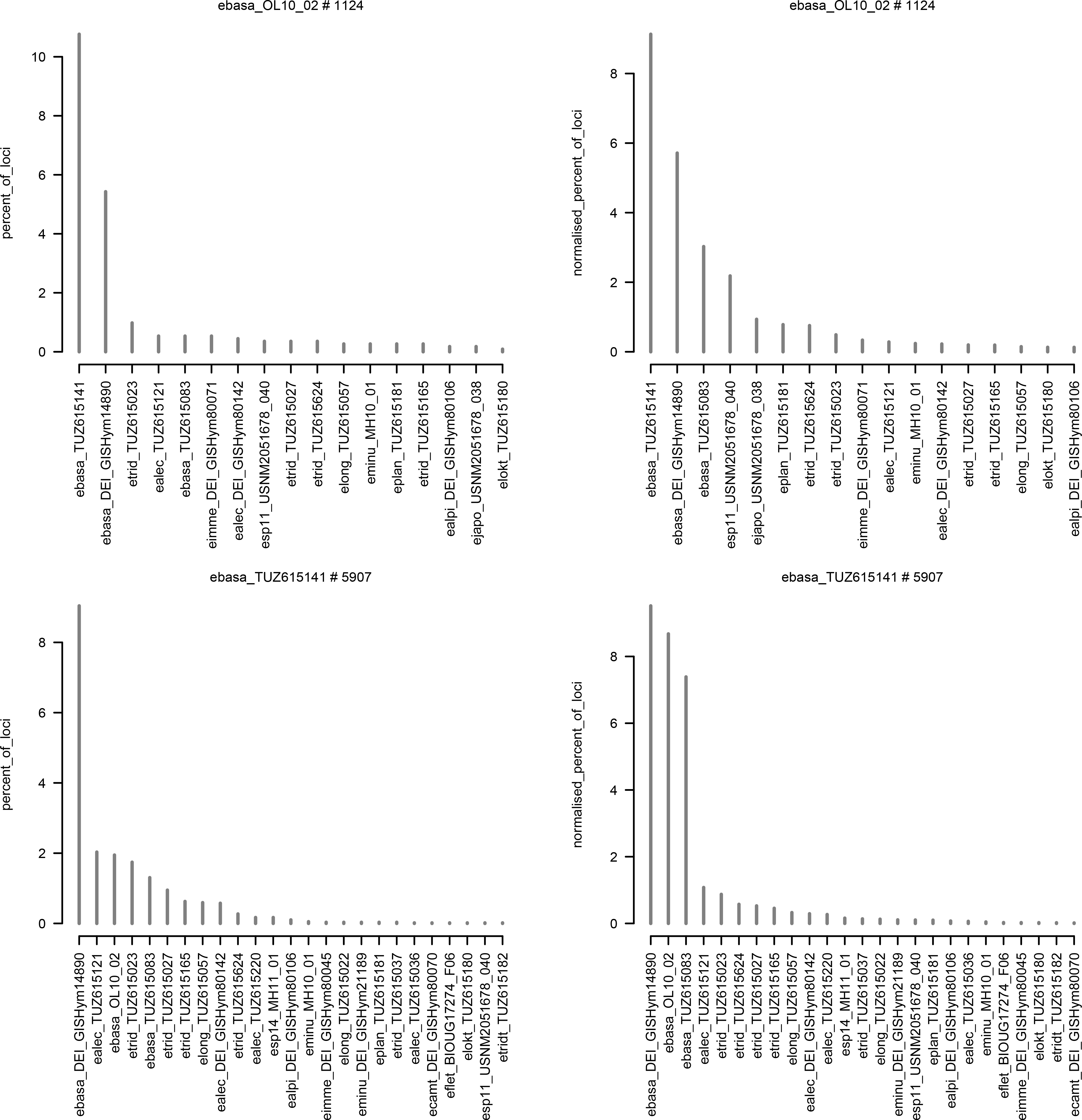

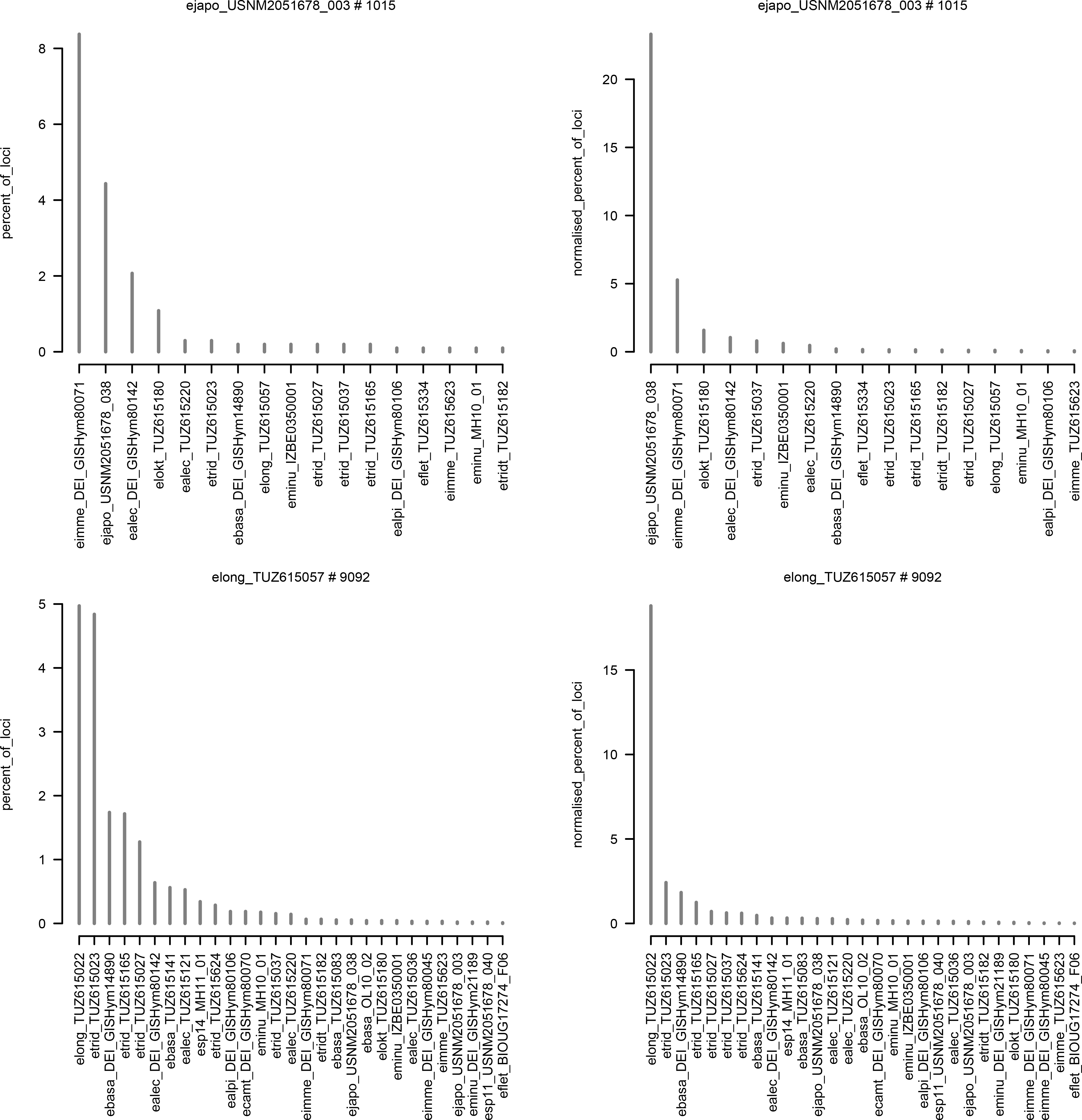

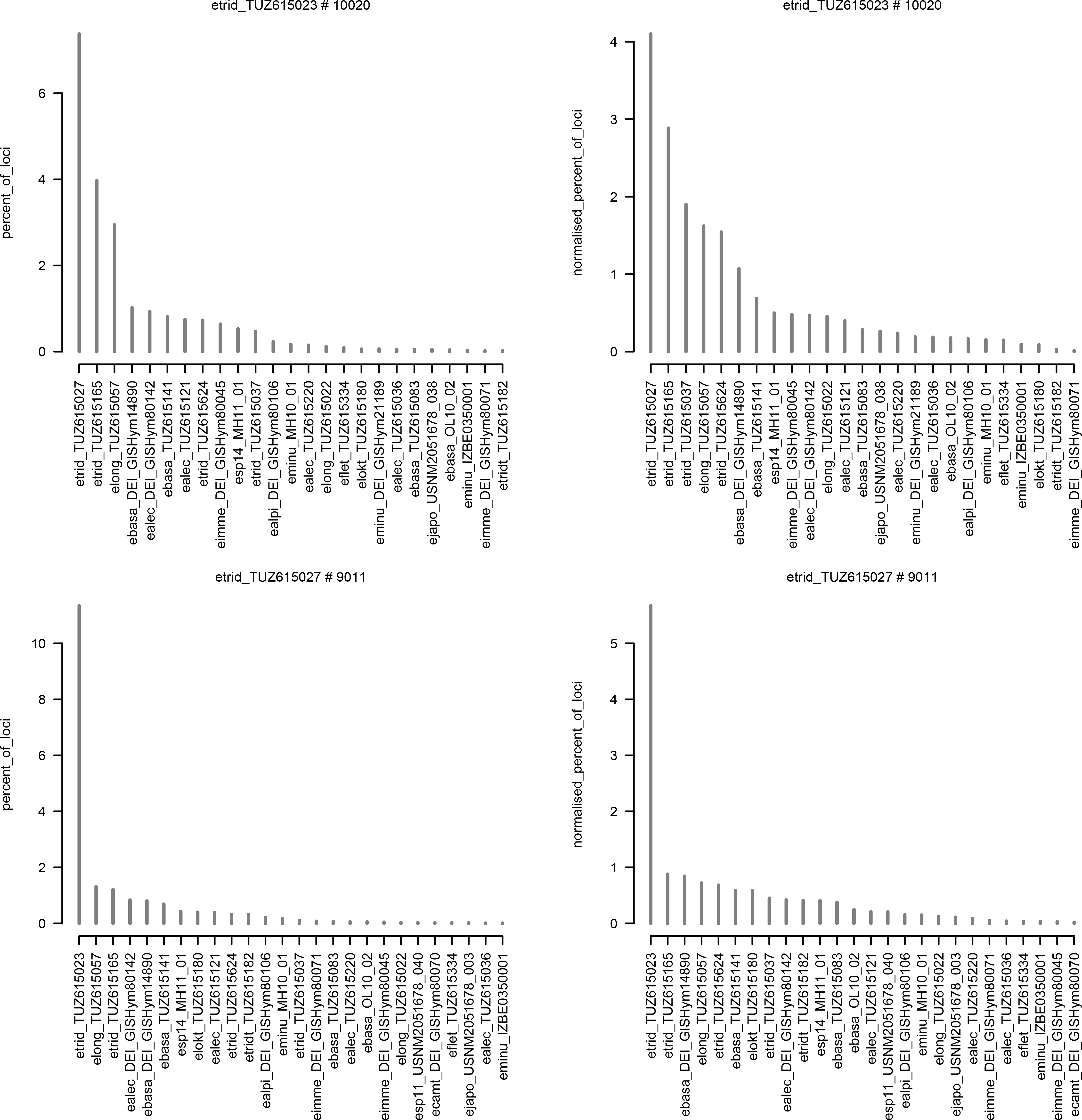

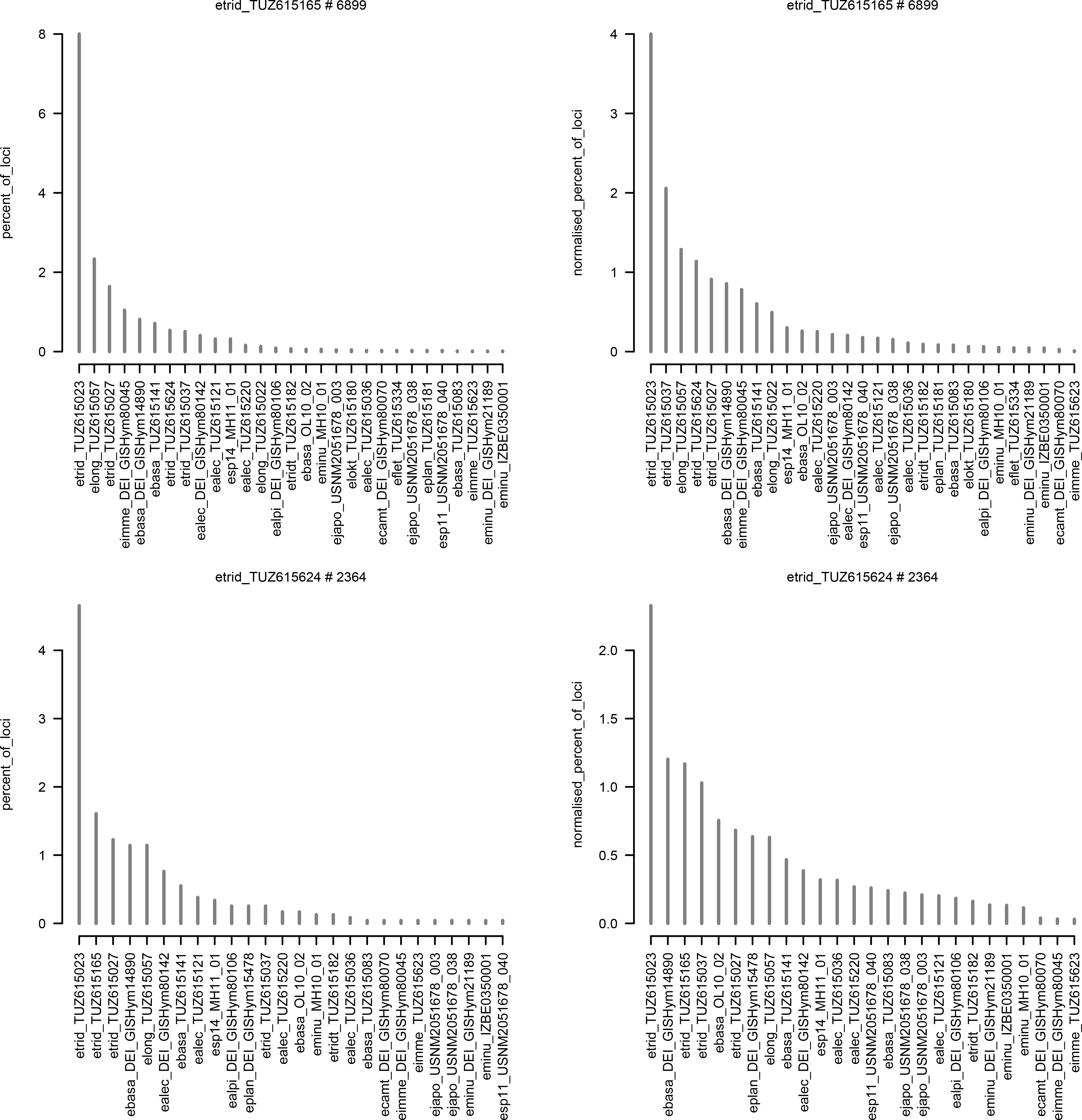

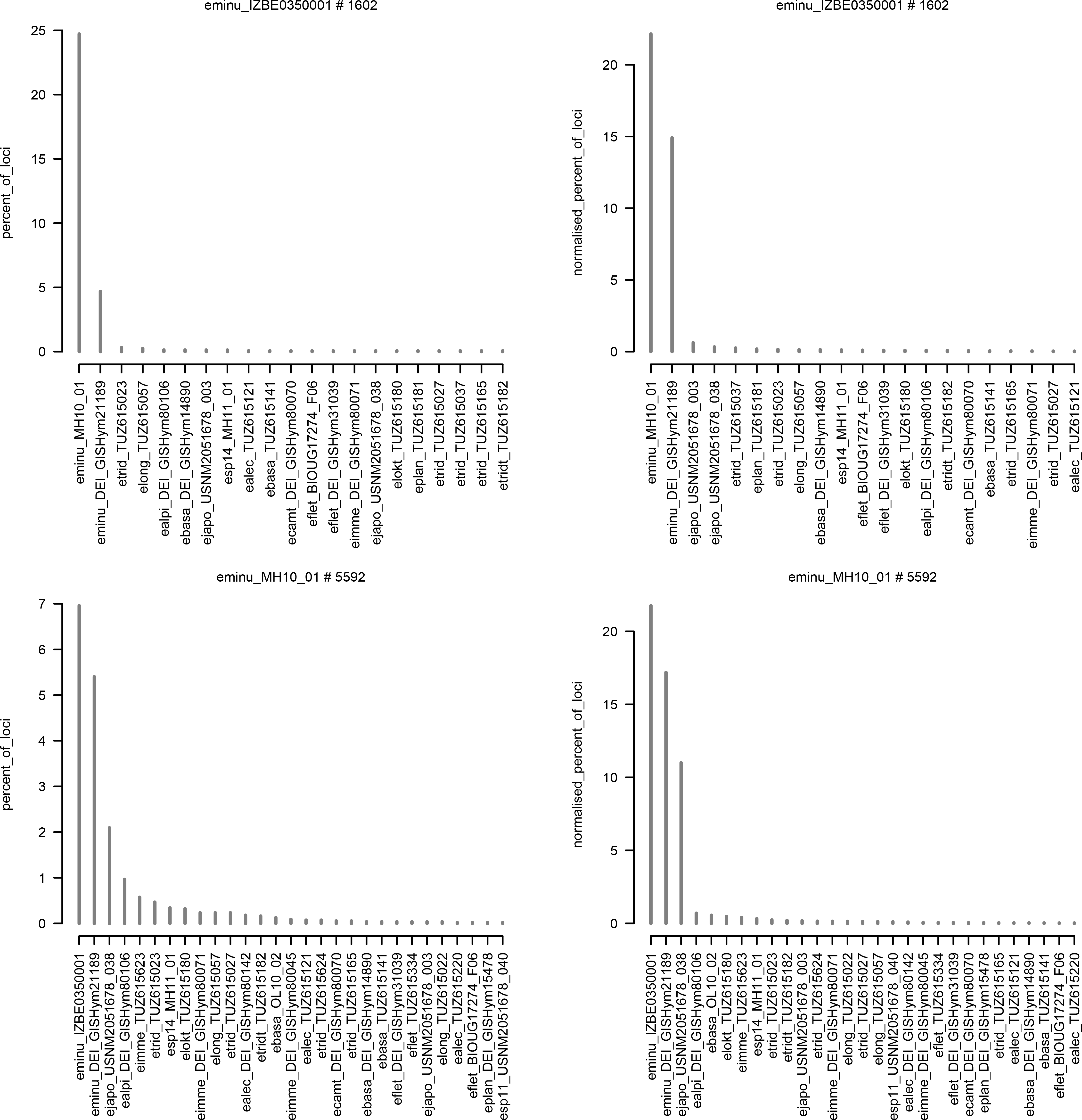

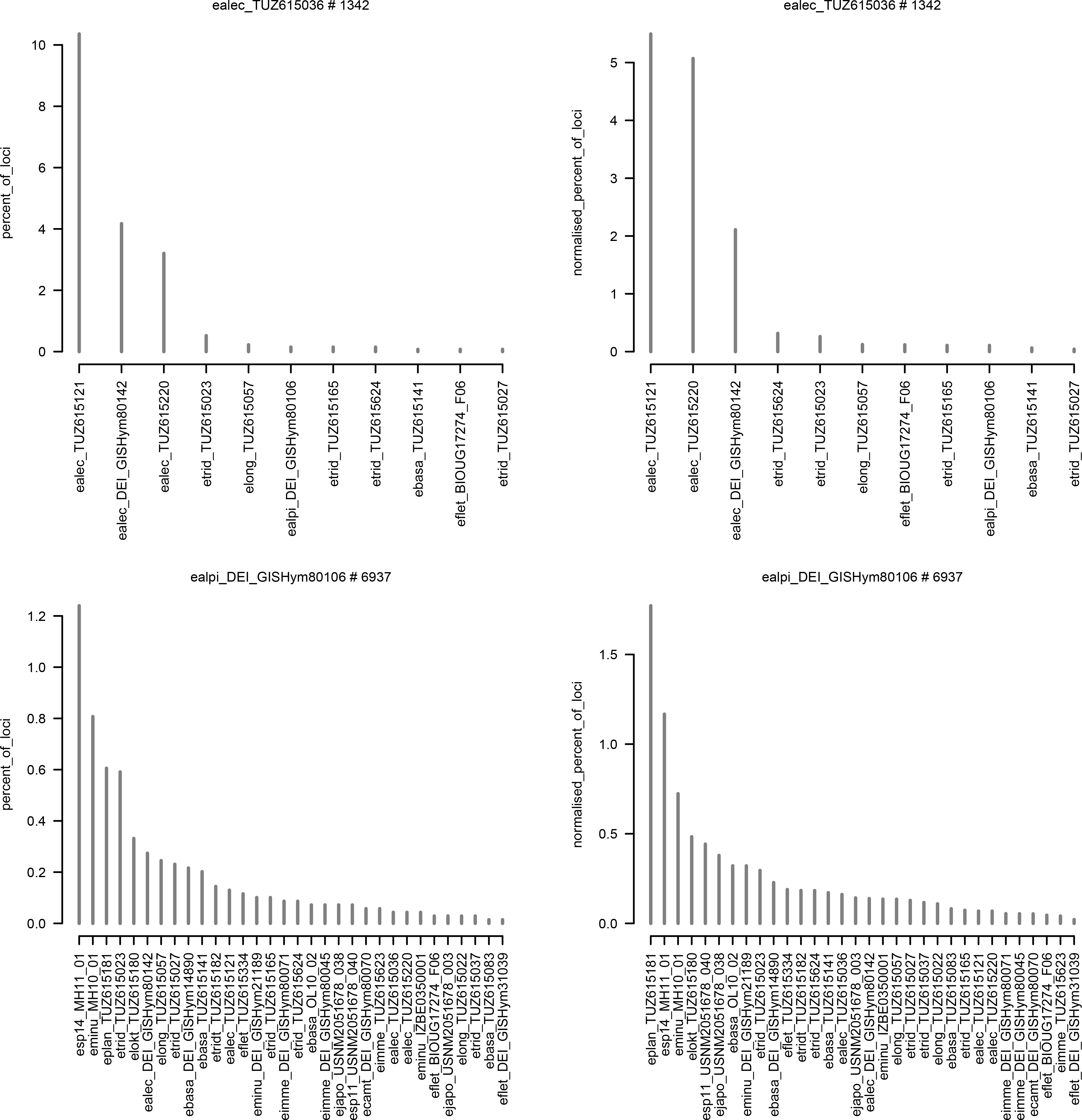

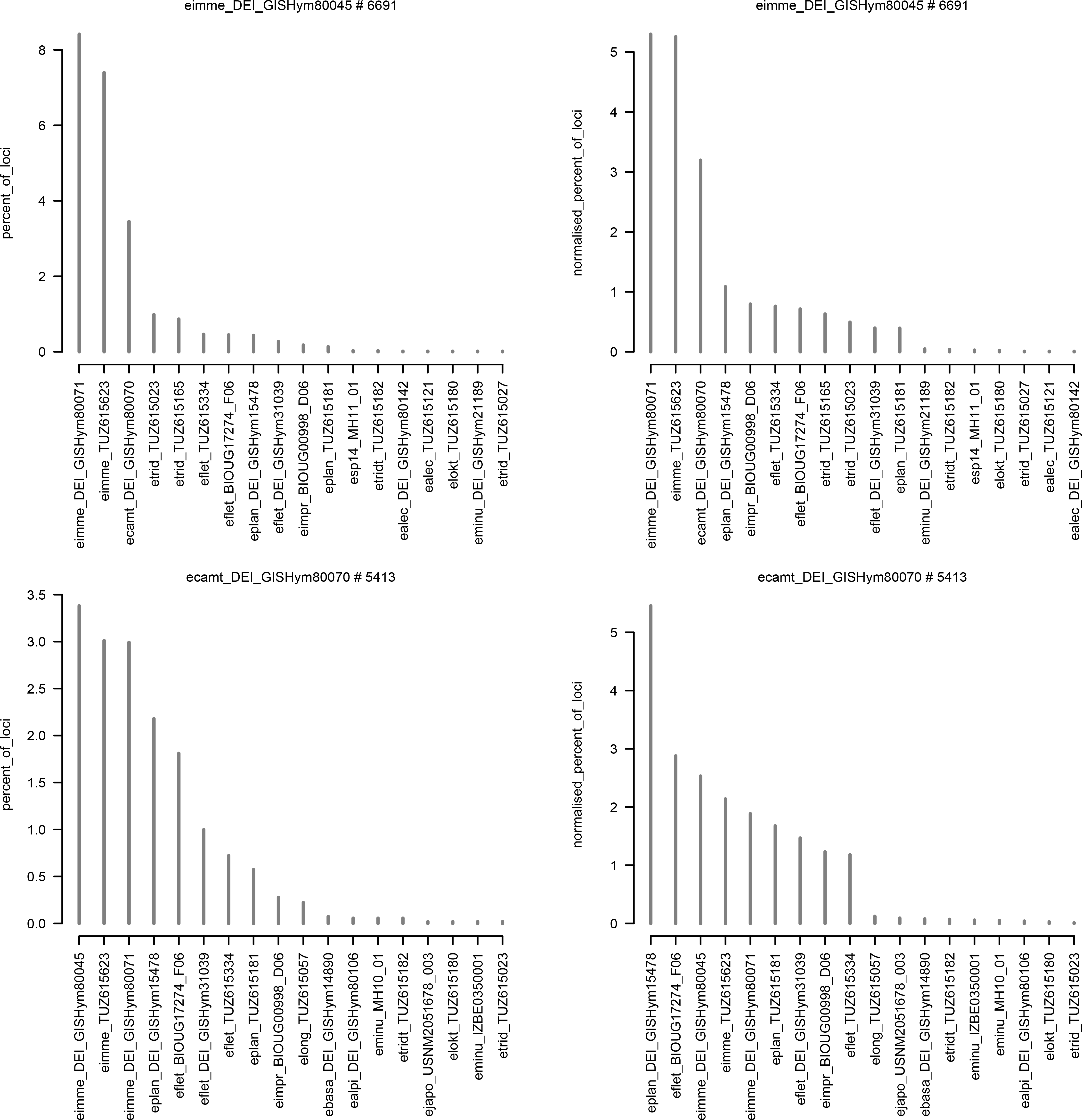

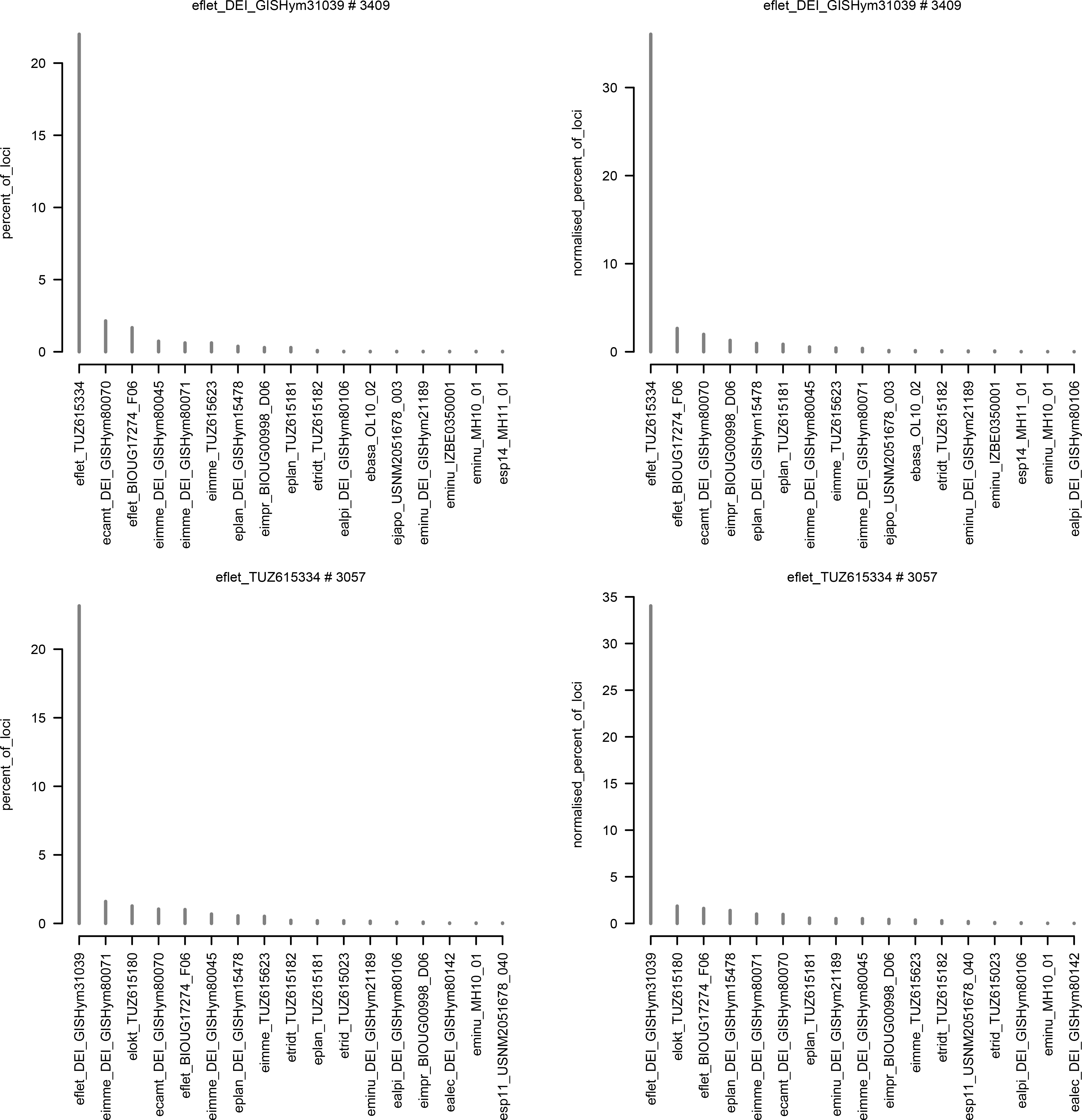

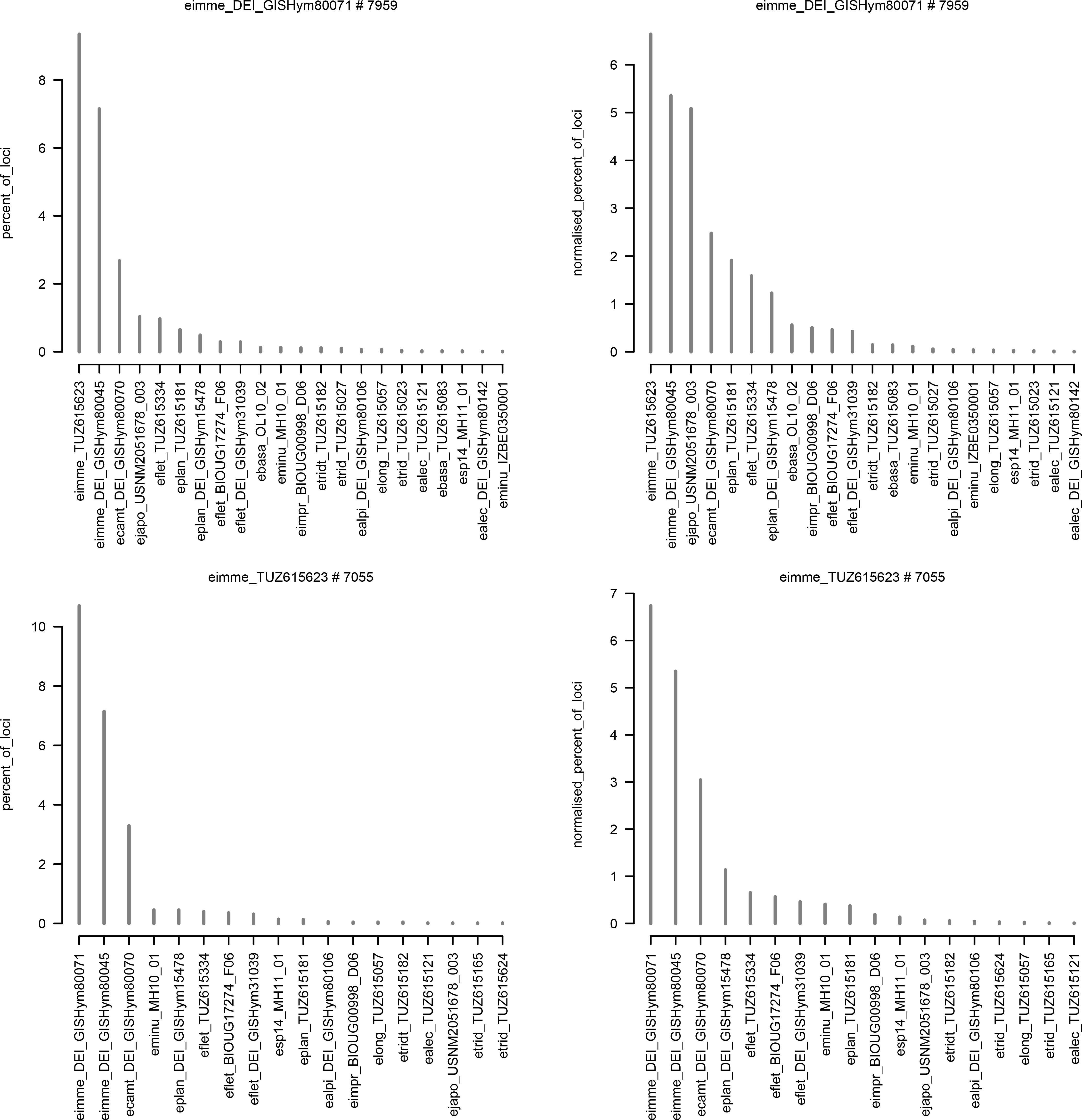

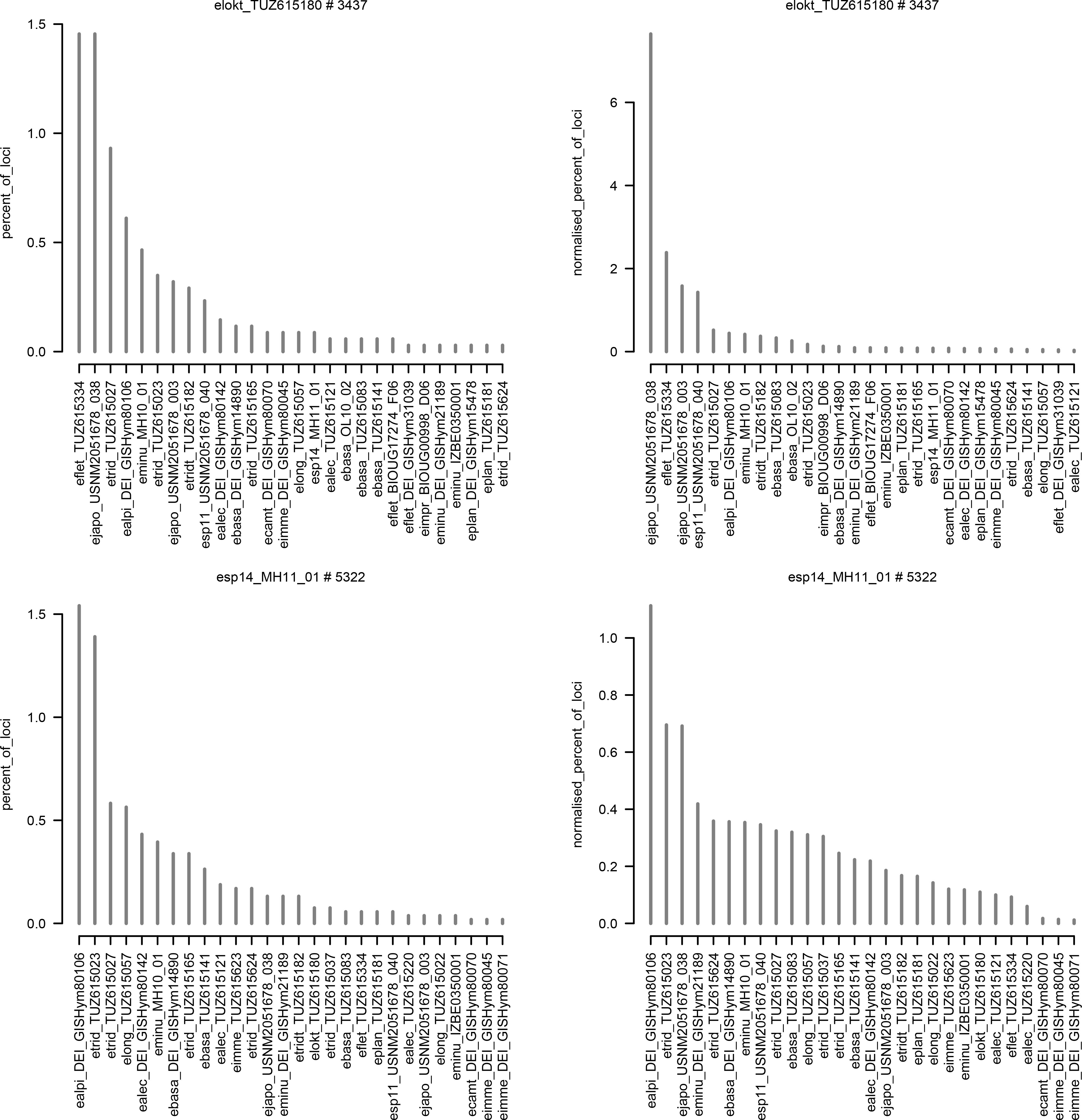

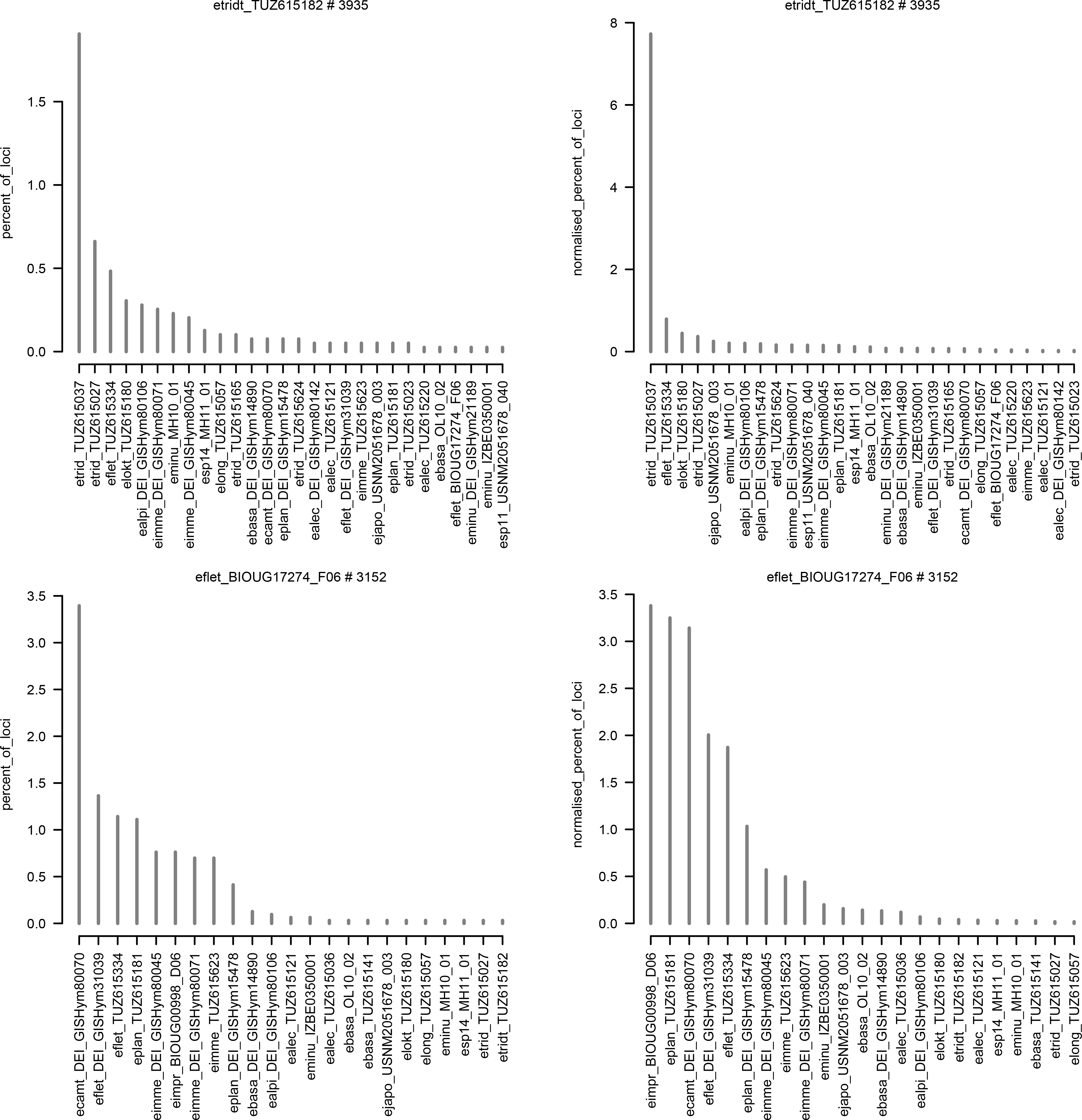

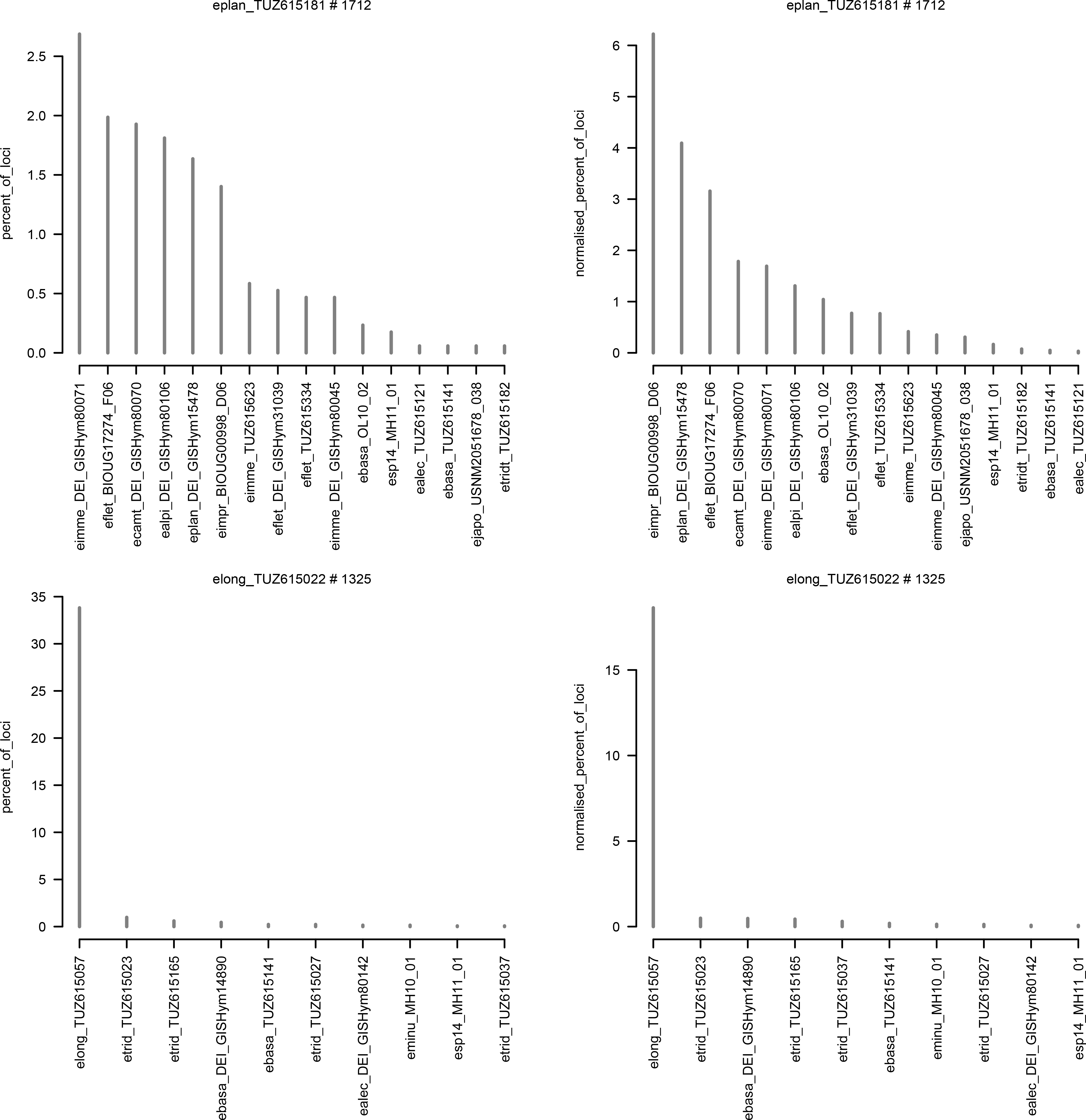

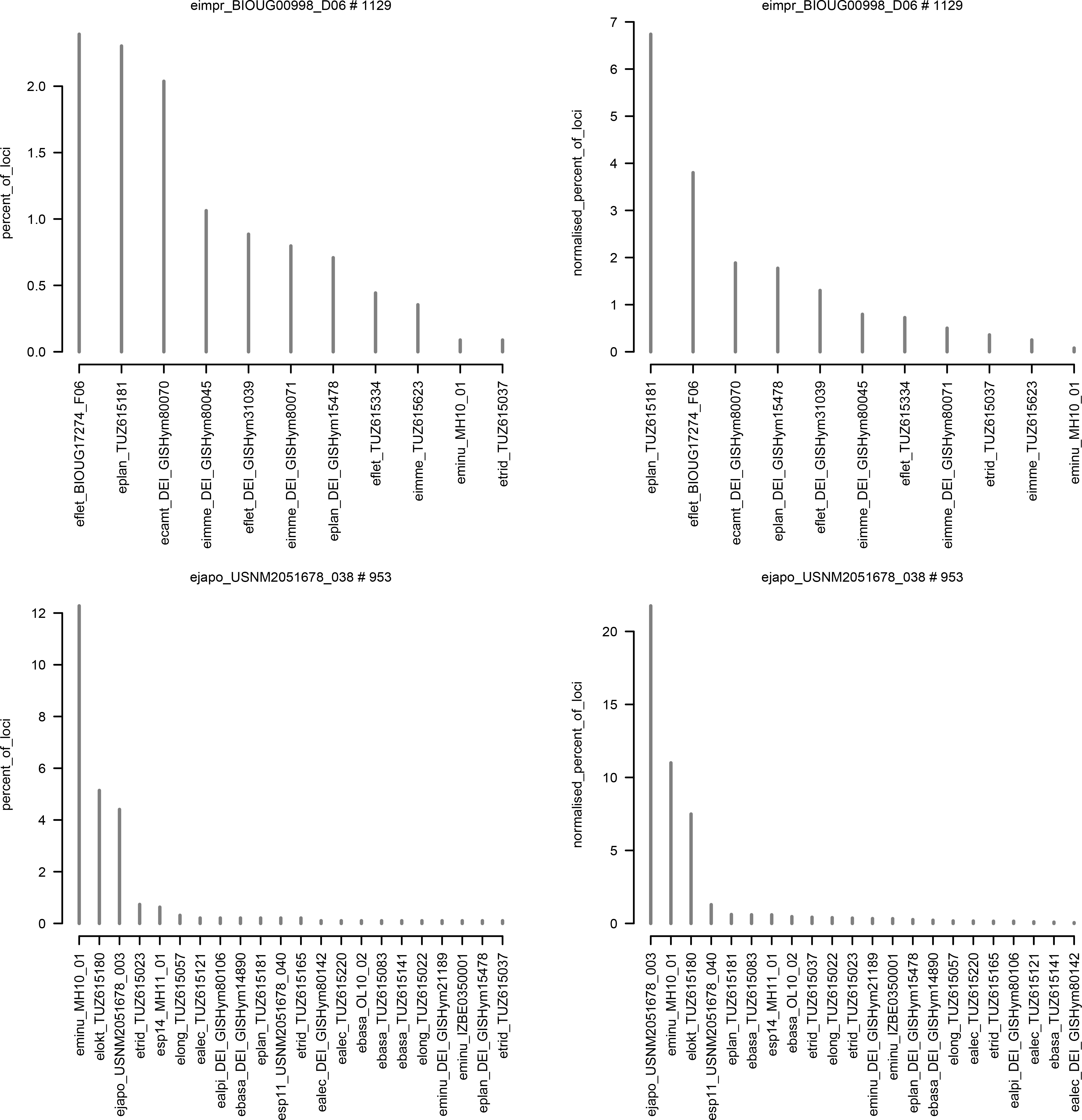

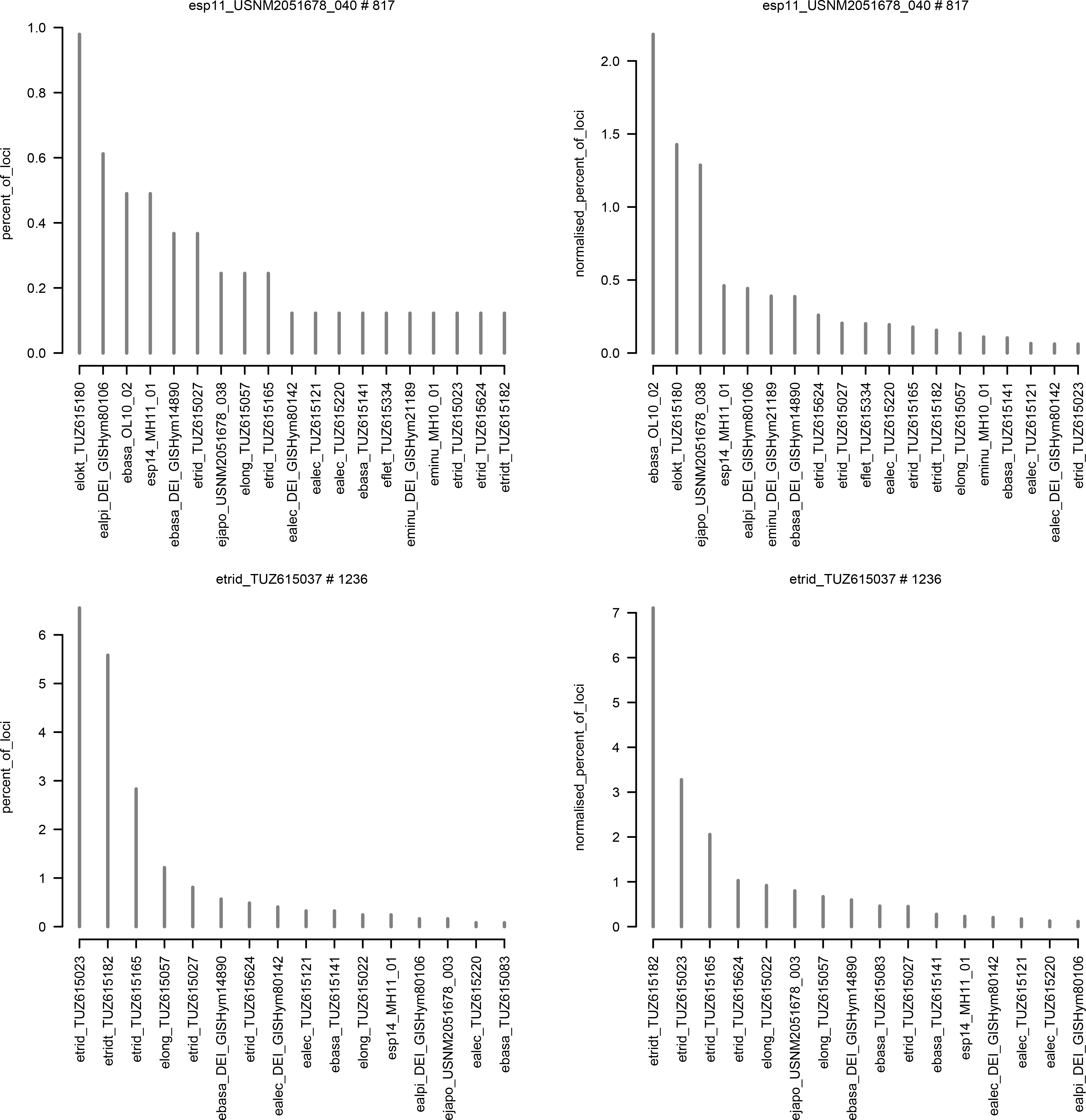

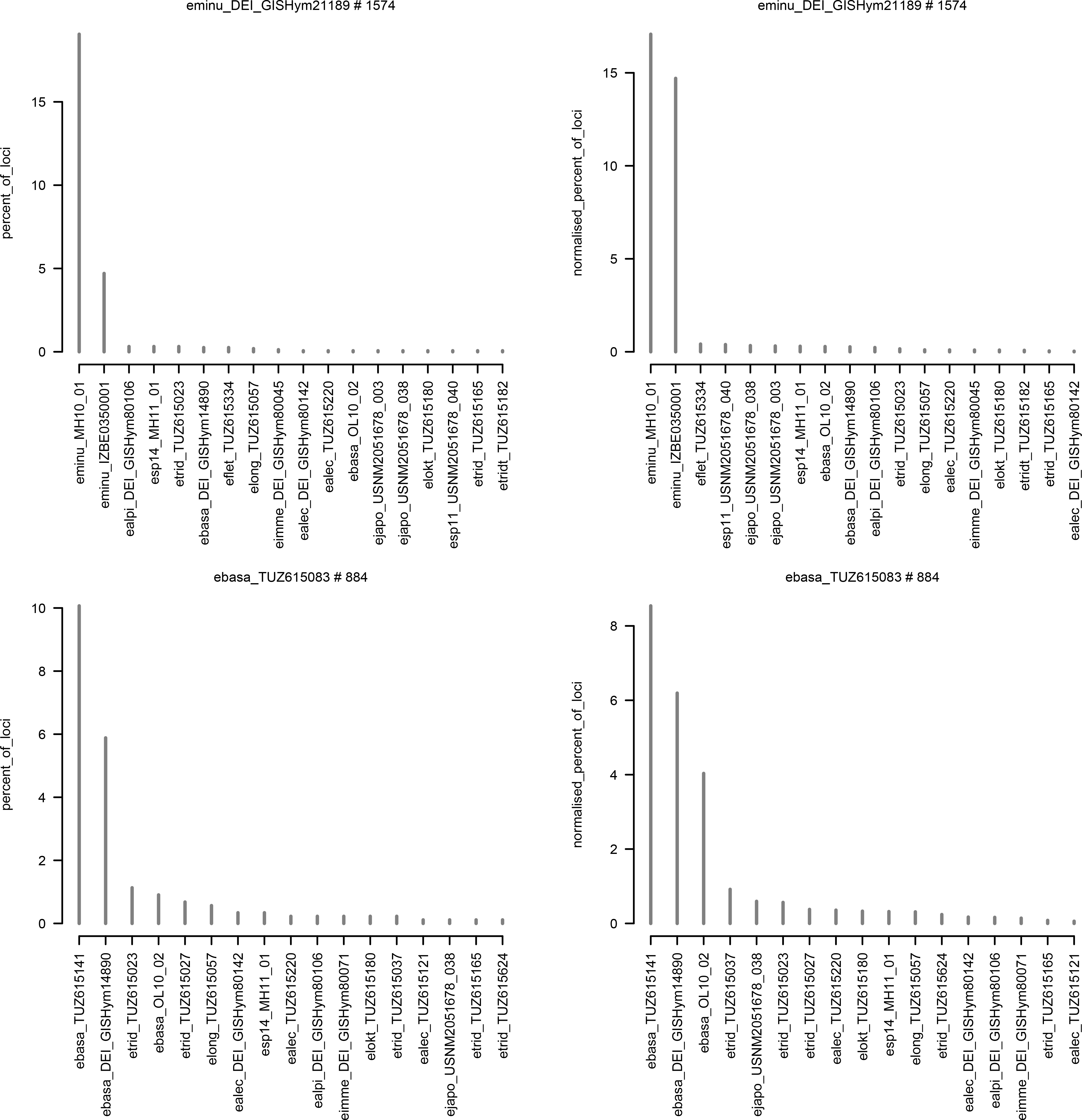

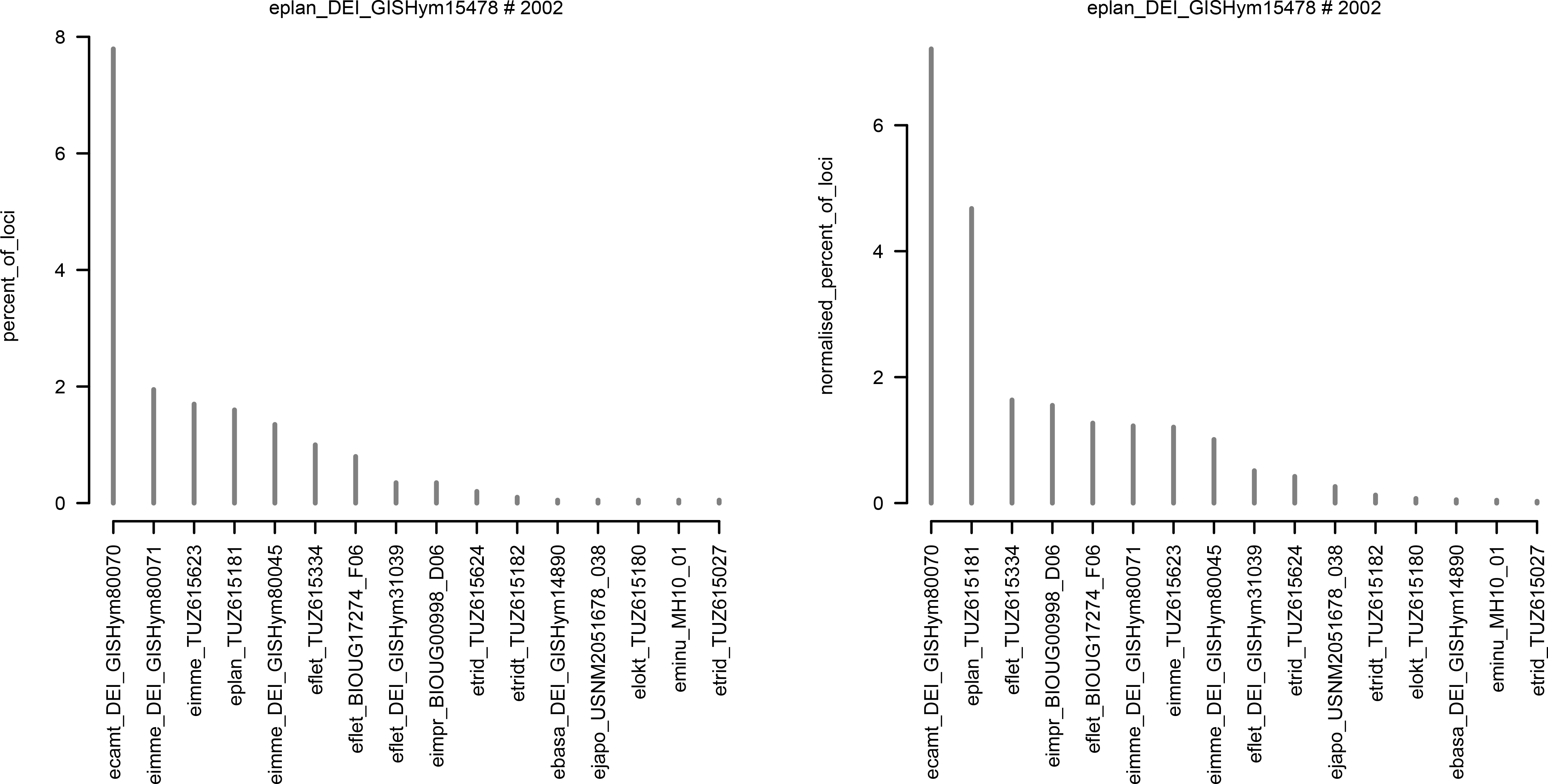

Proportion of two-fold degenerate positions calculated from initial de novo assembly (29 859 loci, clustering threshold of 80% similarity; Supplementary Data S1). f - female, m - male.

**Supplementary Data S6.**
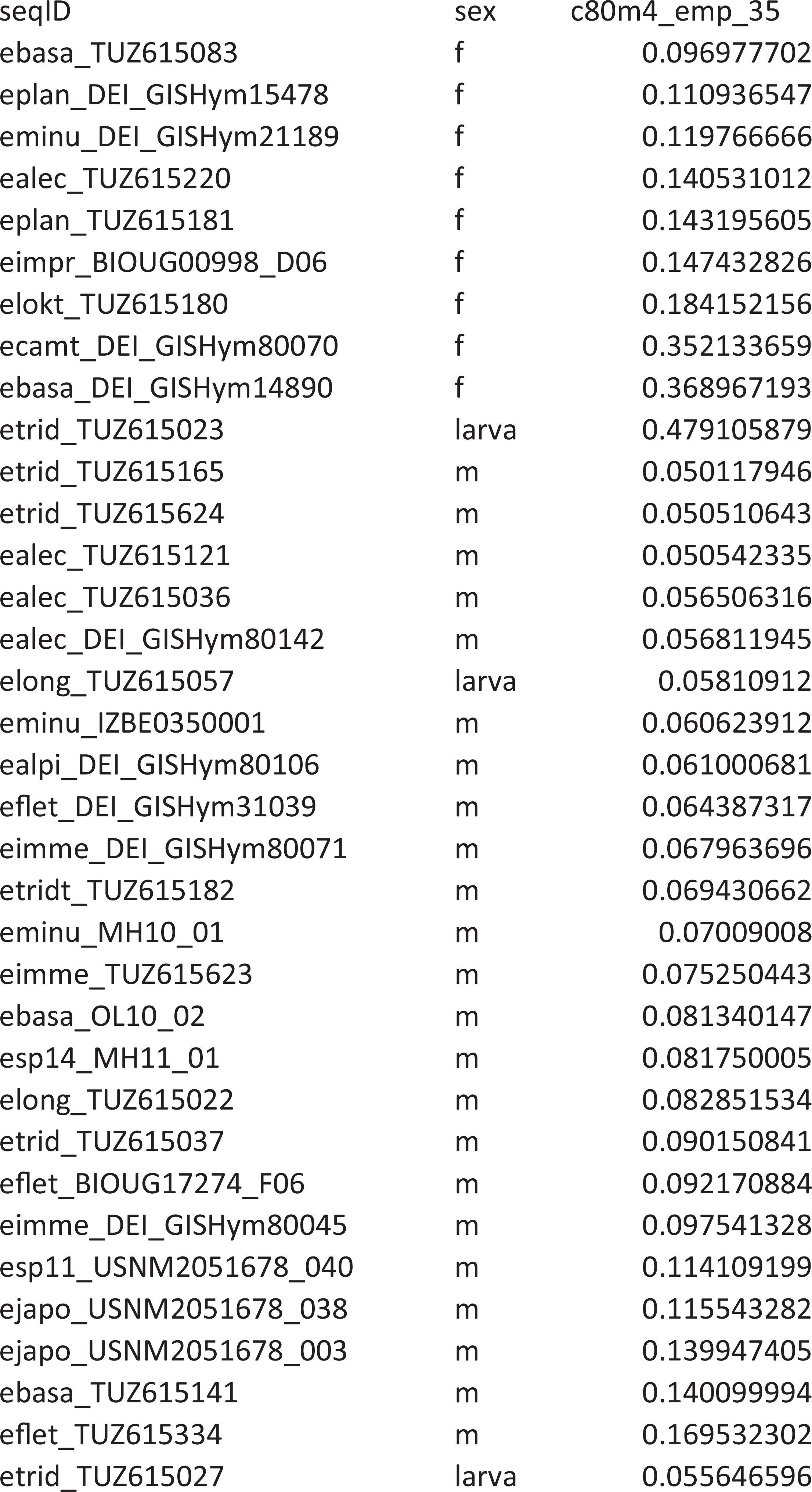

**Supplementary Data S7.**
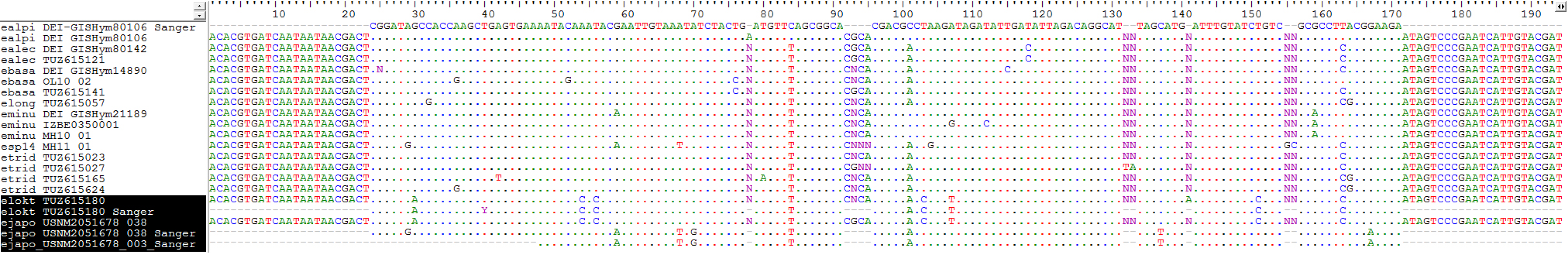

Anonymous ddRAD locus that was chosen for PCR amplification and Sanger sequencing. *Empria loktini* and *E. japonica* are highlighted. “Sanger” refers to PCR amplified Sanger sequences and the others are from *de novo* ddRAD assembly of *longicornis* group that includes only loci present at least in *E. loktini*, one *E. japonica*, and two other specimens. Note that Sanger sequence of *E. japonica* USNM2051678_038 is different from ddRAD sequence, latter of which is identical to *E. loktini* instead.

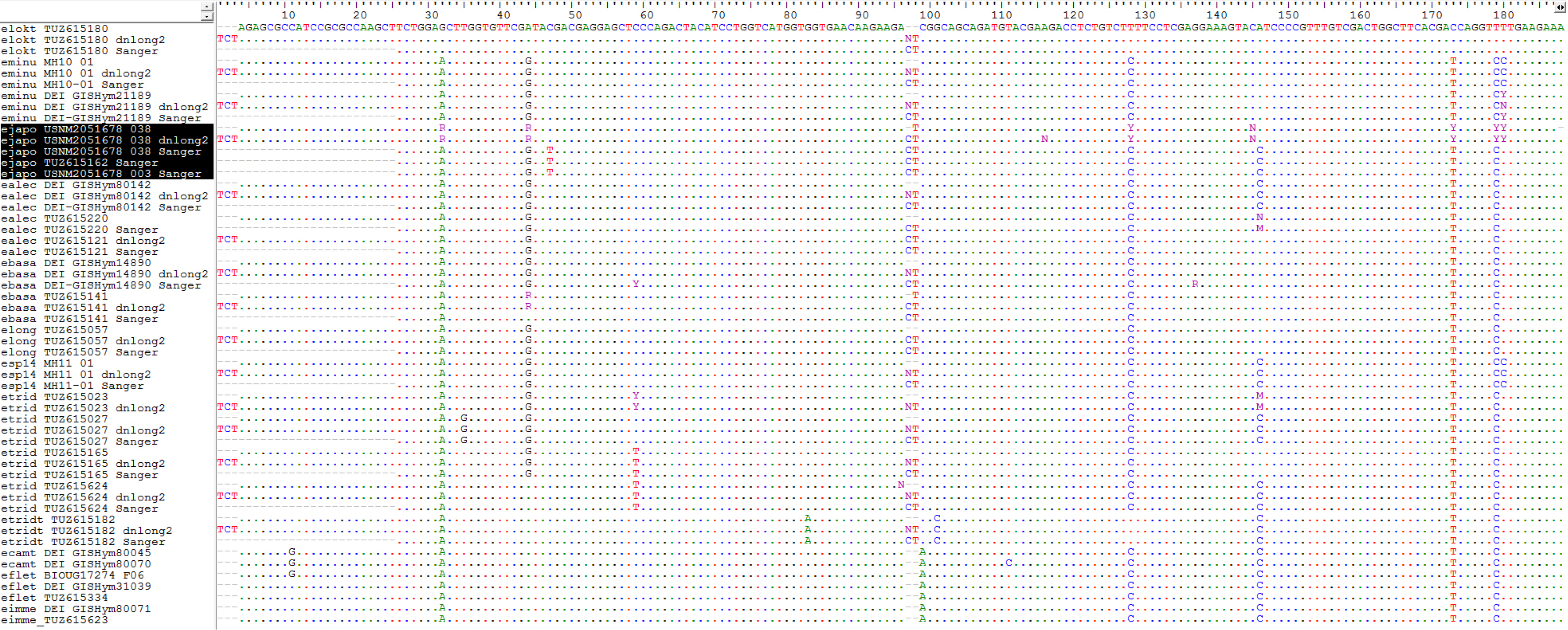

Second ddRAD locus that was chosen for PCR amplification and Sanger sequencing (ZC3H14 gene). *Empria japonica* is highlighted. “dnlong2” refers to *de novo* ddRAD assembly of *longicornis* group that includes only loci present at least in *E. loktini*, one *E. japonica*, and two other specimens, “Sanger” refers to PCR amplified Sanger sequences, and the others are from initial ddRAD assembly (*de novo*, 29 859 loci; Supplementary Data S1). Note that Sanger sequence of *E. japonica* USNM2051678_038 is different from ddRAD sequences, latter of which are identical to 9 (out of 23) other specimens. Based on two-fold degenerate positions, the ddRAD sequences of USNM2051678_038 are apparently combination of two different sequences, one of which is identical to *E. loktini* and the other to *E. minuta* MH10-01.

Specimens of *Empria* analyzed in this study and a summary of the ddRAD data in *de novo* and reference assembly. Removed individuals due to low quality of sequencing results or potential contamination issues were marked with asterisk in Sample ID.

**Supplementary Data S8.**
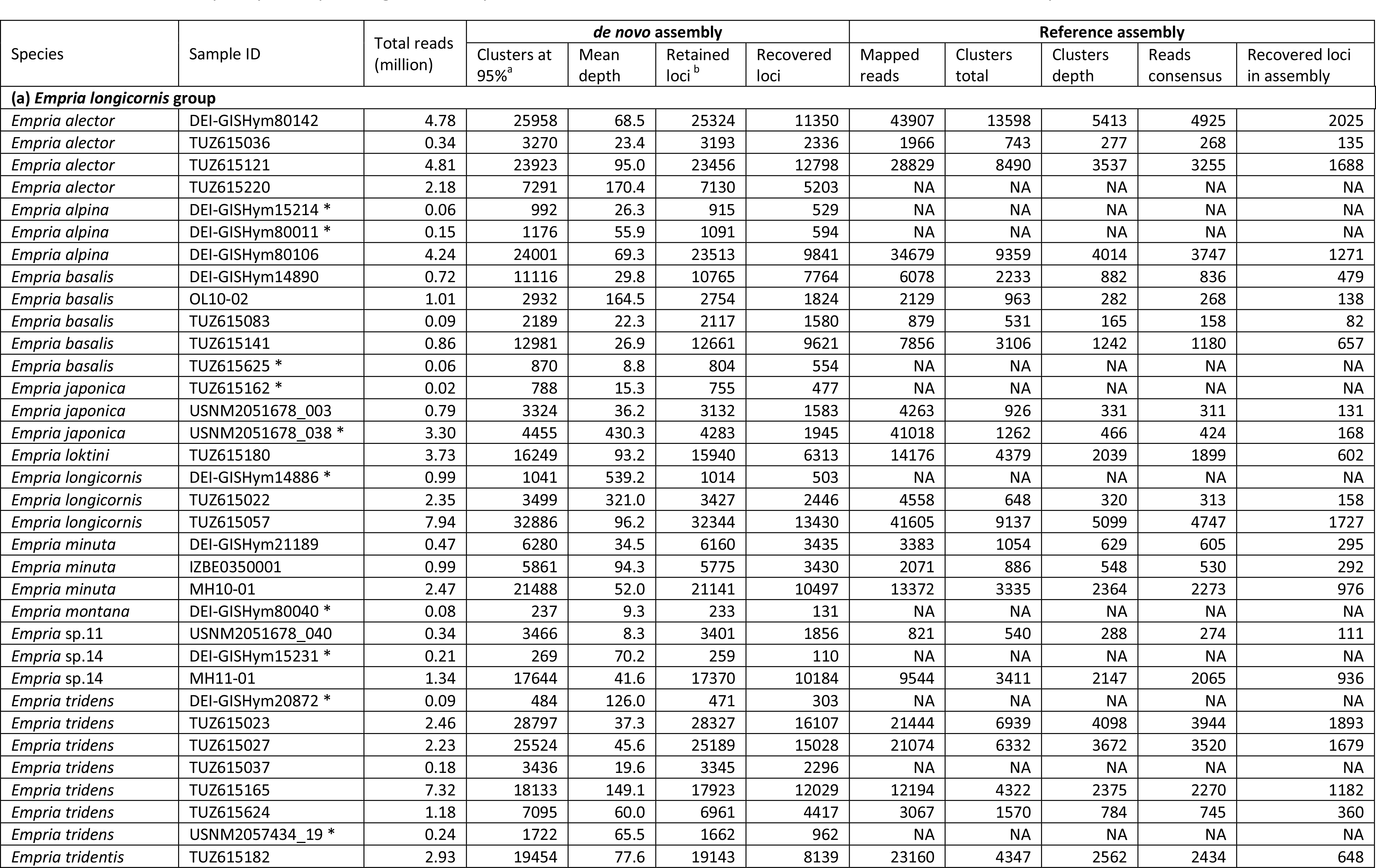

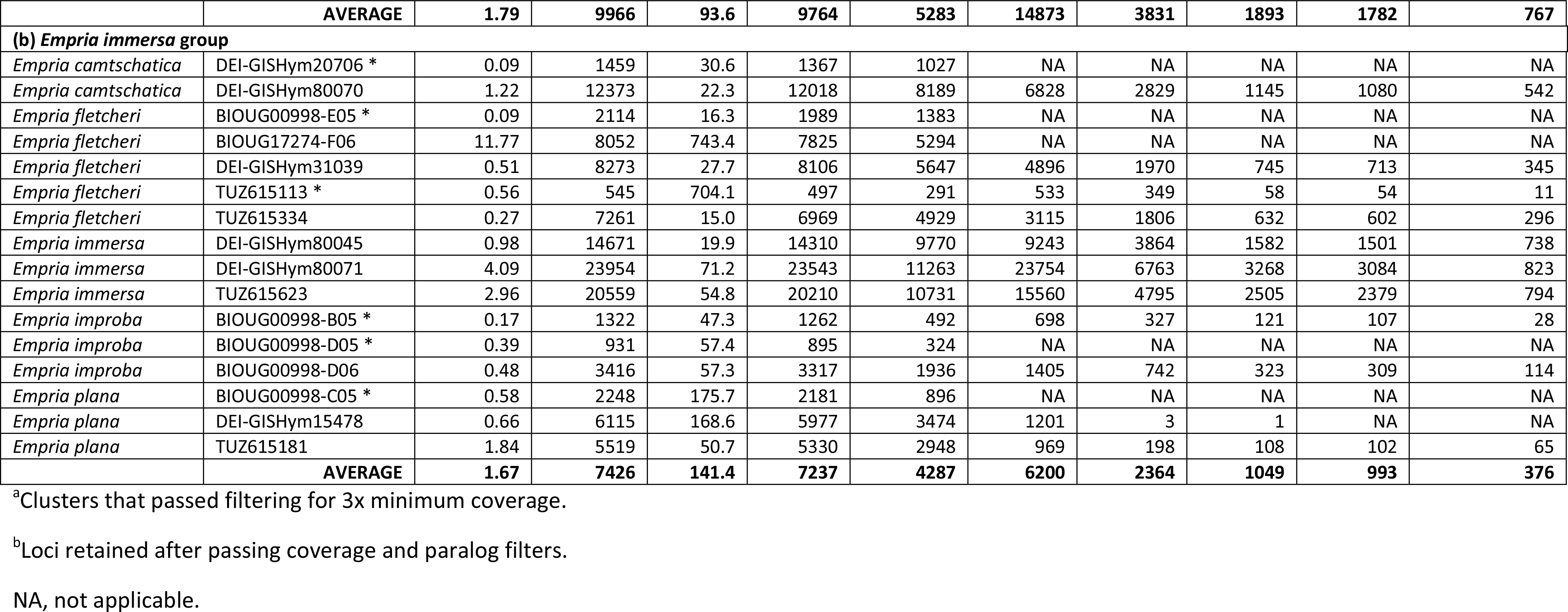

**Supplementary Data S9.**
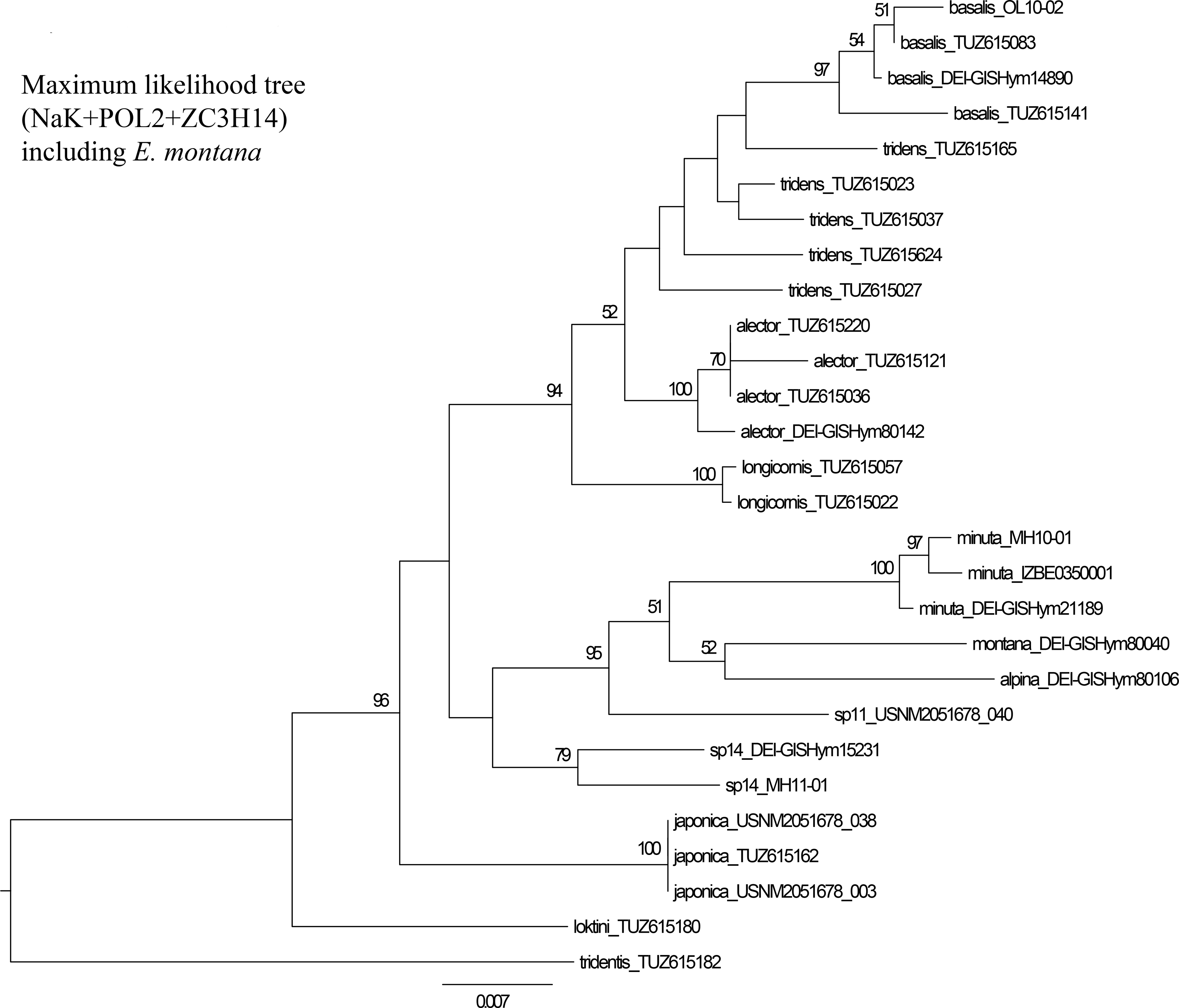

**Supplementary Data S10.**
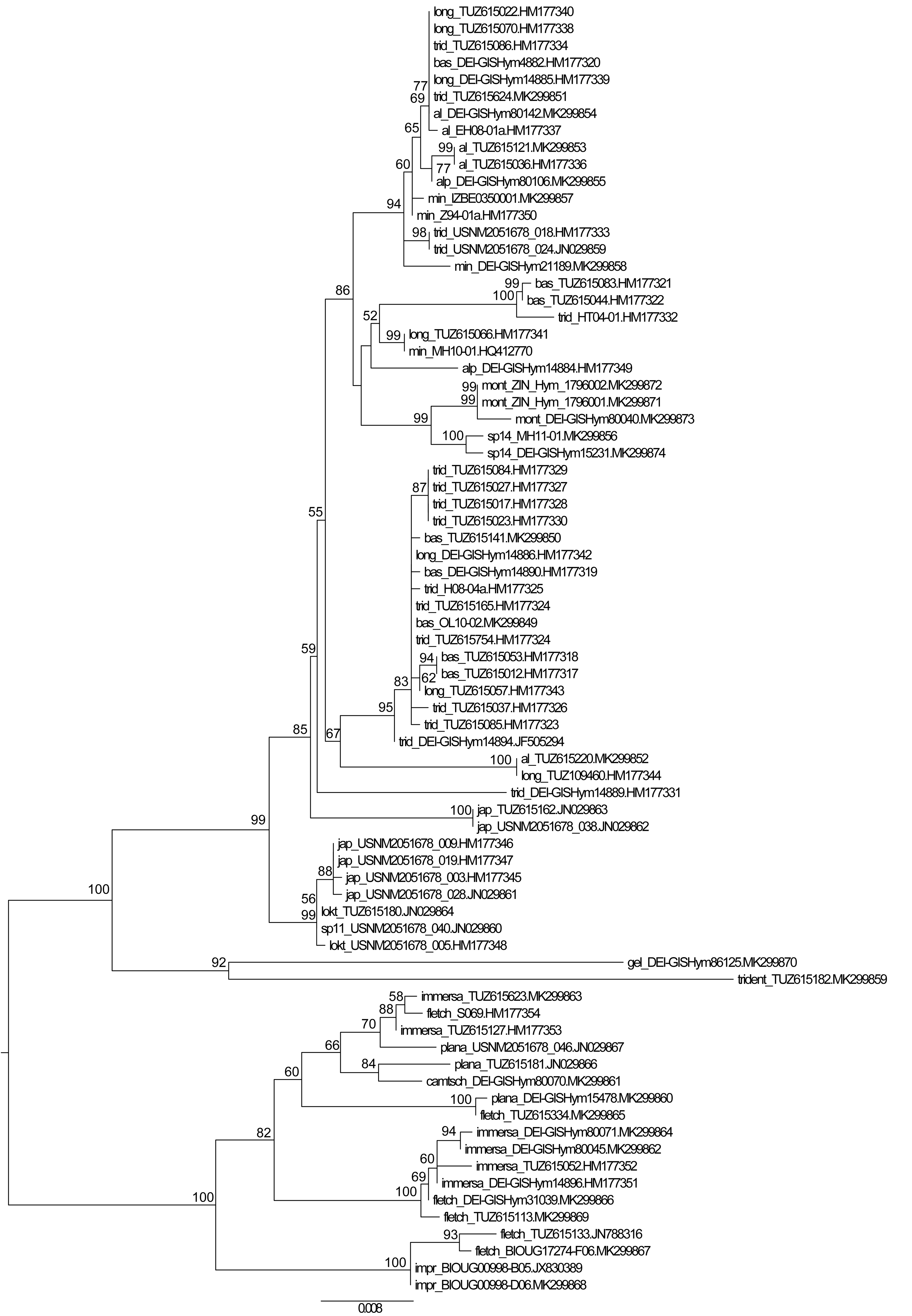

**Supplementary Data S11.**
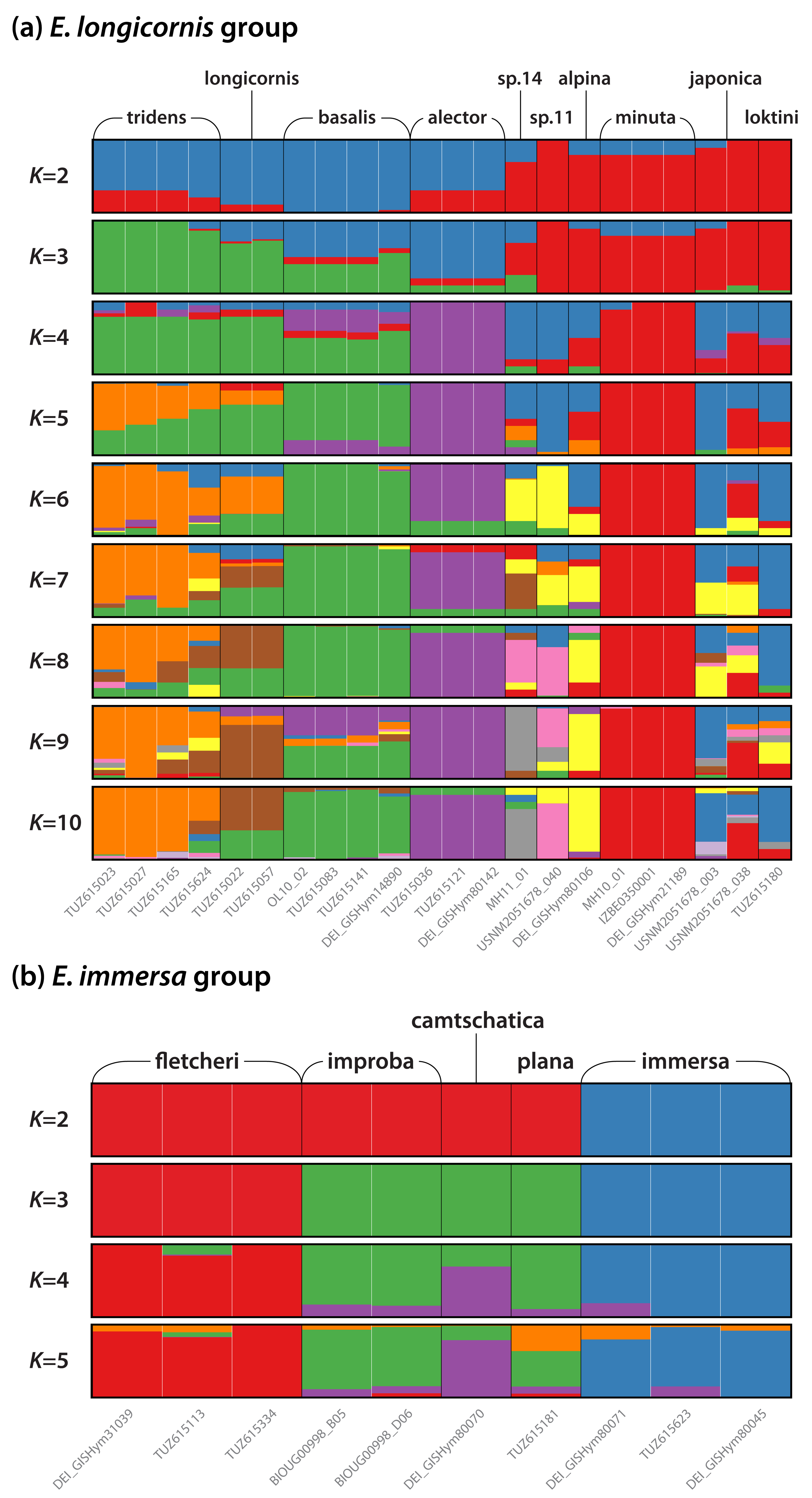

**Supplementary Data S12.**
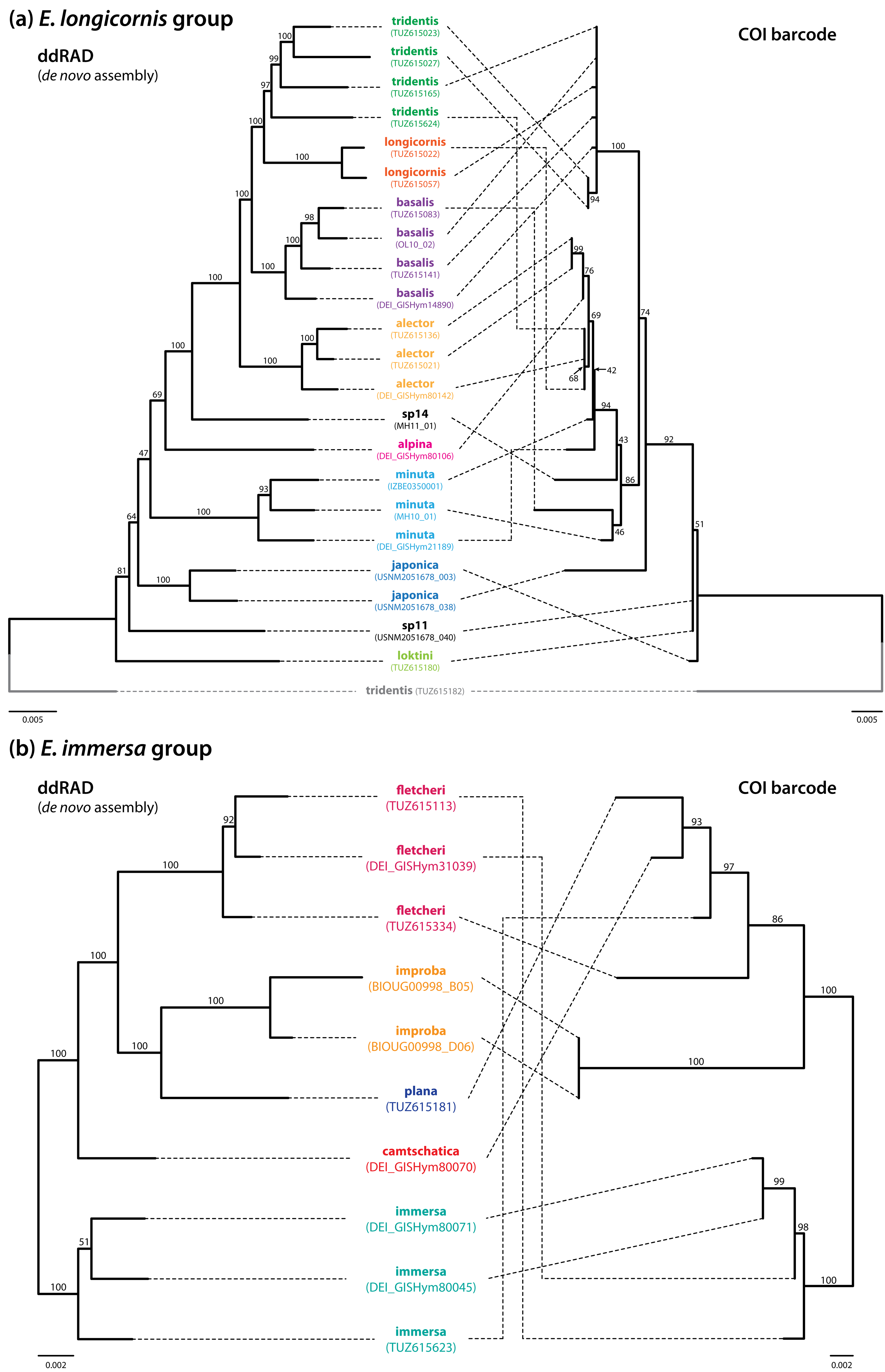

